# The limits of predicting maladaptation to future environments with genomic data

**DOI:** 10.1101/2024.01.30.577973

**Authors:** Brandon M. Lind, Katie E. Lotterhos

**Affiliations:** Department of Marine and Environmental Sciences, Northeastern University 430 Nahant Road, Nahant, MA 01908, USA

**Author notes:** **Corresponding Author** Brandon M. Lind.

**Keywords:** genomic offset, environmental change, climate change, assisted gene flow, genomic forecasting, random forests, redundancy analysis, risk of non-adaptedness

## Abstract

Anthropogenically driven changes in land use and climate patterns pose unprecedented challenges to species persistence. To understand the extent of these impacts, genomic offset methods have been used to forecast maladaptation of natural populations to future environmental change. However, while their use has become increasingly common, little is known regarding their predictive performance across a wide array of realistic and challenging scenarios. Here, we evaluate four offset methods (Gradient Forests, the Risk-Of-Non-Adaptedness, redundancy analysis, and LFMM2) using an extensive set of simulated datasets that vary demography, adaptive architecture, and the number and spatial patterns of adaptive environments. For each dataset, we train models using either *all, adaptive*, or *neutral* marker sets and evaluate performance using *in silico* common gardens by correlating known fitness with projected offset. Using over 4,850,000 of such evaluations, we find that 1) method performance is largely due to the degree of local adaptation across the metapopulation (*LA*_ΔSA_), 2) *adaptive* marker sets provide minimal performance advantages, 3) within-landscape performance is variable across gardens and declines when offset models are trained using additional non-adaptive environments, and 4) despite (1), performance declines more rapidly in novel climates for metapopulations with higher *LA*_ΔSA_ than lower *LA*_ΔSA_. We discuss the implications of these results for management, assisted gene flow, and assisted migration.

## 1 Introduction

The impacts of climate change, habitat loss, and extreme weather events pose urgent challenges to the management of species, communities, habitats, and ecosystem services (Bonan, 2008; Doney et al., 2012; Hoegh-Guldberg & Bruno, 2010). Traditional methods used to infer environmental suitability, such as reciprocal transplants and common gardens, require time and resources that may not be available or feasible for many organisms of management concern, particularly for long-lived organisms where reproductive stages occur after several decades of development. Ecological forecasting models have therefore become increasingly germane to support environmental decision making by managers across both terrestrial and marine systems.

In the context of population viability in the face of environmental change, many of these models rely on theoretical expectations that the limits of species’ distributions are primarily determined by the distribution of environmental conditions (e.g., Good 1931), and that occupancy of highly suitable habitat enables increased abundance through greater survival and reproduction (i.e., fitness) of individuals (Brown, 1984). Such methods, termed species distribution models or ecological niche models (see Elith & Leathwick, 2009 for a discussion on terminology) are correlative approaches that are often used to predict (relative) habitat suitability for a single species (Lee-Yaw et al., 2022). This information is used to understand potential impacts on the species from future climate change. However, these methods often ignore aspects of the species’ evolutionary history that could be important for predicting long-term population persistence, such as the environmental drivers of local adaptation or spatial patterns of adaptive genetic variation (Waldvogel et al., 2020).

Subsequent methods, termed genomic offsets (reviewed in Capblancq et al., 2020; Rellstab et al., 2021), have attempted to address these shortcomings by modeling relationships between environmental and genetic variation to predict maladaptation of natural populations to either future climates *in situ*, or to predict the relative suitability of these populations for the specific environment of a restoration site. Empirical attempts to confirm predictions from genomic offset models are rare and, compared to attempts *in silico* (Láruson et al., 2022), have found relatively weaker relationships between predicted maladaptation to common garden climates and the measurement of phenotypic proxies for fitness from individuals grown in these same environments (e.g., Capblancq & Forester, 2021; Fitzpatrick et al., 2021; Lind et al., 2024). Even so, these empirical results have consistently shown the expected negative relationship between predicted offset and common garden performance. Further, many of these studies found that genomic offsets often perform better than climate or geographic distance alone (e.g., Capblancq & Forester, 2021; Fitzpatrick et al., 2021; Láruson et al., 2022; Lind et al., 2024).

Across empirical and *in silico* studies, little difference in performance was found between models trained using only adaptive markers (i.e., known *in silico*, or candidates from empirical genotype-environment [GEA] associations) and those chosen at random, suggesting that genome-wide data may be sufficient to capture signals relevant to environmental adaptation.

Together, these results suggest that genomic offset methods may provide valuable insight for management. Little is known, however, about how robust these methods are across a wide array of realistic empirical scenarios, nor the extent to which independent methods will arrive at similar conclusions when analyzing the same data. Indeed, concerns regarding the accuracy of ecological forecasting models present a primary limitation towards incorporating inferences from these models into management (Clark et al., 2001; Schmolke et al., 2010) and genomic offset models are no exception. Major questions still remain about how performance is affected by aspects of the evolutionary history of sampled populations, the type of signals in putatively ideal datasets that may mislead offset inference, the importance of identifying environmental drivers of local adaptation *a priori*, and the consistency of predictive performance across contemporary environmental space. Finally, because novel climates with no recent analog are expected to increase in the future (Lotterhos et al., 2021; Mahony et al., 2017) there is also uncertainty regarding the performance of forecasting models when predictions are made to environments that drastically differ from those used to train and build the models themselves (Fitzpatrick et al., 2018; Lind et al., 2024).

While much uncertainty remains regarding the predictive performance of genomic offsets, the domain of applicability (i.e., the circumstances under which a method is acceptably accurate) for these methods can be more precisely defined using simulated data (Lotterhos et al., 2022). Simulated data, where there is no error in the estimation of allele frequencies, environmental variables, individual fitness, or the knowledge regarding the drivers of local adaptation, present ideal circumstances for understanding the limits of genomic offsets and the circumstances under which data from naturally occurring taxa will provide useful inference. To provide relevant inference regarding the domain of applicability, simulations should capture the complexities of empirical data with biological realism (e.g., clinal or patchy environments), present contrasting cases of differing scenarios while controlling for important features of the data (e.g., varying population connectivity but controlling for mean differentiation), and challenge methods using adversarial scenarios that capture extreme characteristics of empirical data (e.g., prediction to novel environments with no current analog available for model training; Lotterhos et al. 2022).

Here, we use a wide array of previously published biologically realistic, contrasting, and adversarial simulations from Lotterhos (2023) in an attempt to more precisely define the limits of predictive performance of five genomic offset methods (Table 1): Gradient Forests (GF_offset_; *sensu* Fitzpatrick & Keller, 2015), the Risk Of Non-Adaptedness (RONA, Rellstab et al., 2016), Latent Factor Mixed Models (LFMM2_offset_, *sensu* _Gain_ & _François, 2021_, and redundancy analysis (RDA_offset_, *sensu* Capblancq & Forester, 2021). The main goal of this study was to understand how the evolutionary and experimental parameters used in the training and evaluation of offset methods affect the accuracy of the methods’ projections of maladaptation under ideal empirical scenarios (i.e., using data with no inherent error). Using these scenarios, we ask the following six questions: 1) Which aspects of the past evolutionary history affect performance of offset methods? 2) How is offset performance affected by the proportion of loci with clinal alleles in the data? 3) Is method performance driven by causal loci or by genome-wide patterns of isolation-by-environment? 4) What is the variation of model performance across the landscape? 5) How does the addition of non-adaptive nuisance environments in training affect performance? 6) How well do offset models extrapolate to novel environments outside the range of environmental values used in training?

**Table 1.** Genomic offset methods used for evaluation. Genomic offset methods differ in their capability to use multivariate environmental data in training as well as whether a correction for population genetic structure is applied. Superscripts apply to the following reference citations: 1 - Fitzpatrick & Keller, 2015; 2 - Capblancq & Forester, 2021; 3 - Gain & François, 2021; 4 - Rellstab et al., 2016.

| Method | abbr. | Multivariate? | Structure correction? |
| --- | --- | --- | --- |
| Gradient Forests<sup>1</sup> | GF<sub>offset</sub> | Yes | No |
| Redundancy Analysis<sup>2</sup> with population structure correction | RDA-corrected | Yes | Yes, with axes loadings from PCA* |
| Redundancy Analysis<sup>2</sup> without population structure correction | RDA-uncorrected | Yes | No |
| Latent factor mixed model from Landscape and Ecological Association Studies R package<sup>3</sup> | LFMM2<sub>offset</sub> | Yes | Yes, with latent factors |
| Risk Of Non-Adaptedness<sup>4</sup> | RONA | No | No |

\* principal component analysis

## 2 Methods

Throughout this manuscript we will be citing code used to carry out specific analyses in-line with the text. Supplemental Notes S1-S2 outlines and describes the sets of scripts or, most often, jupyter notebooks, used to code analyses. Scripts and notebooks are both referenced as Supplemental Code (SC) using a directory numbering system (e.g., SC 02.05). More information regarding the numbering system, archiving, and software versions can be found in the Data Availability section.

### 2.1 Explanation of Simulations and Training Data

To train offset methods we used single nucleotide polymorphism (SNP) and environmental data from a set of previously published simulations (225 levels with 10 replicates each) of a Wright-Fisher metapopulation of 100 demes on a 10 x 10 grid evolving across a heterogeneous landscape (Lotterhos, 2023). Each dataset was simulated under a combination of the following four evolutionary parameters: i) three landscapes (10 populations x 10 populations) that varied in vicariance and environmental gradients (*Estuary - Clines*; *Stepping Stone - Clines*; and *Stepping Stone - Mountain*), ii) five demographies that varied population size and migration rates across the landscape, iii) three genic levels that varied in the effect size and number of mutations underlying adaptation (mono-, oligo-, and polygenic), and iv) five pleiotropy levels that varied the number of quantitative traits under locally stabilizing selection (*n*_traits_ ∈ {1, 2}), presence of pleiotropy (when *n*_traits_ = 2), and variability of selection strength across individual traits (see Fig. 1 in Lotterhos 2023).

**Figure 1.**
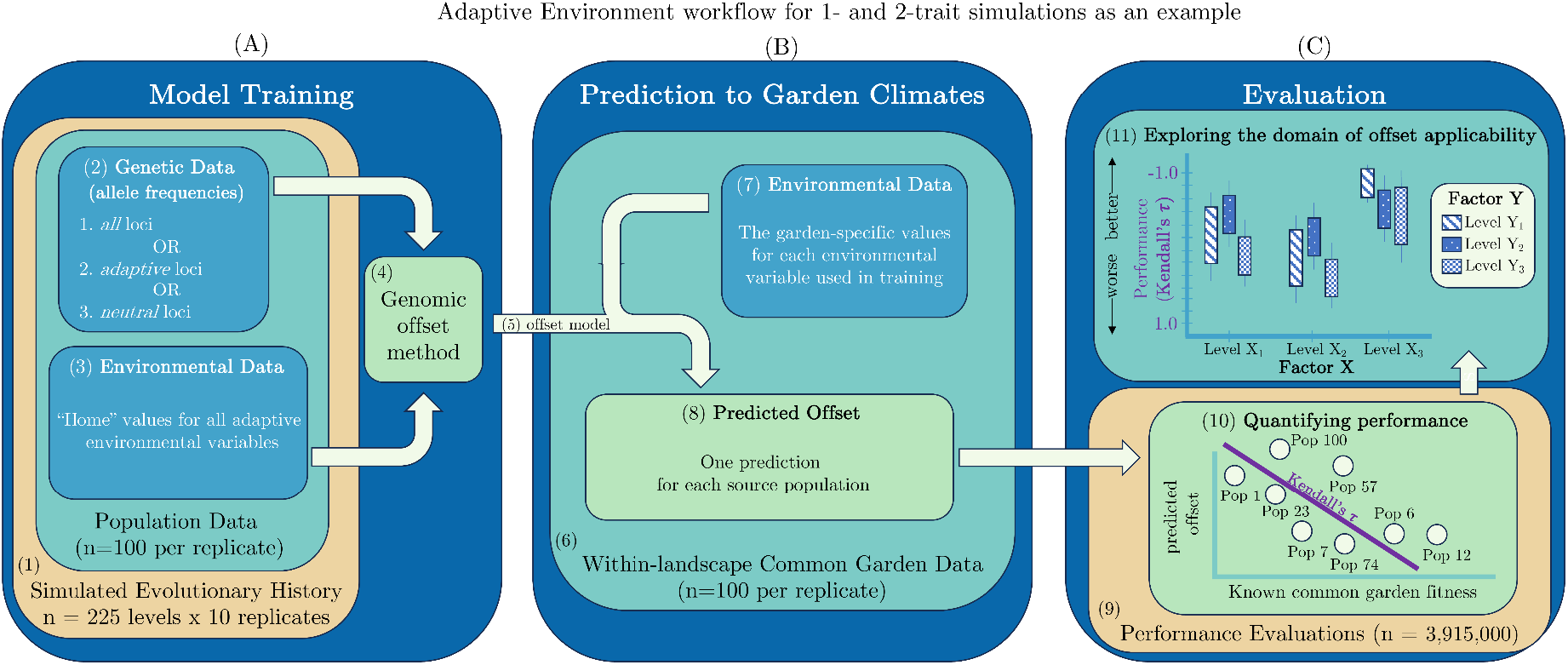
Analysis of 1-, 2-, and 6-trait simulations included three main phases: A) model training, B) model prediction, and C) evaluation of models. The *Adaptive Environment* workflow is shown as an example of the processing of 1- and 2-trait simulation data for genomic offset evaluation. In total, three general workflows are used to evaluate genomic offset methods (Table 2). Subpanels of this schematic are numbered for referencing in Table 2 and the main text.

The adaptive trait(s) were under selection by a different environmental variable, where the optimum trait value was given by the local environment on the landscape. The adaptive trait(s) undergoing selection responded to either a latitudinal temperature gradient (*temp*; *n*_traits_ = 1), or to both *temp* and a longitudinal “*Env2*” gradient (*n*_traits_ = 2). *Env2* represented distinct biological analogies depending on the context: in the *Stepping Stone - Mountain* landscape *Env2* was analogous to elevation (e.g., as with tree species), whereas in the *Estuary - Clines* landscape the *Env2* environment was analogous to gradients of salinity within coastal inlets connected only by the outer marine (ocean) environment (e.g., as with stickleback or oyster species).

**Table 2.** Workflows used to process simulation data for the evaluation of genomic offset methods. Numbers given in column names refer to locations in schematic of Fig. 1. The *Adaptive Environment* workflow processes all population data from 1- and 2-trait (example shown in Fig. 1) as well as 6-trait simulations using only adaptive environmental variables in training, and evaluates performance in each garden on the metapopulation landscape. The *Nuisance Environment* workflow processes 1-, 2-, and 6-trait simulations similarly to the *Adaptive Environment* workflow, except in addition to adaptive environmental variables used in training, non-adaptive (i.e., nuisance) environmental variables are also used - each bracketed set of environmental variables indicate a distinct nuisance level (e.g., “*1-trait 1-nuisance*” = [AE_1-trait_ environments + Env2] and “*1-trait 4-nuisance*” = [AE_1-trait_ environments + Env2 + ISO + PSsd + TSsd]). The *Climate Novelty* workflow uses trained models from the *Adaptive Environment* workflow (Fig. 1A-5) and evaluates offset in 11 novel environments relative to the range of environments used in training. See Supplemental Note S3 for details regarding the choice of *Climate Novelty* environmental values and visualizations of climate data in principal component space. See Supplemental Notes S1-S2 for descriptions of coding workflows. Counts of evaluations were tabulated in SC 02.10.01.

| Workflow | $n_{\text{traits}}$ | (1) Simulations Levels<br>(replicates per level) | (3, 7) Environmental Data | Training and<br>Prediction? | (6) Within-landscape<br>Evaluation? | (9) Total<br>Performance<br>Evaluations |
| --- | --- | --- | --- | --- | --- | --- |
| Adaptive<br>Environment<br>(AE) | 1-trait | 45 (10) | temp | Yes | Yes | 675,000 |
|  | 2-trait | 180 (10) | temp + Env2 |  |  | 3,240,000 |
|  | 6-trait | 1 (1) | MAT + MTwetQ + MTDQ + PDM + PwarmQ + PWM |  |  | 3,000 |
| Nuisance<br>Environment<br>(NE)** | 1-trait | 45 (1) | [AE<sub>1-trait</sub> environments + Env2] +/- [ISO + PSsd] +/- [TSsd] | Yes | Yes | 175,500 |
|  | 2-trait | 180 (1) | [AE<sub>2-trait</sub> environments + ISO + PSsd] +/- [TSsd] |  |  | 432,000* |
|  | 6-trait | 1 (1) | [AE<sub>6-trait</sub> environments + ISO + PSsd + TSsd] |  |  | 1,200* |
| Climate<br>Novelty<br>(CN)<sup>†</sup> | 1-trait | 45 (10) | AE<sub>1-trait</sub> environments | Prediction only<br>(using AE<br>training models) | No | 64,800* |
|  | 2-trait | 180 (10) | AE<sub>2-trait</sub> environments |  |  | 259,200* |
|  | 6-trait | 1 (1) | AE<sub>6-trait</sub> environments |  |  | 144* |

\* excludes RONA
\*\* The set of population values for each unique nuisance environment were the same across traits and landscapes
<sup>†</sup> includes evaluation of climate center and eleven *Climate Novelty* scenarios

Twenty independent linkage groups were simulated. Of these, mutations that had effects on one or more phenotypes under selection (i.e., quantitative trait nucleotides, QTNs) were allowed to evolve on only ten linkage groups, and neutral mutations were added to all 20 linkage groups with tree sequencing (for details see Lotterhos 2023). Adaptive traits were determined additively by effects of QTNs.

In all simulations, phenotypic clines evolved between each trait and the selective environment (Lotterhos, 2023), where populations became locally adapted to their environment, measured at the metapopulation level as the mean difference of demes in sympatry minus allopatry (*LA*_ΔSA_, Blanquart et al., 2013). *LA*_ΔSA_ equates to the average levels of local adaptation at the deme level which can be calculated for each deme by both home-away (*LA*_Δ_HA) and local-foreign (*LA*_ΔLF_) measures.

These simulations represent a wide array of realistic, contrasting, and adversarial scenarios in which we could more precisely define the domain of applicability of offset methods. For instance, in the *Stepping Stone - Mountain* landscape, geographic distance and environmental distance were not strongly correlated, whereas in the *Stepping Stone - Clines* and *Estuary - Clines* they were. Additionally, the proportion of mutations with monotonic frequency gradients (i.e., allelic clines) underlying local adaptation varied across the simulated datasets (Lotterhos, 2023), which may also affect offset performance. These simulations also presented demographic scenarios in which selection was confounded with genetic drift or population genetic structure. For each simulation, ten individuals were randomly chosen per population for a total of 1000 individuals. Individual genotypes were coded as counts of the derived allele. Alleles with global minor allele frequency (MAF) < 0.01 were removed. Using all 100 populations, population-level derived allele frequencies and current environmental values were used as input to train offset methods.

In addition to the 2250 simulated Wright-Fisher datasets (225 levels * 10 replicates), we also included a non-Wright-Fisher case with range expansion from three refugia and secondary contact (Lotterhos 2023). This simulation evolved variable degrees of admixture across the landscape. Six moderately polygenic environmental traits (*n*_traits_ = 6) were under selection from the environment. Environments were based on six weakly correlated environmental variables taken from Bioclim environmental measures of western Canada. The simulation evolved local adaptation at all six traits with unconstrained pleiotropy. For more details on simulations, see (Lotterhos, 2023).

### 2.2 Evaluation of Offset Methods

We investigated the performance of five implementations of four genomic offset methods (Table 1): GF_offset_, RDA_offset_,, LFMM2_offset_, and RONA. While GF_offset_, RDA_offset_, and LFMM2_offset_ can use multivariate environmental data to train models, RONA can only account for a single environment at one time (Table 1). Additionally, while GF_offset_ and RONA do not apply correction for population genetic structure, LFMM2_offset_ does by default, and structure correction with RDA_offset_ is optional. We thus evaluate RDA_offset_ with (RDA-corrected) and without (RDA-uncorrected) population genetic structure correction (Table 1). For additional specifics related to the implementation of each offset method, see Supplemental Note S1.1-S1.4 and Fig. S1, Fig. S2, Fig. S3.

We varied construction of genomic offset training datasets for each replicate of the 1-, 2-, and 6-trait simulations by varying the marker set used in model training (Fig. 1A, Table 2; see *Q3* below). Each model was trained using genetic and environmental data from all 100 populations. The environmental variables used were only those imposing selection pressure. We predict offset from each model for each population to all 100 within-landscape common gardens from a full factorial *in silico* reciprocal transplant design (Fig. 1B). For each common garden, we quantified offset model performance as the rank correlation (Kendall’s **τ**) between the population mean fitness (averaged over sampled individuals, Equation 3 in Lotterhos 2023) and projected population offset (Fig. 1C). Strong negative relationships between fitness and predicted offset indicate higher performance of the method (note y-axes of Kendall’s τ are inverted within figures to show more intuitive performance relationships, Fig. 1C-11). We refer to the preceding processing of data as the *Adaptive Environment* workflow (Fig. 1, Table 2).

To explore the impact of the choice of environmental variables used (see *Q5* below), we used a workflow similar to the *Adaptive Environment* workflow, except instead of using only adaptive environmental variables, we used additional non-adaptive (i.e., nuisance) environmental variables in training and prediction (second row, Table 2). These nuisance variables had relatively weak correlation structure with adaptive environments and each other (Fig. S4). We refer to each of these nuisance levels by the number of traits under selection and the number of nuisance environments used (e.g., *1-trait 3-nuisance*). We refer to this workflow as the *Nuisance Environment* workflow.

Finally, to contrast with within-landscape evaluations, we explored predictive performance of *Adaptive Environment* offset models in novel environments that are beyond the range of values of those used in training (see *Q6* below). In these novelty cases, we use 11 common gardens, each progressively more distant from the average environment used in training (i.e., climate center) and evaluate performance in each garden. We refer to this workflow as the *Climate Novelty* workflow. See Supplemental Note S3 and Fig. S5 for details regarding the choice of environmental values for novelty scenarios.

### 2.3 Study Questions

#### Q1 - Which aspects of the past evolutionary history affect within-landscape performance of offset methods?

For each offset method, we used a fixed-effects type II ANOVA model to test for significant differences in the performance from 2-trait *Adaptive Environment* models trained using *all* markers using the following factors: landscape (*Estuary - Clines, Stepping Stone - Clines, Stepping Stone - Mountain*), demography (five levels describing population size and migration patterns across the landscape), genic level of architecture (three levels from oligogenic to polygenic), presence or absence of pleiotropy, proportion of loci with clinal allele frequencies (as defined in Lotterhos, 2023), degree of local adaptation (ΔSA), and common garden ID. Specifically,

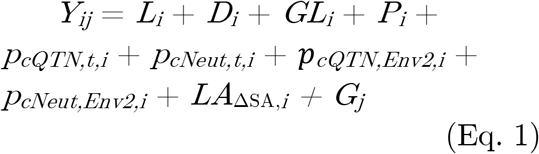

where *Y*_*ij*_ is the within-landscape performance (Kendall’s **τ**) of a single method for garden *j* in simulation *i*, with factors for landscape (*L*), demography (*D*), genic level (*GL*), presence of pleiotropy (*P*), proportion of QTN or neutral alleles with *temp* clines (respectively *p*_*cQTN,t,i*_ and *p*_*cNeut,t,i*_), proportion of QTN or neutral alleles with *Env2* clines (respectively *p*_*cQTN,Env2,i*_ and *p*_*cNeut,Env2,i*_), degree of local adaptation (*LA*_ΔSA_), and garden ID (*G*). The 473 first four factors are illustrated in Fig. 1 of Lotterhos (2023).

#### Q2 - How is offset performance affected by the proportion of clinal alleles in the data?

Clinal alleles (i.e., alleles with monotonic gradients in frequency across space) that covary with environmental clines could be weighted more heavily in offset models that emphasize loci whose allele frequencies explain significant variation across local environmental values. Using 2-trait models trained using *all* markers from the *Adaptive Environment* workflow, we used an ANOVA model (Eq. 2) to test the hypothesis that clinal alleles differentially impact model 488 performance, independent from the other factors from Eq. 1:

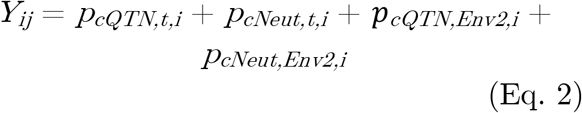

The factors representing clinal alleles in Eq. 2 are the same as those in Eq. 1.

#### Q3 - Is method performance driven by causal loci or by genome-wide patterns of Isolation 497 By Environment?

For each offset method and workflow, we varied the set of input markers for 1-, 2- and 6-trait simulations that were used in training to determine if performance of a method was driven by properties of the evolutionary forces shaping genotype-environment relationships: 1) *adaptive* markers (i.e., QTNs with effects on at least one trait), 2) *neutral* markers (SNPs on linkage groups without QTNs), and 507 3) *all* markers (union of *adaptive* and *neutral* markers, as well as non-QTN markers on the same linkage groups as QTNs). Only loci that passed MAF filtering were included in marker sets. If offset performance is determined solely by adaptive signals in genetic data, offsets trained using *adaptive* markers should have better performance than *all* or *neutral* markers, and *all* markers should have better performance than *neutral* markers.

If the marker set has little impact on offset performance, this could indicate that offset methods are giving weight to genome-wide signals present in the data. Previously, some (e.g., Lachmuth, Capblancq, Keller, et al., 2023; Lind et al., 2024) have postulated that this signal may be related to isolation by environment ((IBE, i.e., when genetic and environmental distances are positively correlated, independent of geographic distance; Wang & Bradburd, 2014).

If IBE is driving patterns of offset performance, we expect 1) performance to be similar between offsets estimated using *adaptive* markers and those estimated using *neutral* markers; 2) a greater proportion of variation in performance to be explained by *p*_*cNeut*_ than *p*_*cQTN*_ (from *Q2*); 3) a strong, positive relationship between performance and *LA* _ΔSA_; and 4) the difference in IBE between two marker sets to be positively correlated with the difference in performance of two models trained with those markers. We measure IBE as the rank correlation (Spearman’s $) between population pairwise *F* _ST_ (Weir & Cockerham, 1984) and Euclidean climate distance of adaptive environmental variables.

#### Q4 - What is the variation of model performance across the landscape?

Within a landscape, offset methods may not have high predictive performance at every site or every environment. Understanding variability in the predictive performance of offset models across the landscape is particularly relevant when offsets are used for restoration or assisted gene flow initiatives (i.e., ranking sources for a given site). If predictive performance is variable across the landscape, this may limit the usefulness of genomic offsets for such purposes even if model performance is validated in one common garden. Using the *Adaptive Environment* workflow, we visualized variation of 1- and 2-trait within-landscape performance with boxplots for each common garden for each method and landscape. To understand if variation in predictive performance was a function of the model quality, we investigated the relationship between a model’s performance variability (i.e., standard deviation across 100 common gardens) and the model’s median performance.

#### Q5 - How does the addition of non-adaptive 578 nuisance environments in training affect performance?

In practice, the environments imposing selection are rarely known *a priori*. Additionally, the inclusion of environmental measures that are not correlated with the main axes of selection may reduce model performance compared to models trained using only causal environments. To investigate the sensitivity of offset methods to environmental input we compared *Adaptive Environment* workflow models from 1-, 2-, and 6-trait simulations – where only the adaptive environment(s) are used in training (*0-nuisance*) – to models from the *Nuisance Environment* workflow trained with the same data but with the addition of nuisance environments (*N-nuisance*, where *N* > 0; Table 2).

We use nuisance environmental variables from Lotterhos (2023) that were real BioClim variables (*TSsd, PSsd*, and *ISO*) taken from British Columbia and Alberta, Canada, which have minimal correlation with causal environments and each other (Fig. S4). These three nuisance environments differ from previous implementations of such variables (Láurson et al. 2022) in that they are spatially autocorrelated whereas nuisance environments in Láruson et al. (2022) were not. For 1-trait scenarios, *Env2* was also used as a nuisance environmental variable.

If offset methods are unaffected by the addition of nuisance environmental variables, performance should not differ between *0-nuisance* and *N-nuisance* implementations. Finally, in empirical settings the set of adaptive environments are not known *a priori*. We also explored whether GF would rank adaptive environments higher than nuisance environments using weighted importance output from GF.

#### Q6 - How well do offset models extrapolate to 625 novel environments outside the range of environments used in training?

Even if offset methods have high within-landscape performance, this does not directly address situations where future environmental conditions are vastly different from the environmental conditions used for training (i.e., novel environments). If performance decreases with increasing environmental novelty relative to training data, this raises questions about the utility of genomic offsets for predicting 1) relative *in situ* vulnerability of populations to future climate change, and 2) the relative suitability of populations to restoration sites that differ drastically than those used in training.

To understand if offset performance degrades with environmental novelty relative to training data, we predicted offset to 10 novel environmental scenarios for the 1-, 2-, and 6-trait simulations using the *Climate Novelty* workflow (Table 2). The novel environmental scenarios were a set of common garden environments, *z*_*E*_, extending outward from the training populations and exceeding values observed on the landscape for all adaptive environmental variables (Supplemental Note S3). We represent these scenarios as standard deviations from the center of environmental values used in training: *z*_*E*_ ∈ {1.72, 2.35, 2.74, 3.13, 3.53, 656 3.92, 4.31, 4.70, 5.09, 5.48, 5.88}. Fitness in novel environments was estimated assuming that the phenotypic optimum continues to have a linear relationship with the environmental variable (Equation 3 in Lotterhos 2023).

## 3 Results

### Q1 - Which aspects of the past evolutionary history affect within-andscape performance of offset

The ANOVA model (Eq. 1) indicated that the degree of local adaptation of the metapopulation (*LA*_ΔSA_) was the primary factor influencing offset performance, followed common garden location, demography, and landscape (Table S1; Fig. S6). Within the simulations, *LA*_ΔSA_ was impacted by pleiotropy, the relative strength of selection, and landscape, (Fig. S7; see also Figs. S2A, S2B in Lotterhos, 2023), so there may be some confounding among these factors.

In line with the ANOVA model, the performance of specific offset methods generally increased with increasing *LA*_ΔSA_ (Fig. 2), but there were some interesting differences among methods. For instance, GF_offset_, LFFM2_offset_, RDA-uncorrected, and RONA_temp_ all improved as *LA*_ΔSA_ increased, while RDA-corrected and RONA_Env2_ showed relatively weaker relationships.

**Figure 2.**
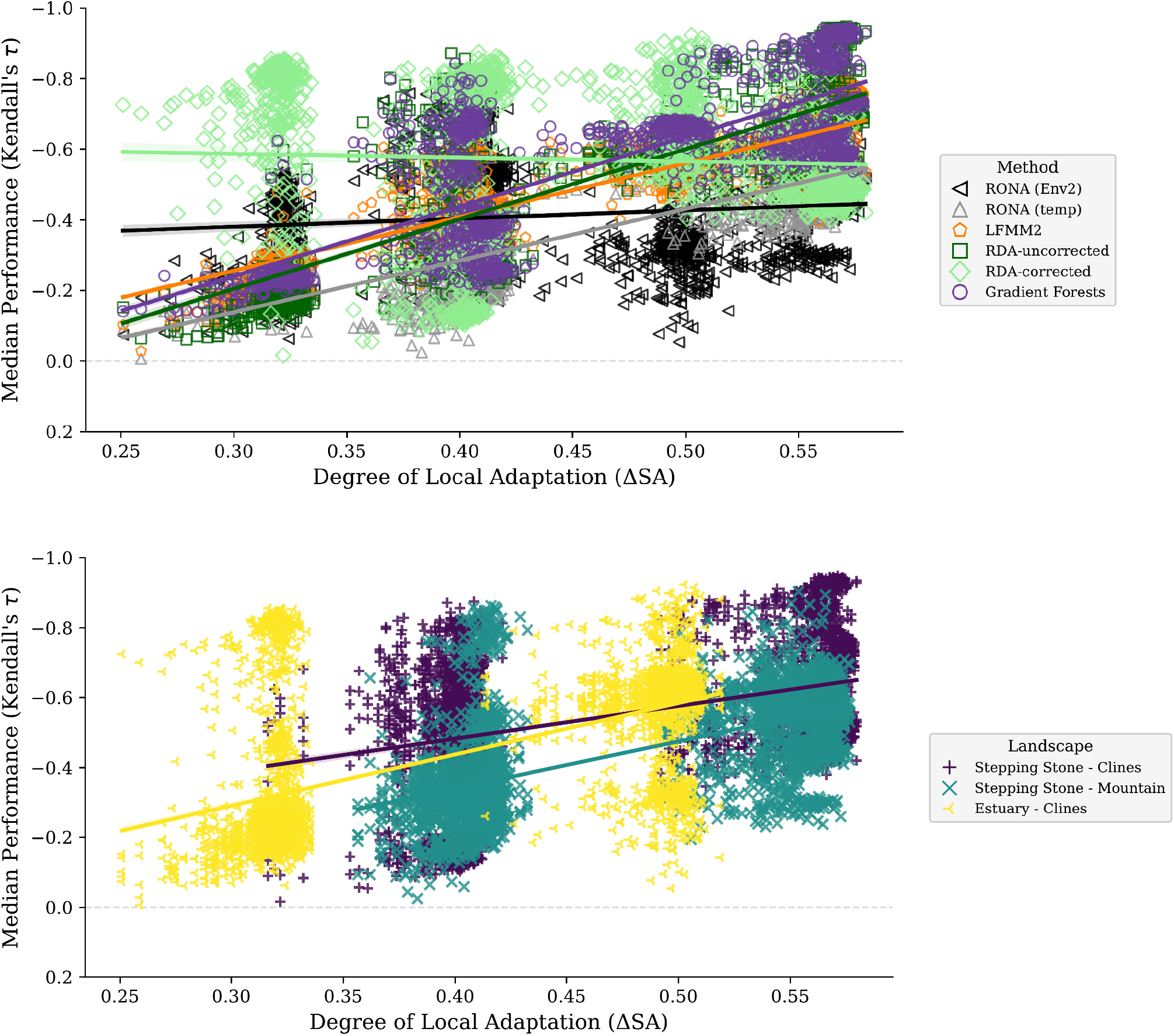
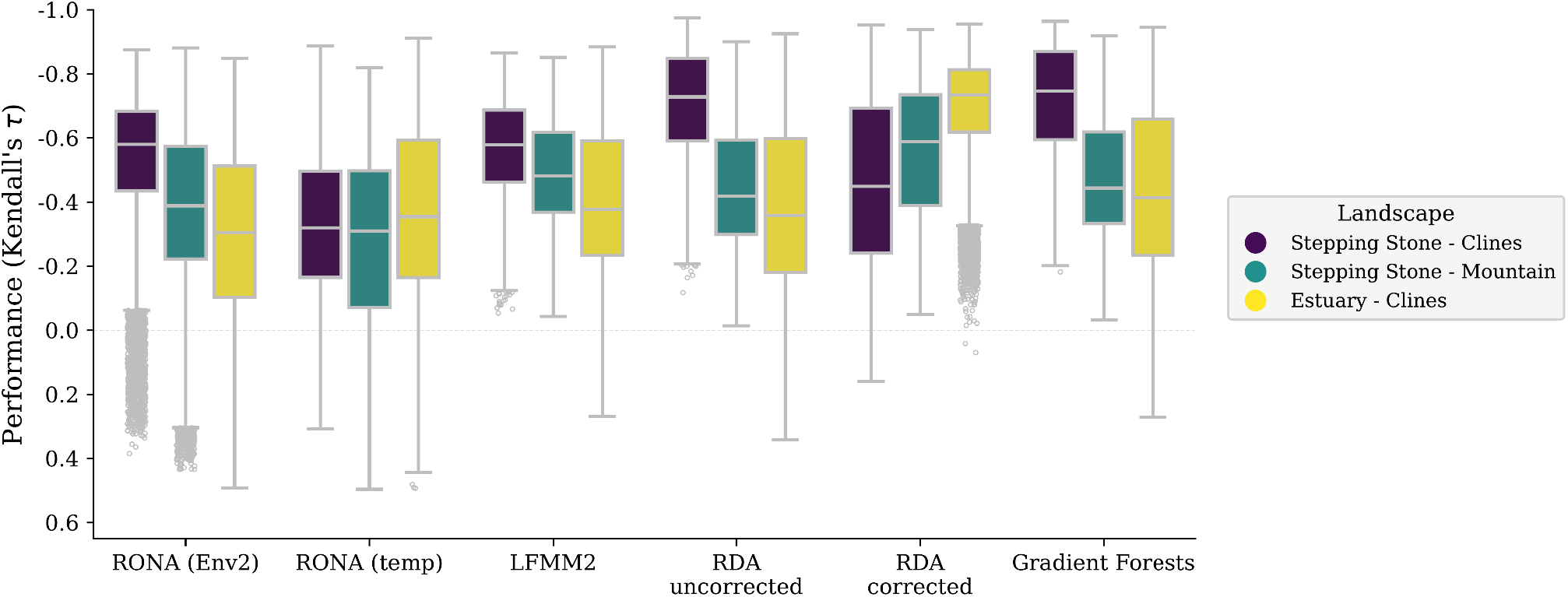
Predictive performance of genomic offset models (y-axes) is driven by the degree of local adaptation (A) and the spatial patterns of adaptive environments across the landscape (B, C). For each model, a median value from performance scores from 100 common gardens is shown for A and B; C shows scores across all common gardens for each model (note that y-axes are inverted, as more negative values have higher performance). Data included in these figures was processed through the *Adaptive Environment* workflow but only includes models trained using 2-trait simulations and *all* loci. Code to create (A) and (B) can be found in SC 02.02.02; code to create (C) can be found in SC 02.02.01.

Across landscapes, offset methods generally had higher performance in *Stepping Stone - Clines* landscapes than *Stepping Stone - Mountain* landscapes (Fig. 2B) despite similar levels of *LA*_ΔSA_ (Fig. 2A). Offset methods also generally performed better in the two *Stepping Stone* landscapes than the *Estuary - Clines* landscape (Fig. 2B). However, there were some interactions between method and landscape (Fig. 2C). For instance, RDA-corrected performed better in the *Estuary - Clines* compared to the two *Stepping Stones* landscapes, while the RDA-uncorrected showed the opposite pattern: performance was higher in the two *Stepping Stones* landscapes compared to *Estuary - Clines*.

The performance of methods was similar across genic levels but increased slightly as the number of QTNs underlying adaptation became more polygenic (Fig. S8). Additionally, while demography primarily influenced population differentiation across the landscape with little impact on *LA*_ΔSA_ within simulations (Table S2 in Lotterhos 2023), migration breaks between populations and latitudinal clines in population size generally decreased offset performance for LFMM2_offset_, GF_offset_, and RDA-uncorrected (Fig. S9).

### Q2 - How is offset performance affected by the proportion of clinal alleles in the data?

The sum of squares from Eq. 1 indicated that the proportion of clinal alleles did not account for meaningful variation in offset performance (Table S1). Even so, results from an ANOVA model with just the proportion of clinal loci as explanatory variables (Eq. 2) indicated that *p*_*cNeut*_ accounted for 4.14-9.65 times the variation than did *p*_*cQTN*_ for GF_offset_, LFMM2_offset_, and RDA-corrected. For GF_offset_ and RDA-uncorrected, *p*_*cNeut,Env2*_ accounted for >16% of the sum of squares (Table S2, Fig. S10).

Overall, relationships between performance and *p*_*cNeut*_ (second column, Fig. S11) had stronger relationships than between performance and *p*_*cQTN*_ (first column, Fig S11). However, sometimes performance increased with *p*_*cNeut*_ and sometimes it decreased, depending on the method (Fig. S11), indicating that each method is differentially sensitive to clinal alleles in the data. Ultimately, strong population genetic structure along environmental clines in 2-trait simulations (Fig. S12) drove relationships with *p*_*cNeut*_ (Fig. S13) which in turn drove relationships with performance (Fig. S14, Fig. S11).

### Q3 - Is method performance driven by causal loci or by genome-wide patterns of Isolation-By-Environment?

Overall, 1- and 2-trait *Adaptive Environment* models had relatively similar performance among marker sets. For instance, models trained using *all* or *neutral* markers had similar performance while models trained using *adaptive* markers performed slightly higher than the other sets. The median increase in performance from *adaptive* compared to *all* or *neutral* models was less than 3%. In total, using *adaptive* markers outperformed 68% of models using *neutral* markers and 67% of models using *all* markers, while 74% of models using *all* markers outperformed *neutral* models (Fig. 3A-C). For RDA-corrected the *neutral* markers performed slightly better than either *adaptive* or *all* markers in 2-trait evaluations (Fig. 3E). A*daptive* markers from 6-trait evaluations provided varied performance advantages across methods (Fig. 4).

**Figure 3.**
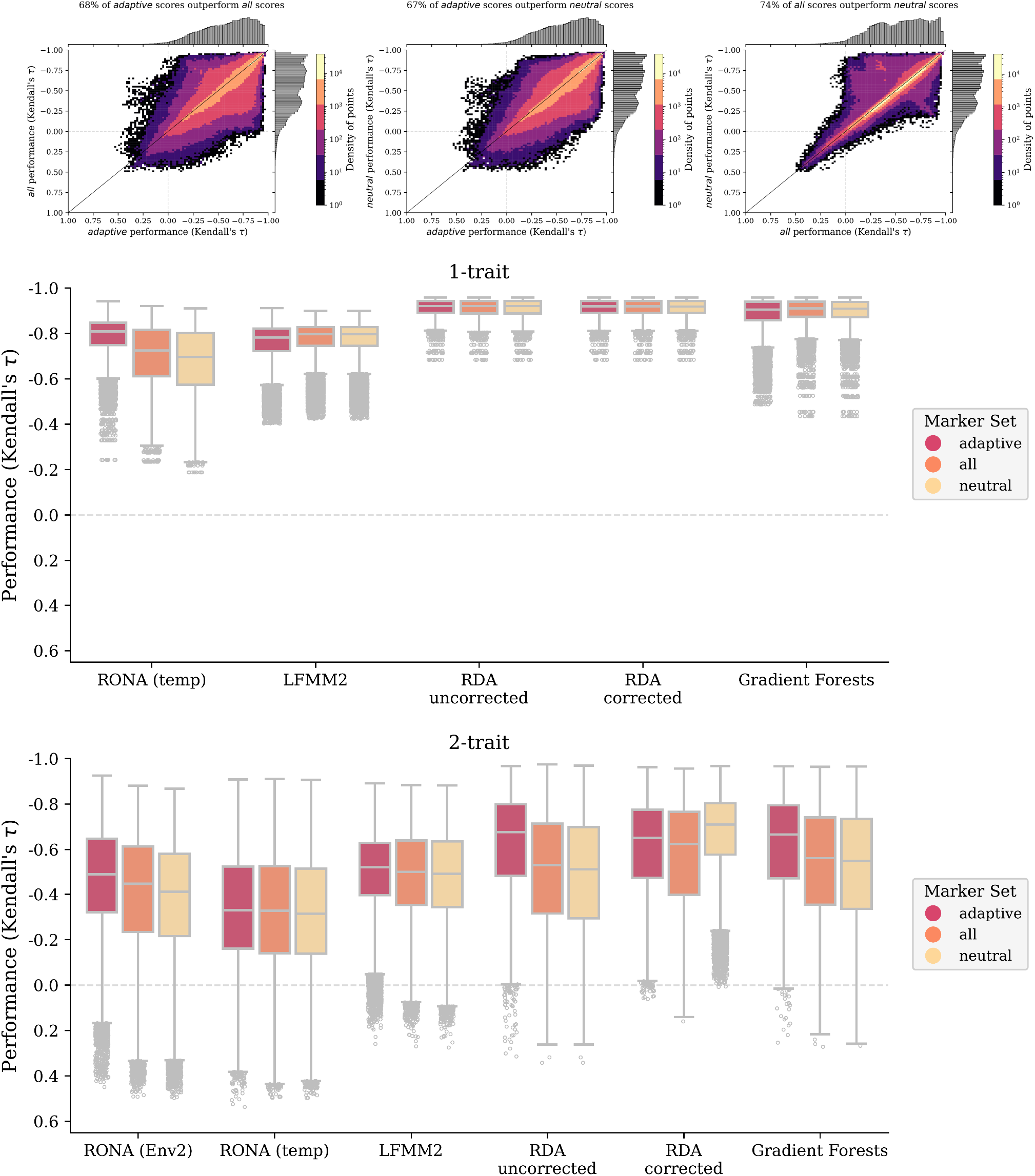
Comparison of marker choice across genomic offset methods for 1- and 2-trait simulations. A-C are scatterplots of pairwise comparisons of performance between marker sets (histograms in each margin) from both 1- and 2-trait models where density of points is indicated by color in legend (note color scale is different for each figure to accentuate patterns in data). D-E are boxplots from the same data in A-C separated by individual traits. Data included in these figures is from all 1- and 2-trait models from the *Adaptive Environment* workflow. Code to create these figures can be found in SC 02.02.03.

**Figure 4.**
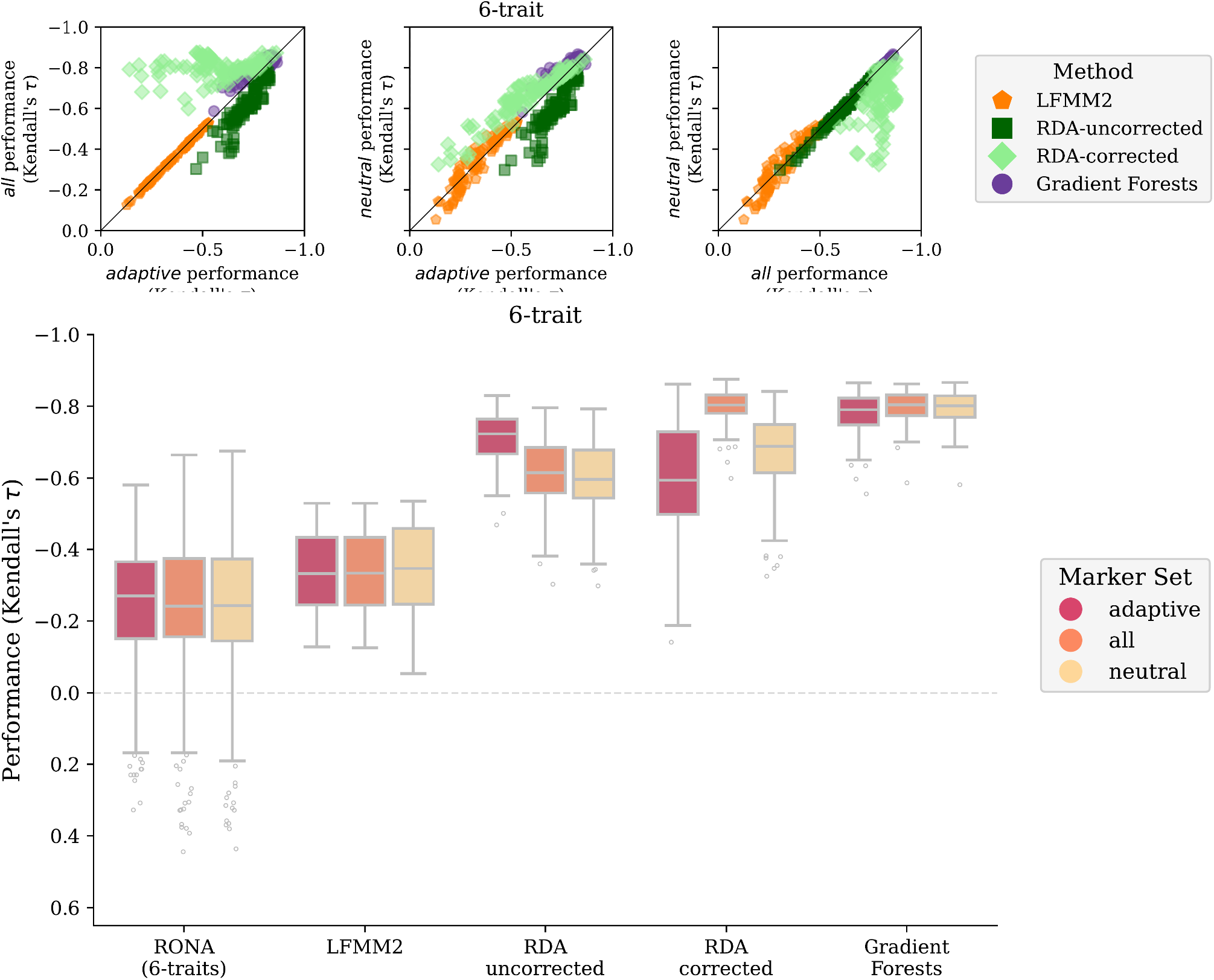
Comparison of marker choice across genomic offset methods for the 6-trait simulation. A-C are scatterplots of pairwise comparisons of performance between marker sets (RONA is not shown, except in SN 02.05.10). D are boxplots from the same data in A-C (RONA_6-traits_ is the combined performance across all six environmental models). Data included in this figure is from the 6-trait models processed through the *Adaptive Environment* workflow. Note there is only one 6-trait replicate, and variation within figures represents the performance across 100 common gardens for each method. Code to create these figures can be found in SN 02.05.10.

The *adaptive* marker sets had relatively elevated levels of *IBE* compared to sets of *neutral* or *all* markers in 1- and 2-trait simulations, but levels of *IBE* were nonetheless quite similar between marker sets (Fig. S15). Consequently, performance of models trained with *adaptive* markers generally had stronger relationships with *IBE* 786 than *LA*_ΔSA_ but this was not the case for models trained with either *all* or *neutral* markers (Fig. S16).

Intriguingly, levels of *IBE* found within a landscape (Fig. S17A) did not correspond to the degree of *LA*_ΔSA_ that developed (Fig. S17B). Even so, while *IBE* was generally unrelated to *LA*_ΔSA_ across all simulations, there were generally positive relationships between *IBE* and *LA*_ΔSA_ when controlling for the number of traits and differences in strengths of selection (Fig. S18). As such, *IBE* from *all* markers explained very little variation in performance when added as a factor to the ANOVA model from Eq. 1 (SC 801 02.02.01), but accounted for some variation in ANOVA models with only *LA*_ΔSA_ and *IBE* as explanatory variables (0-34% for *IBE* vs 0-804 74% for *LA*_ΔSA_; Table S3). Except for RONA, the differences in performance between two 806 models trained with different marker sets was generally unrelated to the differences in *IBE* 808 between the two marker sets used to train the models (Fig. S19).

Together these results indicate that while higher degrees of local adaptation may lead to increased levels of *IBE* in the genome, the signal of *IBE* of input markers generally has minimal and varied impact on performance differences for the scenarios evaluated here. Alternatively, the levels of *IBE* present in the simulated genomes may exceed a minimum 818 threshold of *IBE*, beyond which differences in performance between marker sets are minimized.

### Q4 - What is the variation of model performance across the landscape?

All 1- and 2-trait models exhibited variation in the predictive performance across gardens within a landscape, from essentially no predictive performance to very high predictive performance (Fig. S20, Fig. S21, Fig. S22, Fig. S23). Variation in performance was also observed for 6-trait models (Fig. 4).

While there was variability in predictive performance of 1- and 2-trait models within each landscape, in many cases the best performing models had the lowest levels of performance variation (Figs. S24, S25, S26). Ultimately we found no strong indicator for predicting when a model will be highly variable. Indeed, while performance generally increased with *LA*_ΔSA_ (Fig. 2), variability in performance was not strongly related to the variability in deme-level *LA* on the landscape (Figs. S27, S28, S29). Despite *LA*_ΔSA_ driving performance more generally (from Q1), this indicates that variation in model performance across the landscape is not strongly driven by metapopulation levels of, nor deme-level variation in, *LA*.

### Q5 - How does the addition of non-adaptive nuisance environments in training affect performance?

Training offset models with the addition of non-adaptive nuisance environmental variables generally reduced offset method performance (Fig. 5). This decline was most dramatic for offset trained on 1-trait simulations (Fig. 5A) compared to the decline observed for 2-trait (Fig. 5B) and 6-trait (Fig. 5C) simulations. The only instances for which median performance did not decrease monotonically with nuisance level were for 2-trait simulations evaluated with GF_offset_ (Fig. S30).

**Figure 5.**
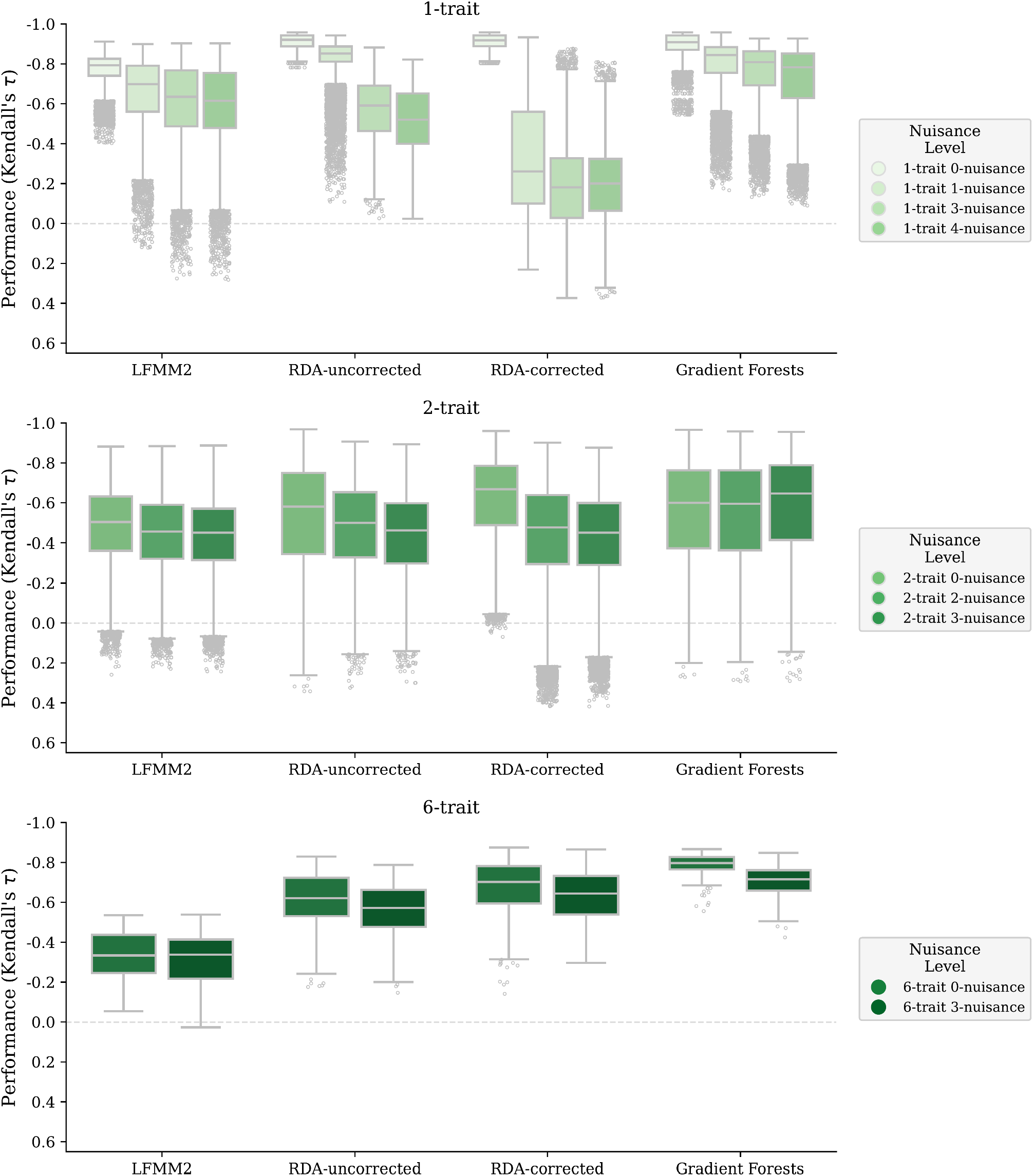
Effect of non-adaptive nuisance environmental variables on offset performance. Shown are evaluations of offsets from 1- and 2-trait models trained using only adaptive environments (*0-nuisance*) or with adaptive environments and the addition of *N*>0 non-adaptive environmental variables (*N-nuisance*). RONA is not shown because it is univariate with respect to environmental variables. Nuisance variables are listed in Table 2. Code to create figures can be found in SC 02.02.06 and SC 02.02.08.

Overall, landscape had the most influence over performance differences due to non-adaptive nuisance environments (Fig. S30), whereas there was little difference across other simulation parameters (not shown except in SC 02.02.06). Even so, *adaptive* markers seemed to provide some advantages in the presence of nuisance environments, particularly for 1-trait datasets where the advantages were more substantial compared to 2-trait datasets (Fig. S31, Fig. S32).

In some cases, the rankings of weighted environmental importance output from GF ranked nuisance variables higher than at least one adaptive environment (Table S4). Across *1-* and *2-trait N-nuisance* models trained with *all* markers, GF incorrectly ranked environmental drivers in 26.9% (133/495) of the cases. Rankings improved somewhat for models trained with *adaptive* markers, incorrectly ranking environmental variables in 20.6% (102/495) of the cases (Table S4).

### Q6 - How well do offset models extrapolate to novel environments outside the range of environments used in training?

The datasets that had the greatest within-landscape performance (i.e., those with higher levels of *LA*_ΔSA_) were also those that experienced the steepest decline in performance with increasing climate novelty (red shade, Fig. 6). Importantly, declines in performance for datasets with greater *LA*_ΔSA_ were not due to instances where all populations had zero fitness (and thus performance was undefined and manually set to 0; Supplemental Note S4, Fig. S33). Despite little change in the median performance for datasets with low levels of LA, most performance scores from these datasets were below Kendall’s **τ** = 0.5, and therefore had little predictive value in novelty scenarios.

**Figure 6.**
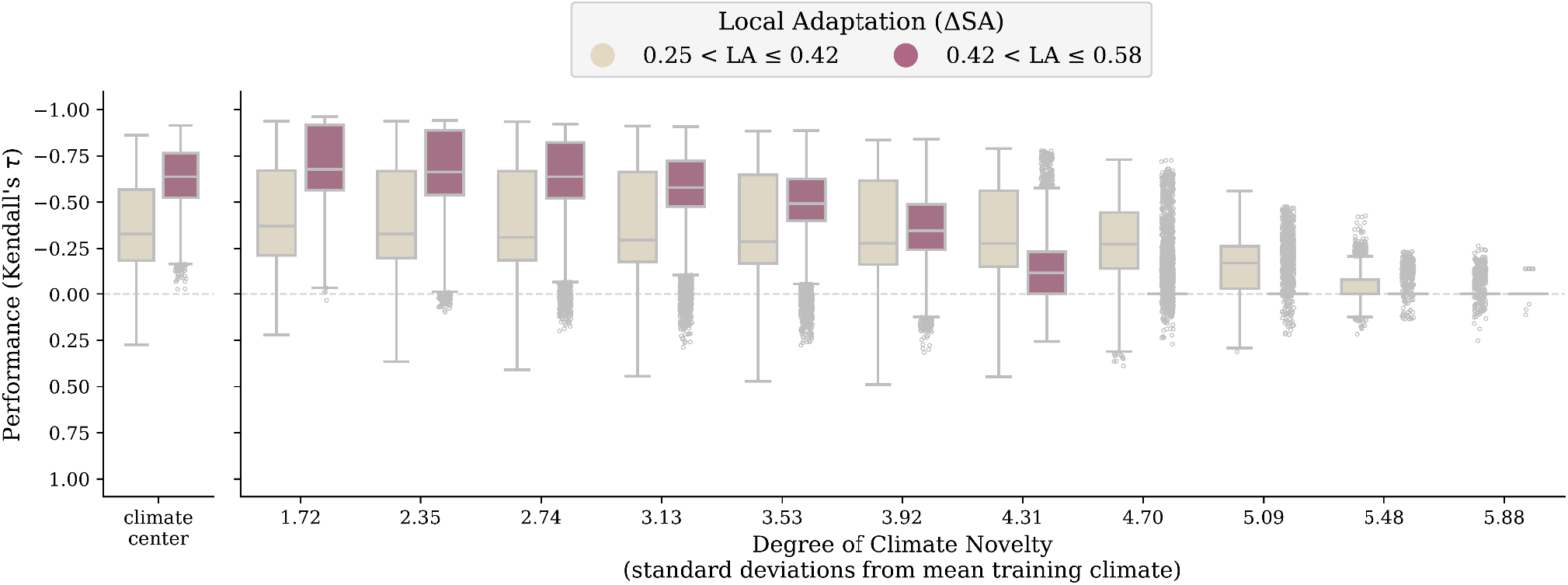
Performance decays with climate novelty relative to training data. Shown is model performance (y-axes) across methods at climate center (A) and across common gardens each representing increasing degrees of climate novelty relative to training data (x-axis of B) where all 100 populations have been transplanted. The standard deviation values (x-axis, B) are applicable to all environments for all landscapes except for *Env2* in the *Stepping Stone - Mountain* landscape; the corresponding standard deviations are 1.55, 2.12, 2.47, 2.82, 3.18, 3.53, 3.88, 4.24, 4.60, 4.95, 5.3. When fitness for all transplanted individuals was zero, a model’s performance was undefined and manually set to 0; no method predicted a single offset value for all populations in these situations. Setting undefined performance to 0 did not substantially impact patterns between performance and climate novelty, and is explored in Supplemental Text S3. Data included in this figure are from models trained using 1- and 2-trait simulations from the *Climate Novelty* workflow, which excludes both RONA_temp_ and RONA_Env2_. Code used to create this figure can be found in SC 02.04.05.

Advantages of *adaptive* marker sets were much less prevalent across methods for *Climate Novelty* scenario performance than either *Adaptive Environment* or *Nuisance Environment* scenarios (Fig. S34).

## 4 Discussion

Solutions are needed to mitigate the negative impacts of global change on biodiversity. In the last decade, genomic offset methods have been identified as a complement to other ecological forecasting models because they incorporate intraspecific variation (Keller & Fitzpatrick, 2015; Capblancq et al., 2020; Rellstab et al., 2021). Our evaluations show that offset methods are differentially impacted by both the evolutionary history of sampled populations as well as the decisions made during model training. Our analyses emphasize the importance of sampling locally adapted populations, identifying the drivers underlying environmental selection pressures *a priori*, and restricting offset projections to climates similar to those used in training. Below, we discuss the implications of these findings towards restoration, conservation, and the management of biodiversity.

### 4.1 The importance of local adaptation

A basic assumption of genomic offset methods is that the sampled populations are adapted to their local environment (Rellstab et al., 2016, 2021), but this assumption has not been formally tested. Our analyses show that indeed the degree of local adaptation (*LA*_ΔSA_) is one of the primary factors that determine model performance for most methods. A value of *LA*_ΔSA_ ∼ 0.5 indicates that fitness in demes is on average 50% higher in sympatry than allopatry. Values of *LA*_ΔSA_ represent the average deme-level magnitudes of *LA*_ΔHA_ and *LA*_ΔLF_ across the metapopulation (Blanquart et al., 2013). Previous metaanalyses of studies measuring local adaptation of natural populations have used different measures of *LA* from the ones we calculate here, but do show that some species evolve large fitness differences among populations (Hereford, 2009; Leimu & Fischer, 2008). Given the prevalence of *LA* found previously (Hereford, 2009; Leimu & Fisher, 2010), we may therefore expect some genomic offset methods to do reasonably well when local adaptation in the metapopulation is high (i.e., when *LA*_ΔSA_ > 0.5).

### 4.2 The importance of the signals within genomic marker sets

Because of the assumption that locally adapted populations will be necessary for satisfactory model performance, initial implementations of genomic offset models focussed on putatively adaptive markers where this signal may be strongest (Keller & Fitzpatrick, 2015; Rellstab et al., 2016). More recently, investigators have varied the set of markers used to train models but have found little influence on performance (Fitzpatrick et al., 2021; Lachmuth, Capblancq, Keller, et al., 2023; Láruson et al., 2022; Lind et al., 2024). Our results are similar to previous investigations, finding that the *adaptive* marker sets provide minimal advantage over *all* or *neutral* marker sets, but not universally or by great margins.

One hypothesis put forth as to why adaptive marker sets perform similarly to all markers is that genome-wide data captures sufficient signatures of IBE (Lachmuth, Capblancq, Keller, et al., 2023; Lind et al., 2024). Our analysis found weak positive relationships between performance and levels of *IBE* within marker sets. Even so, and except for RONA, there were no universal relationships within methods between the difference in *IBE* of marker sets and the difference in performance of the models trained with these markers.

While we found little impact of levels of *IBE* on overall performance, the way in which we measured IBE may have masked causative relationships. For instance, in our measure of IBE we correlated environmental distance with pairwise *F* _ST_. In doing so, our measure of IBE distills genetic distance down to a single value from a large number of loci, and gives less weight to loci with rare alleles. In future studies, creating a marker set by ranking loci by single-locus measures of IBE offers another opportunity to understand the impact of IBE on performance. Such marker sets could be used to compare to performance from putatively adaptive marker sets or marker sets composed of all or random loci. Empirical datasets will also be able to specifically address geographical distances while quantifying IBE (e.g., Bradburd et al., 2013).

While measures of IBE are one signal remaining to be explored in future analyses, the proportion of clinal neutral loci within marker sets was shown to impact performance, sometimes being positively related to performance and sometimes negatively depending on the context. These and other signals within data that could improve or mislead offset models also warrant further investigation.

### 4.3 The importance of adaptive environmental variables

In empirical settings, the environmental drivers of local adaptation are rarely known *a priori*. Even so, our results emphasize the importance of identifying these variables before training offset models, as there were often declines in performance between models trained using only adaptive environmental variables (*0-nuisance*) and those trained using additional non-adaptive nuisance environmental variables (*N-nuisance*).

The importance of identifying these selective environments may be particularly germane to two general empirical scenarios. In the first empirical scenario,, sparsely sampling an environmentally heterogeneous range may enrich genetic signals (e.g., coincident population structure) most correlated to environmental variables that maintain a gradient across this extent, and miss signals relevant to more local scales. In the second empirical scenario, identifying the environmental variables underlying selection is particularly important when a specific genomic offset method is ill-suited to differentiate importance among input variables. For instance, RDA (and therefore RDA_offset_) assumes that the environmental variables used to build models are not collinear; (as implemented here; Capblancq & Forester, 2021; Legendre & Legendre, 2012). Because of this, empirical datasets must be limited to a subset of available environmental measures. The process of excluding environmental variables in this way may omit signals of adaptive drivers (particularly when true drivers are not well measured), or perhaps incorporate environmental variables that do not coincide with drivers of selection. In these cases, performance is likely to decline. As such, this may indicate that methods such as RDA_offset_ are likely to perform worse in, or less uniformly across, realistic empirical settings than what our current findings suggest.

On the other hand, users of GF may be tempted to include a large number of environmental variables in training, hoping that GF can accurately attribute the correct environmental variation to adaptive genetic structure. Our results show that it is not necessarily the case that GF will give the highest importance values to the true adaptive environmental variables. Indeed, weighted feature importance scores from GF models still incorrectly ranked the adaptive environments below neutral environments in 20%-27% of the datasets, depending on which marker set was used. These importance values ultimately affect the model predictions. Including all available environmental variables may therefore negatively impact GF_offset_ performance, and could have weakened overall performance in previous empirical evaluations that used a large number of environmental measures in training (e.g., Lind et al., 2024).

There are some differences between the nuisance environmental variables 1109 implemented here and those that have been implemented previously. For instance, Láruson et al. (2022) created nuisance variables by randomly sampling a multivariate normal distribution. In contrast to findings here, Láruson et al. (2022) found that model performance was relatively unaffected with the addition of nuisance variables. The minimal influence of nuisance variables on performance found by Láruson et al. (2022) may differ from the performance declines reported here because the nuisance variables we used were spatially autocorrelated, while those from Láruson et al. (2022) were not. Inclusion of nuisance variables that are spatially autocorrelated may mislead offset models more generally than variables with little spatial autocorrelation because of the spurious relationship between environmental structure and genetic structure.

### 4.4 The effect of environmental novelty

While within-landscape performance generally increased with *LA* _ΔSA_, the datasets with the greatest levels of *LA*_ΔSA_ were also the datasets where performance declined most readily with climate novelty. This occurred because locally adapted metapopulations were under strong selection to be fine-tuned to their environment, and as a result most individuals suffered severe fitness declines with environmental change. In contrast, less locally adapted metapopulations were under weaker selection, and suffered less steep fitness declines with environmental change. This result highlights an interesting paradox: offset methods that have the highest performance in common garden transplants under current climates (because of strong local adaptation) may have the lowest performance in predicting “genomic vulnerability” as the range of climate variables become more novel compared to the ranges used in training the model.

Thus, genomic offset models are likely ill-suited for estimating fitness ranks of populations in environments that differ drastically from those used to train the models themselves. This is particularly relevant for applications of offset methods that attempt to estimate *in situ* risk of climate change to years or climate scenarios where the environment is expected to be increasingly novel. While climate novelty is often measured with respect to historical variability (e.g., Lotterhos et al., 2021; Mahony et al., 2017; Williams et al., 2007), indices of local climate change indicate that local environments in terrestrial systems could experience change in excess of three standard deviations relative to historic values (Williams et al., 2007). Similar indices in marine systems indicate potential for even greater novelty (Lotterhos et al., 2021). We observed performance declines below the analogous *z*_E_=3.13 *Climate Novelty* scenario, indicating offset predictions will likely be inaccurate in many real-world climate change predictions. These issues are also germane to measures derived from offset values (Gougherty et al., 2021; Lachmuth, Capblancq, Keller, et al., 2023; Lachmuth, Capblancq, Prakash, et al., 2023), which currently do not consider the degree of climate novelty in the prediction (but see DeSaix et al., 2022).

Our results present a best-case scenario for predicting performance in novel environments, as in many cases there will be biological reasons as to why climate-fitness relationships will differ in future environments from relationships measured within the contemporary climate space (see Fig. 5 in Capblancq et al., 2020). For the simulations here, the relationship between contemporary and novel environments with fitness was the same.

### 4.5 Genomic offsets in practice

Our evaluations show that genomic offset methods hold promise for predicting maladaptation to environmental change within a historical baseline, in metapopulations that evolve strong local adaptation. However, our analyses also emphasize the limits of these methods in some systems or scenarios. In practice, species that are locally adapted to measurable environmental variables will be best suited for offset methods when predicting the relative performance of populations in a contemporary common garden, but paradoxically these species may be least suited to using these methods to predict their vulnerability to novel climates.

Together, these results indicate that some genomic offset methods may be suited to guide initiatives such as near-term assisted gene flow, where targeted restoration sites within a species range have climates that are similar to those used to train offset models. Even so, our results also show that the performance of these methods are often variable across a landscape, indicating that high performance at one site does not mean the offset model will perform well at another. While genomic offset methods may be suitable for assisted gene flow initiatives, they may be less suited for assisted migration programs where populations are moved outside of their native range and environments differ from training data.

Before genomic offsets can be incorporated into management plans, considerable thought must be put into the sensitivity of model outcomes from input data (Lind et al., 2024), the uncertainty inherent in environmental or climate forecasts (Lachmuth, Capblancq, Keller, et al., 2023), as well as the degree of novelty of future climates (DeSaix et al., 2022, this study). While accurate predictions are limited for novel climates of the future, these offset methods could still be used to guide management in the intervening time in a stepwise manner where experiments can be used to validate model performance in practice. Using simulations tailored to the life history of target species also presents a promising avenue to understand limitations of these methods for specific management cases.

## Acknowledgements

This research was funded by NSF-2043905 (KEL) and Northeastern University. The funding bodies did not have any role in the design of the study, analysis, interpretation of results, or in writing of the manuscript.

## Author Contributions

KEL received funding. KEL and BML conceptualized the project and methodology. With input, editing, and feedback from KEL, BML wrote code to train and evaluate offset models, created figures, curated coding and records for archiving, and wrote the manuscript.

## Data Availability

We reference the analysis code in the text of our documents by designating Supplemental Code (SC) using a directory numbering system from our servers (as opposed to the order listed in the manuscript). Supplemental Code includes both executable scripts (*.R, *.py) as well as jupyter notebooks (*.ipynb). For example, for Script 3 in Directory 1, we refer to SC 01.03; for Notebook 5 in Subfolder 3 of Directory 2, we will refer to SC 02.03.05. Each Directory will be archived on Zenodo.org and include a citation below, which will also link to the GitHub repository. Notebooks are best viewed within a local jupyter or jupyter lab session (to enable cell output scrolling / collapsing), but can also be viewed at nbviewer.jupyter.org using the web link in the archive’s README on GitHub. Analyses were carried out primarily using python v3.8.5 and R v3.5.1 and v4.0.3. Exact package and code versions are available at the top of each notebook. More information on coding workflows and coding environments can be found in Supplemental Note S1 and Supplemental Note S2.

All directories, notebooks, and scripts can be found on GitHub, and will be archived on Zenodo. https://github.com/ModelValidationProgram/MVP-offsets

