## Supplemental Material for "The limits of predicting maladaptation to future environments with genomic data"

11 January 2024

**Running Title**: *The limits of genomic offsets*

**Keywords**: genomic offset, environmental change, climate change, assisted gene flow, genomic forecasting, restoration

***Corresponding Author**

Brandon M. Lind

Table S3 Results from Type II ANOVAs regressing two factors - degree of local adaptation (final_LA) and levels of isolation-by-environment in all marker sets) on offset performance..... .............. 28

Fig S8 Effect of polygenicity on performance of offset methods trained using all markers on simulations with two adaptive traits........ ..................................................................................... 36

Fig S9 Effect of demography on performance of offset methods trained using all markers on simulations with two adaptive traits........ ..................................................................................... 37

Fig S13 Relationship between the proportion of clinal neutral loci for *temp* (y-axes, first row) or *Env2* (y-axes, second row) with the strength of the relationship between environmental variables and axes of population genetic structure. Purple = *Stepping Stone -*

Fig S26 Variability across evaluations of genomic offsets often decreases with increasing average performance across marker set.......... 74

Fig S27 Variability across evaluations of genomic offsets is often unrelated to the variability in the degree of local adaptation across populations.. 75

Fig S28 Variability across evaluations of genomic offsets is often unrelated to the variability in the degree of local adaptation across populations.. 76

Fig S29 Variability across evaluations of genomic offsets is often unrelated to the variability in the degree of local adaptation across populations.. 77

Fig S30 Effect of non-adaptive nuisance environmental variables on offset performance faceted by landscape.. 78

Fig S31 Effect of non-adaptive nuisance environmental variables on offset performance faceted by marker set.. 79

Fig S32 Pairwise comparison of performance differences between marker sets for *Nuisance Environment* scenarios.. 84

Fig S34 Pairwise comparison of performance differences between marker sets for *Climate Novelty* scenarios.. 86

**Supplemental References 87**

### Supplemental Notes

#### S1 - Implementation of Offset Methods

See Supplemental Note S2 for specific citations of code.

##### 1.1 | Gradient Forests

For a given set of input loci (*all*, *adaptive*, or *neutral*; see Q3 in Methods), and for all workflows, Gradient Forests ($GF_{Offset}$) is trained using `ntree=500`, `corr.threshold=0.5`, and `maxLevel=`$(0.368 * \frac{N}{2})$, where $N$ is the number of populations. Using default linear extrapolation, the trained model is projected onto the landscape using the `` `predict` `` function and the same environmental values used in training. This creates the "current" projection used to calculate offset below.

The trained models are then fit to the climate of each of 100 common gardens on the landscape for the *Adaptive Environment* and *Nuisance Environment* scenarios, or to each of the 11 *Climate Novelty* scenarios. Specifically, for each garden, the `` `predict` `` function is used to take the trained model and the garden's climate to create a projection similar to that using current climate data (previous paragraph). Then the Euclidean distance is taken between the current and future projections to calculate offset.

#### 1.2 | The Risk Of Non-Adaptedness

For a given set of input loci (*all*, *adaptive*, or *neutral*; see Q3 in Methods), we first discarded any locus that did not have significant ($p \leq 0.05$) linear models relating population-level allele frequencies with environmental variables. p-values were not corrected for multiple testing. For each common garden, and once for each environmental variable, RONA offset for each population was calculated by averaging the absolute allele frequency difference between the population's current frequency and that predicted by using each locus' linear model fit using climate of the common garden,

$$\text{RONA} = \frac{1}{n}\sum_{i=1}^{n} \left|(S_{\text{present}_i} * \text{EF}_{\text{future}} + I_{\text{present}_i}) - \text{AAF}_{\text{present}_i}\right|$$

where $n$ is the total number of loci with significant linear models; $S_{\text{present}}$ and $I_{\text{present}}$ are respectively the slope and intercept from the linear model for $\text{locus}_i$ relating current climate and allele frequencies from all populations; $\text{AAF}_{\text{present}}$ is the current allele frequency for the population under consideration; and $\text{EF}_{\text{future}}$ is the environmental value for the common garden. RONA can only be calculated for a single population and environmental variable at a time.

RONA was excluded from *Nuisance Environment* and *Climate Outlier* workflows because of its poor (Fig. 2A) and variable (Fig. 2C, Fig. 4) performance from evaluations from the *Adaptive Environment* workflow.

Of note, in some instances, particularly *Adaptive Environment* datasets simulated with oligogenic architectures, there were no loci with significant linear relationships with environmental variables and these instances were given NA performance values (i.e., excluded from analyses).

#### 1.3 | Landscape and Ecological Association (LEA) Studies R package

We used the `genetic.offset` function in the LEA package to estimate LFMM2$_{offset}$ for each workflow (Fig. 1). The `genetic.offset` function was used with default settings, except for *K*, the number of subdivisions within the data. To determine *K* needed for the `genetic.offset` function for 1- and 2-trait simulations, we first used filtered SNP data (see Section 2.1) to estimate 21 principal components (PCs) using principal component analysis (PCA). Then we equated *K* to the number of PC axes that explain greater than 1.3x the variation of the next subsequent axis (see line 677-697 of [c-AnalyzeSimOutput.R] from Lotterhos, 2023). This resulted in varied *K* across simulation levels and replicates (Fig S1). For the 6-trait simulation, it was never the case that a PC axis explained >1.3x the variation explained by the previous axis, so we used the elbow rule to estimate *K*=7 (Fig S2).

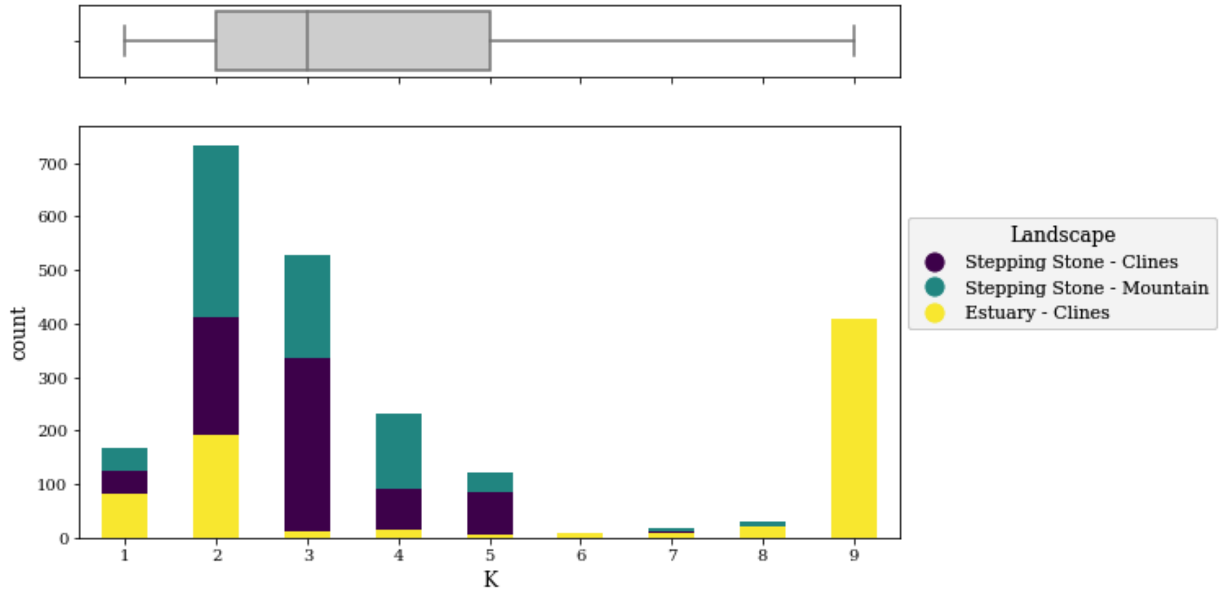

**Fig S1** Distribution of *K* used for the lfmm2 `genetic.offset` function for 1- and 2-trait simulations. *K* was estimated by determining the number of principal component axes that explain at least 1.3x times the amount of variation of the subsequent axis. Code used to create this figure can be found in SC 02.09.01.

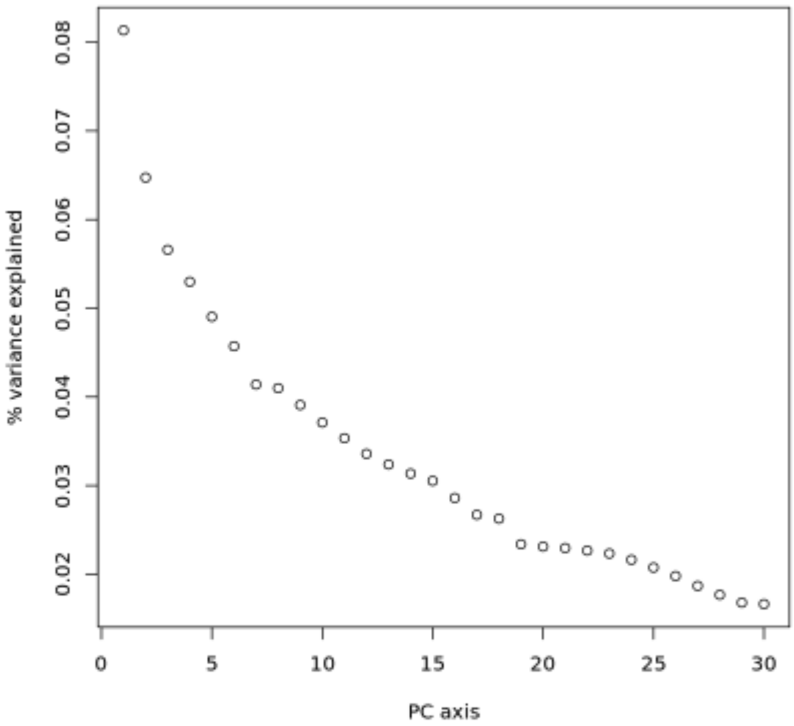

**Fig S2** Percent variance explained from principal component (PC) axes from principal component analysis of SNP data from the 6-trait simulation. The "elbow rule" was used to estimate *K*=7 for this simulation. Code used to create this figure can be found in SC 02.05.11.

#### 1.4 | Redundancy Analysis

$RDA_{offset}$ was implemented as in Capblancq & Forester (2021). Note that the environmental variables used here across workflows had minimal correlation, as required by RDA (Fig. S4). In addition to the three marker sets used as input (*all*, *adaptive*, or *neutral*; see Q3 in Methods), we also used *RDA-outliers* as input to RDA offset estimation. *RDA-outlier* loci were those from separate RDA models trained using *all* loci and adaptive environments, and were included in this study because of their use in the original implementation of $RDA_{offset}$ by Capblancq & Forester (2021). *RDA-outliers* were identified as in Capblancq et al. (2018) for loci with q-values < 0.05. For each 1-, 2-, and 6-trait simulation replicate, $RDA_{offset}$ was estimated with (RDA-corrected) and without (RDA-uncorrected) correction for population genetic structure. When correcting for structure, the loadings for the first two PCs from PCA estimated with *all* loci were used. Because *RDA-outliers* performed on par with or worse than other marker sets in 1-trait (Fig S2A), 2-trait (Fig S3B), and 6-trait (Fig S3C) evaluations from the *Adaptive Environment* workflow (Fig. 1) we focus on *all*, *neutral*, and *adaptive* marker sets for the main text.

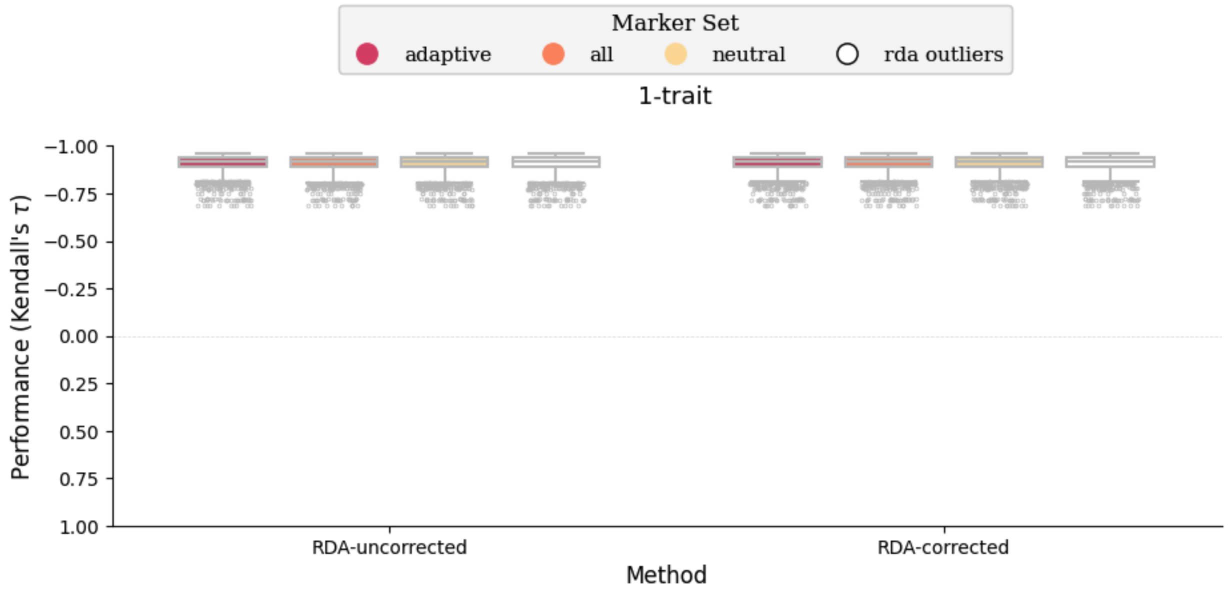
Marker Set
adaptive
all
neutral
rda outliers
1-trait
Performance (Kendall's τ)
−1.00
−0.75
−0.50
−0.25
0.00
0.25
0.50
0.75
1.00
RDA-uncorrected
RDA-corrected
Method

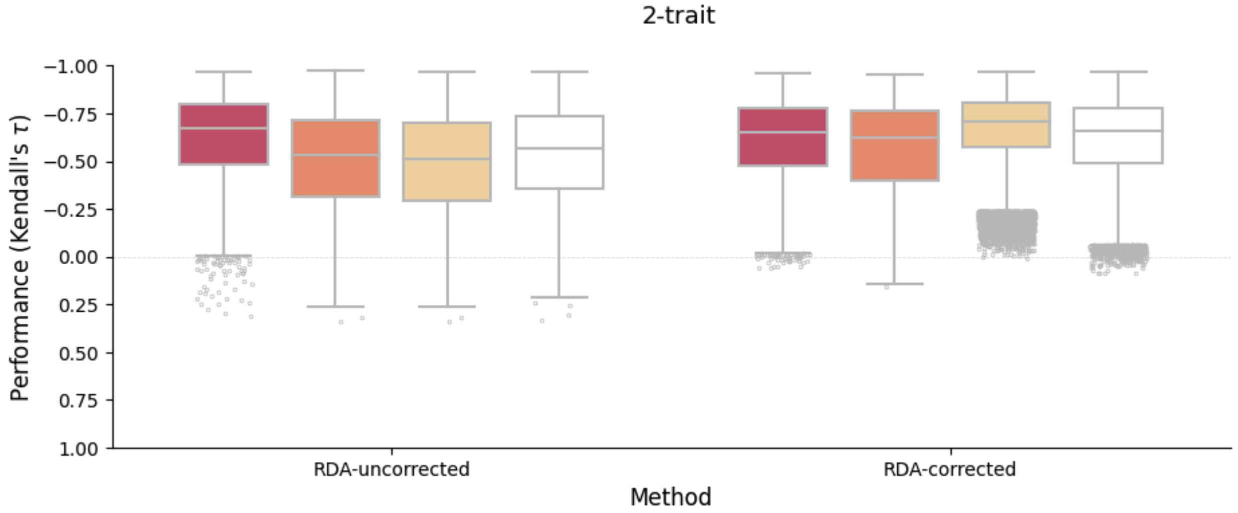
2-trait
Performance (Kendall's τ)
−1.00
−0.75
−0.50
−0.25
0.00
0.25
0.50
0.75
1.00
RDA-uncorrected
RDA-corrected
Method

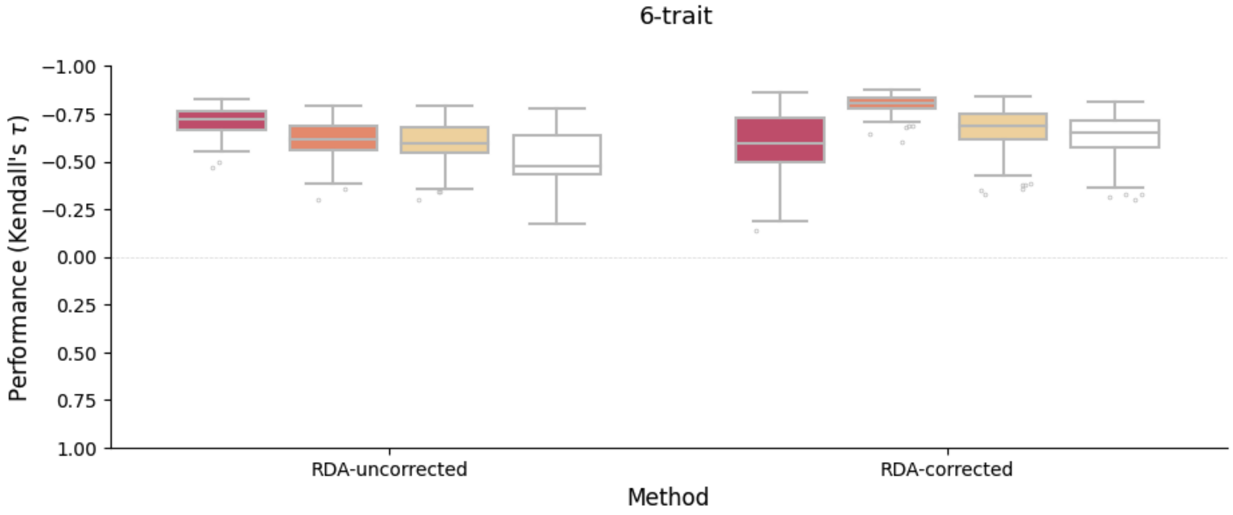
6-trait
Performance (Kendall's τ)
−1.00
−0.75
−0.50
−0.25
0.00
0.25
0.50
0.75
1.00
RDA-uncorrected
RDA-corrected
Method

**Fig S3** Performance of RDA-outlier markers are on par with other marker sets for (A) 1-trait, (B) 2-trait, and (C) 6-trait evaluations of offset estimated with (RDA-corrected) or without (RDA-uncorrected) population structure correction. Data in this figure is from the *Adaptive Environment* workflow. Code to create this figure can be found in SC 02.06.02.

### S2 - Coding workflows

Below we reference the scripts (`*.R`, `*.py`) and notebooks (`*.ipynb`) used to analyze data in this manuscript using the naming convention described in the Data Availability section (e.g., SC 05.02). Scripts are often written using only functions, instead of a linear development of code. This allows the functions to be imported/sourced in other scripts or notebooks to avoid code redundancy. At the top of all script files are detailed instructions for use. The "`main`" function in many script files gives a general outline for the code and calls all other functions.

All python scripts and notebooks are run in the "mvp_env" (python v3.8) Anaconda environment. All GF scripts are run in R within the "r35" (R v3.5) Anaconda environment . All other R code is run within the "MVP_env_R4.0.3" (R v4.0.3) Anaconda environment. All Anaconda environments can be recreated using their `.yml` files found in the code archive. These files contain all package and library versions at the time of saving. Package and library versions that were used are found at the top of each notebook - look for "Click to view session information" (python notebooks) or printouts from `sessionInfo()` (R notebooks).

1- and 2-trait simulations are often processed separately from the 6-trait simulation. Descriptions of coding workflows reflect this.

All scripts referenced by name are in the SC 01 directory.

Notebooks used to create figures and tables are not described here (but see coding archive README). Instead, these notebooks are referenced within the caption of all figures and tables, or in the main text when appropriate. These notebooks (mainly within SC 02.02 directory) rely on data processed through the coding workflows described below. Similarly, code previously described in Supplemental Note S1 is not redescribed here.

Simulation data used below within scripts and notebooks has been processed from SLiM output separately by Lotterhos (2023) into more user-friendly forms - see here for more information: https://github.com/ModelValidationProgram/MVP-NonClinalAF/tree/main/sim_ouput_20220428_metadata

#### 1.1 | The *Adaptive Environment* coding workflow

The *Adaptive Environment* workflow represents the general pipeline for processing simulations and running genomic offset methods, most other processing code is built on top of this main pipeline (i.e., scripts and notebooks source/import functions from these scripts to avoid code redundancy).

##### *1.1.1 | 1- and 2-trait simulations*

The *Adaptive Environment* pipeline is kicked off using SC 01.00, which allows the user to decide which method to run. All analyses were generally run in batches of 225 simulation levels (one replicate per level). SC 01.00 can call SC 01.01 (for GF), SC 01.05 (for RONA), SC 01.10 (for LFMM), SC 01.07 (for pairwise $F_{ST}$), or scripts related to RDA (more details below).

$GF_{offset}$ : SC 01.01 processes the data into formats suitable for GF input. This includes converting genotype data into derived allele frequencies, asserting MAF cutoffs, and reformatting environmental data. This script creates `.sh` files for the slurm HPC and trains GF models using `` `MVP_gf_training_script.R` ``. The slurm `.sh` files call SC 01.02, which takes the trained GF model and predicts offset to each of the 100 environments (population sources) on the landscape using `` `MVP_gf_fitting_script.R` ``. Performance of GF offset predictions are then validated using SC 01.03. Performance results are saved in a nested dictionary. Environmental importance is extracted from each GF model using SC 01.04 within SC 02.10.02.

RONA : Using files created from SC 01.01, SC 01.05 creates files suitable for RONA analyses and calculates RONA itself. Performance of RONA is validated with SC 01.06. As with GF, performance results are saved in a nested dictionary.

$LFMM_{offset}$ : SC 01.10 creates files suitable for LFMM in R and submits jobs to the slurm HPC to train LFMM with `` `MVP_process_lfmm.R` ``. SC 01.10 also submits SC 01.11 to validate LFMM offsets. `` `MVP_watch_for_failure_of_train_lfmm2_offset.py` `` watches for failed jobs and reruns them. Performance of LFMM is validated in SC 01.11. As with GF and RONA, performance results are saved in a nested dictionary.

$RDA_{offset}$ : `` `MVP_pooled_pca_and_rda.R` `` creates principal component analysis data and RDA objects using allele frequencies of *all* loci; it also creates additional files needed downstream. Next, SC 01.12 is run to estimate RDA offset. Performance of $RDA_{offset}$ is validated with SC 01.13. As with GF, RONA, and LFMM, performance results are saved in a nested dictionary.

Nested dictionaries containing validation results from each method are reformatted and combined into a single object in notebooks within the SC 02.01.00

directory. These combined objects are used throughout remaining analyses in jupyter notebooks found in subdirectories of SC 02.

##### *1.1.2 | 6-trait simulation*

The 6-trait simulation was processed through the *Adaptive Environment* workflow using code found in the SC 02.05 directory. 6-trait simulations needed extra formatting in order to be comparable to the 1- and 2-trait evaluations. First, SC 02.05.00 assigns individuals to populations using a gridded system. Population-level environmental values are the average climate from assigned individuals on the landscape (each environmental variable is averaged independently). Genetic and environmental data was formatted as with 1- and 2-trait simulations. Fitness for each population in each environment was calculated using `MVP_climate_outlier_fitness_calculator.R`. The script `MVP_climate_outlier_fitness_calculator.R` was validated against previous fitness estimates from 1- and 2-trait simulations in SC 02.05.01.

GF was trained using the same script as 1- and 2-trait simulations (`MVP_gf_training_script.R`). GF offset was predicted manually in SC 02.05.02, and validated manually in SC 02.05.03. In SC 02.05.04 - 02.05.05 LFMM was trained and validated manually. Similarly, RDA was trained and validated in SC 02.05.06 - SC 02.05.07, and RONA trained and validated in SC 02.05.08 - SC 02.05.09.

#### 1.2 | The *Climate Novelty* coding workflow

Fitness was calculated for 1- and 2-trait populations within the *Climate Novelty* scenarios (x-axis, Fig. 6, Supplemental Note S3) using `MVP_climate_outlier_fitness_calculator.R` in 02.04.01.

Using 1- and 2-trait offset models output from the *Adaptive Environment* workflow, the following code predicted offset to *Climate Novelty* scenarios (GF: SC 01.14; LFMM: SC 01.16; RDA: SC 01.18; and RONA: SC 01.20) which was subsequently validated against known fitness (GF: SC 01.15; LFMM: SC 01.17; RDA: SC 01.19; and RONA: SC 01.21). A few examples of code executions are shown in SC 02.04.03.

Fitness of 6-trait populations for *Climate Novelty* scenarios was calculated in SC 02.04.06. 6-trait GF models used the same scripts as 1- and 2-trait runs (SC 01.14 - SC 01.15); executed from SC 02.04.07. Commands to train LFMM were created in SC 02.04.07, which called on `MVP_complex_sims_process_lfmm.R`. RDA was trained manually in SC 02.04.07. Offset from both LFMM and RDA were validated manually in SC 02.04.08.

#### 1.3 | The *Nuisance Environment* coding workflow

Environmental files for *Nuisance Environment* scenarios were created in SC 02.07.02.02.

Files for 1- and 2-trait simulations were created in SC 02.07.02.01 to train GF using `MVP_gf_training_script.R`. SC 01.02 and SC 01.03 are used for predicting and validating GF offset, respectively, executed in SC 02.07.02.02. Code for LFMM was executed in SC 02.07.02.07 and used `MVP_process_lfmm.R` for training and SC 01.11 for validation. Commands for RDA were created in SC 02.07.02.06 similarly to *Adaptive Environment* workflow (calling `MVP_pooled_pca_and_rda.R`) and used `MVP_nuisance_RDA_offset.R` for training and `MVP_nuisance_rda_validation.py` for validation.

6-trait sims were processed for GF exactly as they were for 6-trait data in the *Adaptive Environment* workflow (with updated environmental files) and executed in SC 02.07.02.03, SC 02.07.02.04, and validated manually in SC 02.07.02.10. Code to train both LFMM and RDA was executed in SC 02.07.02.12, which called on `MVP_complex_sims_process_lfmm.R` for LFMM. LFMM was validated in SC 02.07.02.13; RDA was validated in SC 02.07.02.14.

#### 1.4 | Misc

`MVP_summary_functions.py` contains much of the API used within notebooks for loading and filtering data as well as creating figures. It is often imported using the alias `mvp` within python scripts and notebooks.

### S3 - Defining *Climate Novelty* scenarios

To understand if genomic offset models maintained predictive performance in environments differentiated from training environments, we created 11 climates, each progressively more distant from the mean training environment. Specifically, for each environmental variable, we used a standardized set of z-scores ($z_E \in \{1.72, 2.35, 2.74, 3.13, 3.53, 3.92, 4.31, 4.70, 5.09, 5.48, 5.88\}$) to calculate corresponding environmental values. In other words, we used the distribution of the within-landscape values from which to identify the appropriate value for a given z-score for each environmental variable independently. The *temp* environment and all six of the 6-trait environments were given positive values for *Climate Novelty* scenarios, and *Env2* was given negative values.

Novelty climates for 6-trait and 2-trait evaluations are shown Fig S5A and B, respectively. In this and other figures related to performance in *Climate Novelty* scenarios, we also include $z_E$=0.00 for comparison of novelty climates to the mean training climate (i.e., climate center). We chose *z*-scores over Mahalanobis distances because of 1) the reduced correlation structure among environmental variables (where z-scores and Mahalanobis distances should be roughly equivalent; Fig. S4), and 2) the large number of combinations of values from environmental variables that could be used for a given Mahalanobis distance. The standard deviation values that we used are applicable to all environments and for all landscapes except for *Env2* in the *Stepping Stone - Mountain* landscape; the corresponding standard deviations for this case are $z_E \in \{1.55, 2.12, 2.47, 2.82, 3.18, 3.53, 3.88, 4.24, 4.60, 4.95, 5.3\}$.

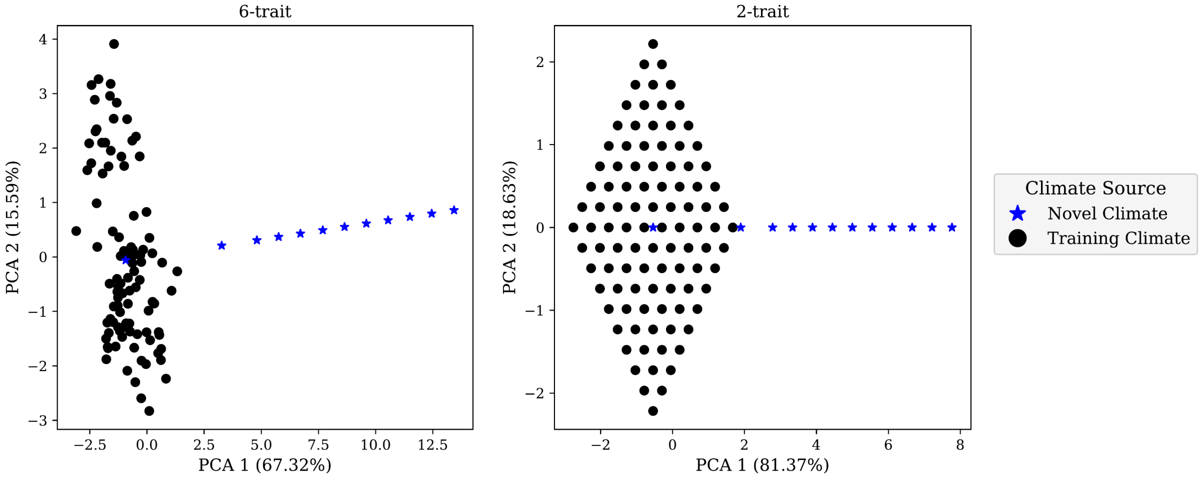

**Fig S5** Differentiation of *Climate Novelty* environments (blue stars, including climate center) from within-landscape environments (black circles) using Principal Component Analysis (PCA) of environmental data. Environmental data is centered and standardized relative to the within-landscape environmental values. Scatter plots show the first two principal components (PCs) of environmental data used to evaluate 6-trait (A) and 2-trait (B) *Climate Novelty* scenarios. There is no figure for 1-trait evaluations because there would only be one PC axis. Code to create these figures can be found in SC 02.04.10.

### S4 - Missing data in *Climate Novelty* evaluations

When calculating fitness of populations in *Climate Novelty* scenarios, it could be the case that all populations have zero fitness because of the extremity of the novel climate. In these cases the calculation of performance is technically undefined due to the lack of variability in one of the vectors (i.e., the code returns "NAN"), but for Figure 7 we replaced these undefined values with 0 (because there was no predictive performance of the offset method). We refer to these cases as missing data below. It is therefore important to explore the effect of these missing data points on patterns observed between performance and climate novelty (i.e., in the context of Fig. 7 of the main text) to ensure patterns before and after setting missing data to 0 do not affect inferences.

To understand impacts of missing data, we created figures that grouped simulation and experimental levels across novelty scenarios (Fig. S33). We also printed out specific scenarios in the code (SC 02.04.05). Importantly, missing data is not substantial until *Climate Novelty (CN) Scenario 4.31*, which is preceded by the drop in performance from datasets with elevated $LA_{\Delta SA}$. After *CN Scenario 4.31* missing data begins to increase because of climate novelty, first with datasets where high levels of $LA_{\Delta SA}$ take place through oligogenic architectures, then missing data is more uniform across simulation and experimental parameters for the remaining *CN Scenarios* (Fig. S33). (Before *CN Scenario 4.31*, missing data is not due to all populations having zero fitness - instead missing data is primarily due to 1-trait oligogenic scenarios evaluated by RONA where there are no *adaptive* alleles with significant clines with *temp* in the *Estuary - Clines* landscape (Fig. S33; SC 02.04.05).) Finally, we also explored patterns presented in Fig. 6 before setting undefined performance scores to zero and found nearly identical trends (not shown).

(Fig S33)

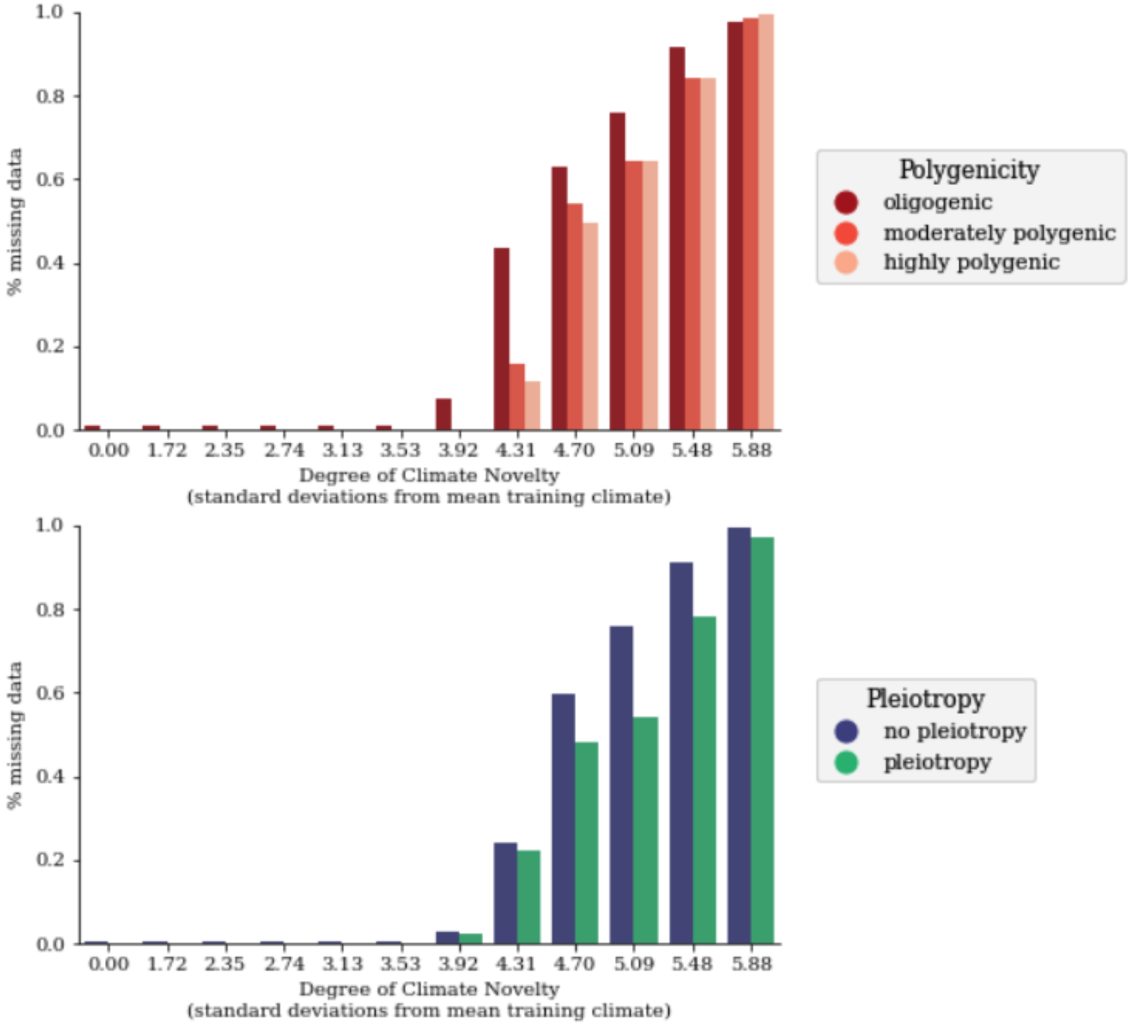
% missing data
1.0
0.8
0.6
0.4
0.2
0.0
0.00
1.72
2.35
2.74
3.13
3.53
3.92
4.31
4.70
5.09
5.48
5.88
Degree of Climate Novelty
(standard deviations from mean training climate)
Polygenicity
oligogenic
moderately polygenic
highly polygenic
% missing data
1.0
0.8
0.6
0.4
0.2
0.0
0.00
1.72
2.35
2.74
3.13
3.53
3.92
4.31
4.70
5.09
5.48
5.88
Degree of Climate Novelty
(standard deviations from mean training climate)
Pleiotropy
no pleiotropy
pleiotropy

(Fig S33 continued)

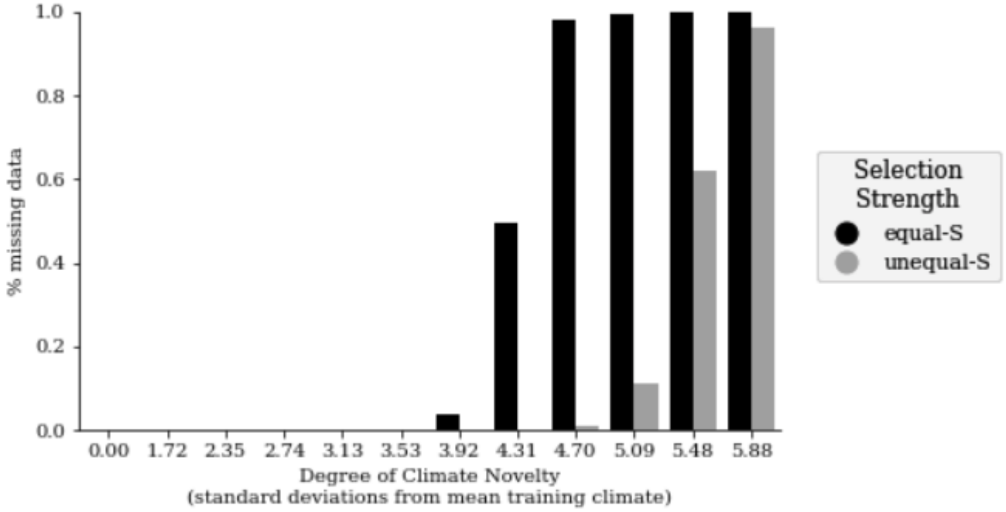

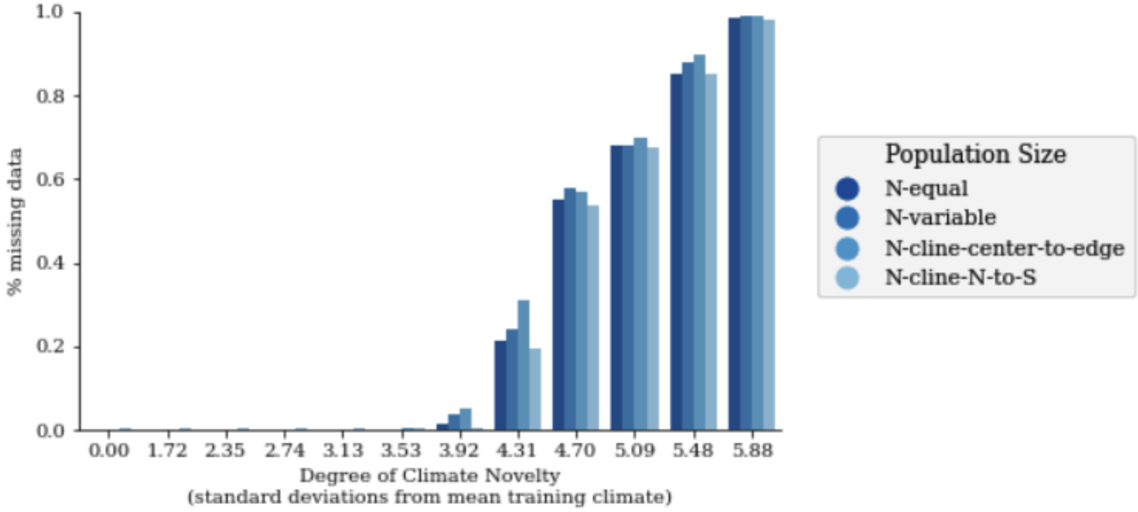

(Fig S33 continued)

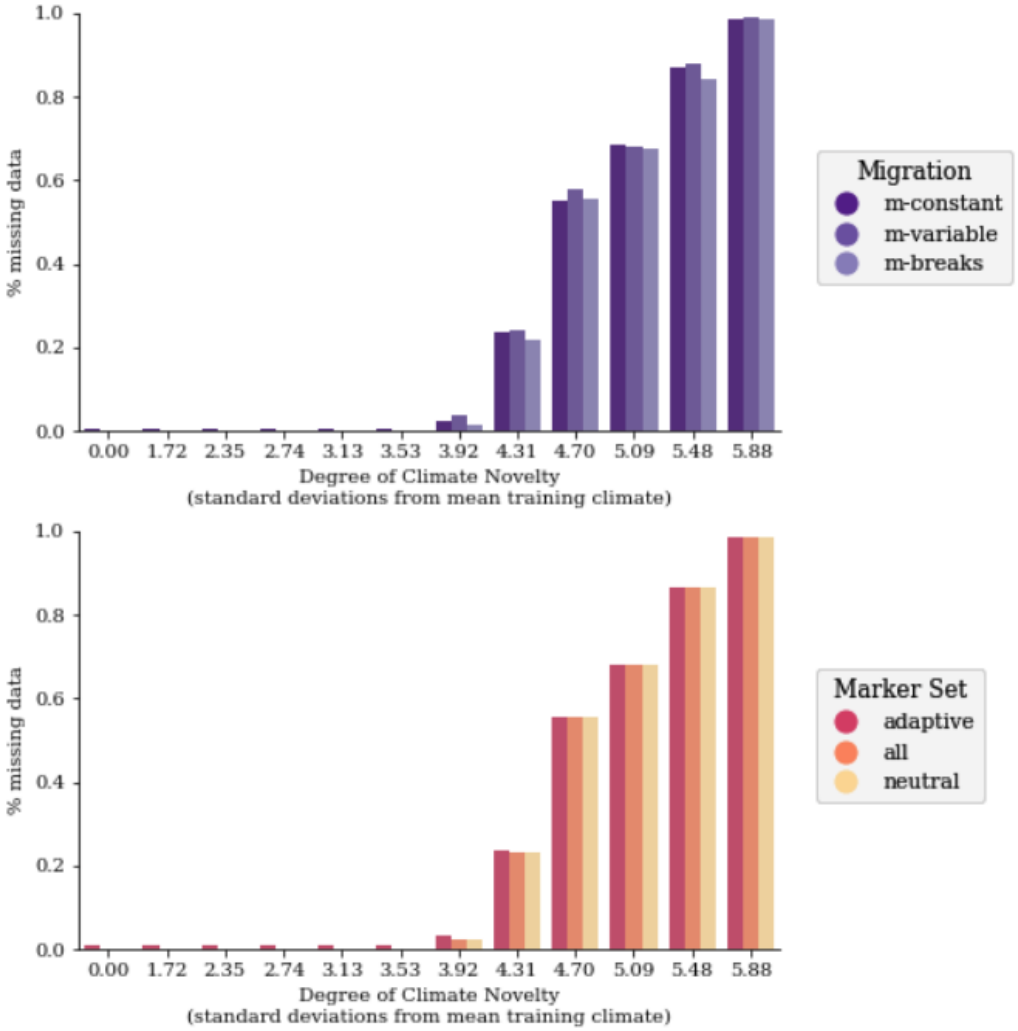

(Fig S33 continued)

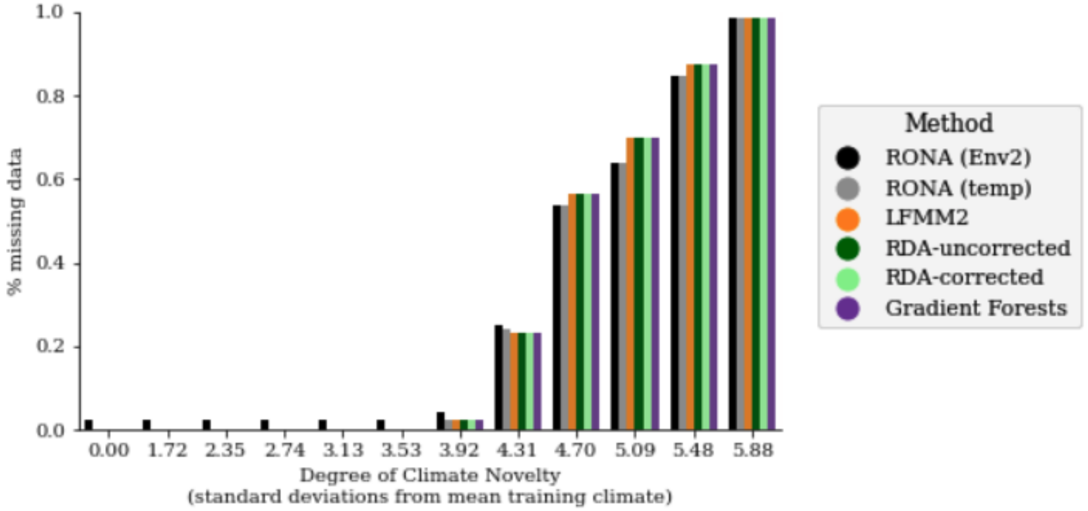

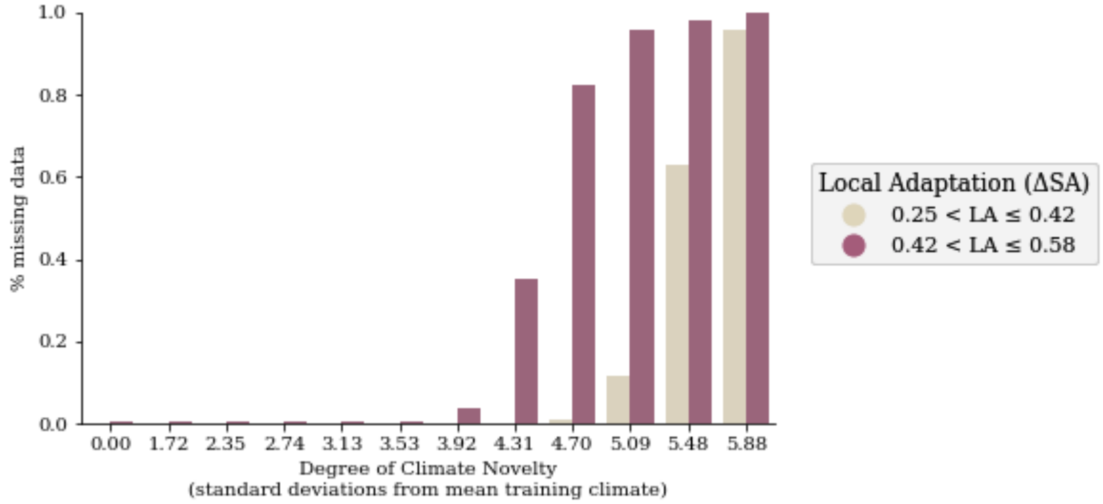

(Fig S33 continued)

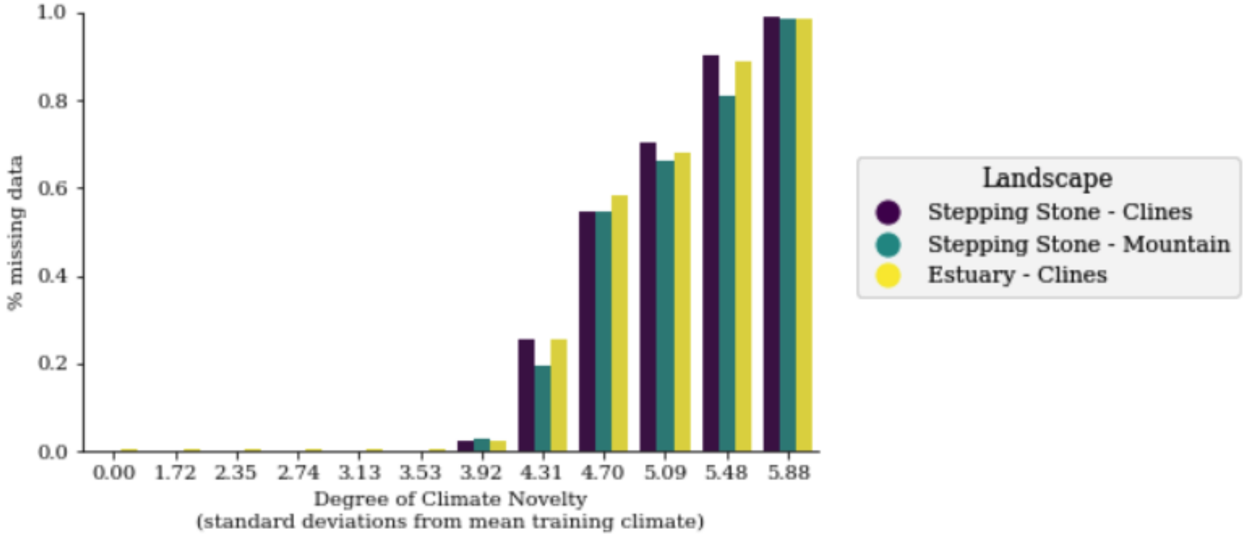

**Fig S33** The effect of simulation parameters on missing data for *Climate Novelty* scenarios. Shown are the percent missing data (y-axes) due to experimental and simulation parameters (legends). Missing data is when all populations have zero fitness in a given novelty scenario, and thus performance cannot be defined (though we manually set it to zero for other figures). Data included in these figures are from 1- and 2-trait evaluations of *Climate Novelty* scenarios. Code to create these figures can be found in SC 02.04.05.

### Supplemental Tables

**RONA-sal_opt**

| | sum_sq | df | F | PR(>F) | perc_sum_sq |
|---|---|---|---|---|---|
| glevel | 15.864775 | 2.0 | 210.709735 | 3.953193e-92 | 0.17 |
| landscape | 149.664742 | 2.0 | 1987.788561 | 0.000000e+00 | 1.59 |
| demography | 433.301763 | 4.0 | 2877.472267 | 0.000000e+00 | 4.60 |
| plevel_pleio | 0.033619 | 1.0 | 0.893039 | 3.446564e-01 | 0.00 |
| C(garden) | 1941.690095 | 99.0 | 520.985219 | 0.000000e+00 | 20.61 |
| cor_TPR_temp | 0.724989 | 1.0 | 19.258049 | 1.142528e-05 | 0.01 |
| cor_TPR_sal | 3.874653 | 1.0 | 102.923263 | 3.536502e-24 | 0.04 |
| cor_FPR_temp_neutSNPs | 0.897974 | 1.0 | 23.853067 | 1.040651e-06 | 0.01 |
| cor_FPR_sal_neutSNPs | 14.829886 | 1.0 | 393.929492 | 1.433616e-87 | 0.16 |
| final_LA | 87.744615 | 1.0 | 2330.779320 | 0.000000e+00 | 0.93 |
| Residual | 6771.995848 | 179886.0 | NaN | NaN | 71.88 |

**RONA-temp_opt**

| | sum_sq | df | F | PR(>F) | perc_sum_sq |
|---|---|---|---|---|---|
| glevel | 3.264399 | 2.0 | 99.857527 | 4.534018e-44 | 0.03 |
| landscape | 74.659979 | 2.0 | 2283.838967 | 0.000000e+00 | 0.64 |
| demography | 26.679201 | 4.0 | 408.056627 | 0.000000e+00 | 0.23 |
| plevel_pleio | 1.505757 | 1.0 | 92.121798 | 8.249029e-22 | 0.01 |
| C(garden) | 4962.472456 | 99.0 | 3066.694571 | 0.000000e+00 | 42.52 |
| cor_TPR_temp | 1.035753 | 1.0 | 63.367084 | 1.725510e-15 | 0.01 |
| cor_TPR_sal | 0.021308 | 1.0 | 1.303608 | 2.535568e-01 | 0.00 |
| cor_FPR_temp_neutSNPs | 2.155381 | 1.0 | 131.865626 | 1.640714e-30 | 0.02 |
| cor_FPR_sal_neutSNPs | 0.039487 | 1.0 | 2.415797 | 1.201186e-01 | 0.00 |
| final_LA | 3659.697867 | 1.0 | 223899.355101 | 0.000000e+00 | 31.35 |
| Residual | 2940.287212 | 179886.0 | NaN | NaN | 25.19 |

**GF**

| | sum_sq | df | F | PR(>F) | perc_sum_sq |
|---|---|---|---|---|---|
| glevel | 3.109424 | 2.0 | 158.728388 | 1.336264e-69 | 0.05 |
| landscape | 344.465503 | 2.0 | 17584.110829 | 0.000000e+00 | 5.24 |
| demography | 104.816048 | 4.0 | 2675.299841 | 0.000000e+00 | 1.59 |
| plevel_pleio | 0.373620 | 1.0 | 38.144788 | 6.582481e-10 | 0.01 |
| C(garden) | 392.397617 | 99.0 | 404.665183 | 0.000000e+00 | 5.97 |
| cor_TPR_temp | 3.305288 | 1.0 | 337.453511 | 2.682308e-75 | 0.05 |
| cor_TPR_sal | 0.498155 | 1.0 | 50.859112 | 9.961078e-13 | 0.01 |
| cor_FPR_temp_neutSNPs | 46.646231 | 1.0 | 4762.349106 | 0.000000e+00 | 0.71 |
| cor_FPR_sal_neutSNPs | 37.556091 | 1.0 | 3834.290889 | 0.000000e+00 | 0.57 |
| final_LA | 3881.402690 | 1.0 | 396271.989655 | 0.000000e+00 | 59.02 |
| Residual | 1761.946397 | 179886.0 | NaN | NaN | 26.79 |

**lfmm2**

| | sum_sq | df | F | PR(>F) | perc_sum_sq |
|---|---|---|---|---|---|
| glevel | 0.648503 | 2.0 | 26.071108 | 4.776417e-12 | 0.01 |
| landscape | 69.824926 | 2.0 | 2807.098902 | 0.000000e+00 | 1.34 |
| demography | 75.595816 | 4.0 | 1519.549983 | 0.000000e+00 | 1.45 |
| plevel_pleio | 0.088264 | 1.0 | 7.096738 | 7.723125e-03 | 0.00 |
| C(garden) | 391.270017 | 99.0 | 317.774174 | 0.000000e+00 | 7.50 |
| cor_TPR_temp | 3.333636 | 1.0 | 268.037430 | 3.359626e-60 | 0.06 |
| cor_TPR_sal | 1.434571 | 1.0 | 115.345148 | 6.738112e-27 | 0.03 |
| cor_FPR_temp_neutSNPs | 18.408493 | 1.0 | 1480.114981 | 1.679823e-322 | 0.35 |
| cor_FPR_sal_neutSNPs | 0.000221 | 1.0 | 0.017772 | 8.939475e-01 | 0.00 |
| final_LA | 2419.263042 | 1.0 | 194518.232194 | 0.000000e+00 | 46.37 |
| Residual | 2237.278977 | 179886.0 | NaN | NaN | 42.88 |

**rda-nocorr**

| | sum_sq | df | F | PR(>F) | perc_sum_sq |
|---|---|---|---|---|---|
| glevel | 2.722620 | 2.0 | 114.252516 | 2.583753e-50 | 0.04 |
| landscape | 351.685552 | 2.0 | 14758.193717 | 0.000000e+00 | 5.17 |
| demography | 86.983434 | 4.0 | 1825.093984 | 0.000000e+00 | 1.28 |
| plevel_pleio | 0.043391 | 1.0 | 3.641764 | 5.634878e-02 | 0.00 |
| C(garden) | 384.324179 | 99.0 | 325.815086 | 0.000000e+00 | 5.65 |
| cor_TPR_temp | 4.653284 | 1.0 | 390.542466 | 7.801493e-87 | 0.07 |
| cor_TPR_sal | 1.660051 | 1.0 | 139.325332 | 3.842628e-32 | 0.02 |
| cor_FPR_temp_neutSNPs | 57.083425 | 1.0 | 4790.917538 | 0.000000e+00 | 0.84 |
| cor_FPR_sal_neutSNPs | 39.340402 | 1.0 | 3301.774980 | 0.000000e+00 | 0.58 |
| final_LA | 3728.556367 | 1.0 | 312931.576881 | 0.000000e+00 | 54.83 |
| Residual | 2143.328255 | 179886.0 | NaN | NaN | 31.52 |

**rda-structcorr**

| | sum_sq | df | F | PR(>F) | perc_sum_sq |
|---|---|---|---|---|---|
| glevel | 19.270968 | 2.0 | 303.886012 | 1.763727e-132 | 0.29 |
| landscape | 54.926356 | 2.0 | 866.139726 | 0.000000e+00 | 0.81 |
| demography | 644.354351 | 4.0 | 5080.447174 | 0.000000e+00 | 9.54 |
| plevel_pleio | 2.784965 | 1.0 | 87.832851 | 7.201110e-21 | 0.04 |
| C(garden) | 184.307173 | 99.0 | 58.714343 | 0.000000e+00 | 2.73 |
| cor_TPR_temp | 13.822608 | 1.0 | 435.940440 | 1.078989e-96 | 0.20 |
| cor_TPR_sal | 0.004197 | 1.0 | 0.132380 | 7.159771e-01 | 0.00 |
| cor_FPR_temp_neutSNPs | 1.803296 | 1.0 | 56.872745 | 4.671127e-14 | 0.03 |
| cor_FPR_sal_neutSNPs | 45.534733 | 1.0 | 1436.084373 | 5.262511e-313 | 0.67 |
| final_LA | 85.096985 | 1.0 | 2683.807346 | 0.000000e+00 | 1.26 |
| Residual | 5703.746286 | 179886.0 | NaN | NaN | 84.43 |

**Table S1** Results from Type II ANOVAs from regressing simulation factors on offset performance (see Equation 1 of the main text). In this table, the common garden ID was included as a categorical factor (n=100 per simulation). Code to create these tables can be found in SC 02.02.01.

**RONA (Env2)**

| | sum_sq | df | F | PR(>F) | perc_sum_sq |
|---|---|---|---|---|---|
| $p_{cQTN,\,temp}$ | 11.076279 | 1.0 | 203.959434 | 3.027995e-46 | 0.11 |
| $p_{cQTN,\,Env2}$ | 7.130868 | 1.0 | 131.308340 | 2.171984e-30 | 0.07 |
| $p_{cNeut,\,temp}$ | 350.186388 | 1.0 | 6448.358576 | 0.000000e+00 | 3.40 |
| $p_{cNeut,\,Env2}$ | 165.769820 | 1.0 | 3052.497975 | 0.000000e+00 | 1.61 |
| **Residual** | 9774.859470 | 179995.0 | NaN | NaN | 94.82 |

**RONA (temp)**

| | sum_sq | df | F | PR(>F) | perc_sum_sq |
|---|---|---|---|---|---|
| $p_{cQTN,\,temp}$ | 461.435097 | 1.0 | 6854.545092 | 0.000000e+00 | 3.54 |
| $p_{cQTN,\,Env2}$ | 321.669143 | 1.0 | 4778.344045 | 0.000000e+00 | 2.47 |
| $p_{cNeut,\,temp}$ | 21.729532 | 1.0 | 322.788745 | 4.136405e-72 | 0.17 |
| $p_{cNeut,\,Env2}$ | 119.950781 | 1.0 | 1781.849799 | 0.000000e+00 | 0.92 |
| **Residual** | 12116.925221 | 179995.0 | NaN | NaN | 92.91 |

**RDA-uncorrected**

| | sum_sq | df | F | PR(>F) | perc_sum_sq |
|---|---|---|---|---|---|
| $p_{cQTN,\,temp}$ | 326.478563 | 1.0 | 8168.479995 | 0.0 | 3.29 |
| $p_{cQTN,\,Env2}$ | 171.117641 | 1.0 | 4281.356235 | 0.0 | 1.72 |
| $p_{cNeut,\,temp}$ | 589.314088 | 1.0 | 14744.613849 | 0.0 | 5.93 |
| $p_{cNeut,\,Env2}$ | 1649.284886 | 1.0 | 41265.038898 | 0.0 | 16.61 |
| **Residual** | 7194.056785 | 179995.0 | NaN | NaN | 72.45 |

**RDA-corrected**

| | sum_sq | df | F | PR(>F) | perc_sum_sq |
|---|---|---|---|---|---|
| $p_{cQTN,\,temp}$ | 36.345449 | 1.0 | 970.987128 | 1.344472e-212 | 0.48 |
| $p_{cQTN,\,Env2}$ | 26.680765 | 1.0 | 712.790185 | 1.002191e-156 | 0.35 |
| $p_{cNeut,\,temp}$ | 281.267846 | 1.0 | 7514.213388 | 0.000000e+00 | 3.70 |
| $p_{cNeut,\,Env2}$ | 512.076667 | 1.0 | 13680.388309 | 0.000000e+00 | 6.74 |
| **Residual** | 6737.472479 | 179995.0 | NaN | NaN | 88.72 |

**Gradient Forests**

| | sum_sq | df | F | PR(>F) | perc_sum_sq |
|---|---|---|---|---|---|
| $p_{cQTN,\,temp}$ | 350.833368 | 1.0 | 9184.348121 | 0.0 | 3.69 |
| $p_{cQTN,\,Env2}$ | 210.540691 | 1.0 | 5511.673549 | 0.0 | 2.21 |
| $p_{cNeut,\,temp}$ | 375.213154 | 1.0 | 9822.578289 | 0.0 | 3.94 |
| $p_{cNeut,\,Env2}$ | 1702.524610 | 1.0 | 44569.816059 | 0.0 | 17.89 |
| **Residual** | 6875.637914 | 179995.0 | NaN | NaN | 72.26 |

**LFMM2**

| | sum_sq | df | F | PR(>F) | perc_sum_sq |
|---|---|---|---|---|---|
| $p_{cQTN,\,temp}$ | 256.979852 | 1.0 | 8357.260709 | 0.0 | 3.72 |
| $p_{cQTN,\,Env2}$ | 173.645491 | 1.0 | 5647.137802 | 0.0 | 2.51 |
| $p_{cNeut,\,temp}$ | 718.499204 | 1.0 | 23366.365591 | 0.0 | 10.40 |
| $p_{cNeut,\,Env2}$ | 227.994551 | 1.0 | 7414.627593 | 0.0 | 3.30 |
| **Residual** | 5534.718859 | 179995.0 | NaN | NaN | 80.08 |

**Table S2** Results from Type II ANOVAs from regressing the proportion of clinal QTNs (cor_TPR_tmp and cor_TPR_sal) and clinal neutral alleles (cor_FPR_temp_neutSNPs, cor_FPR_sal_neutSNPs) on offset performance (see Equation 2 of the main text). Code to create these tables can be found in SC 02.02.05.

**RONA-sal_opt**

| | sum_sq | df | F | PR(>F) | perc_sum_sq |
|---|---|---|---|---|---|
| **all** | 13.066056 | 1.0 | 939.445081 | 2.543708e-166 | 34.04 |
| **final_LA** | 0.321152 | 1.0 | 23.090730 | 1.673759e-06 | 0.84 |
| **Residual** | 24.993162 | 1797.0 | NaN | NaN | 65.12 |

**RONA-temp_opt**

| | sum_sq | df | F | PR(>F) | perc_sum_sq |
|---|---|---|---|---|---|
| **all** | 0.086501 | 1.0 | 8.324025 | 0.003959 | 0.18 |
| **final_LA** | 30.251825 | 1.0 | 2911.137332 | 0.000000 | 61.72 |
| **Residual** | 18.673983 | 1797.0 | NaN | NaN | 38.10 |

**lfmm2**

| | sum_sq | df | F | PR(>F) | perc_sum_sq |
|---|---|---|---|---|---|
| **all** | 3.673454 | 1.0 | 873.091702 | 9.874623e-157 | 8.64 |
| **final_LA** | 31.290136 | 1.0 | 7436.913517 | 0.000000e+00 | 73.58 |
| **Residual** | 7.560714 | 1797.0 | NaN | NaN | 17.78 |

**GF**

| | sum_sq | df | F | PR(>F) | perc_sum_sq |
|---|---|---|---|---|---|
| **all** | 8.505224 | 1.0 | 749.172746 | 3.638160e-138 | 10.54 |
| **final_LA** | 51.787680 | 1.0 | 4561.657568 | 0.000000e+00 | 64.18 |
| **Residual** | 20.401019 | 1797.0 | NaN | NaN | 25.28 |

**rda-nocorr**

| | sum_sq | df | F | PR(>F) | perc_sum_sq |
|---|---|---|---|---|---|
| **all** | 10.763364 | 1.0 | 827.118396 | 6.020460e-150 | 12.54 |
| **final_LA** | 51.668736 | 1.0 | 3970.521018 | 0.000000e+00 | 60.21 |
| **Residual** | 23.384518 | 1797.0 | NaN | NaN | 27.25 |

**rda-structcorr**

| | sum_sq | df | F | PR(>F) | perc_sum_sq |
|---|---|---|---|---|---|
| **all** | 3.192007 | 1.0 | 78.914320 | 1.537690e-18 | 4.20 |
| **final_LA** | 0.062105 | 1.0 | 1.535395 | 2.154664e-01 | 0.08 |
| **Residual** | 72.686890 | 1797.0 | NaN | NaN | 95.71 |

**Table S3** Results from Type II ANOVAs regressing two factors - degree of local adaptation (final_LA) and levels of isolation-by-environment in *all* marker sets) on offset performance. Code to create these tables can be found in SC 02.02.11.

| Nuisance Level | *Adaptive* models | *All* models | *Neutral* models |
|---|---|---|---|
| 1-trait 1-nuisance | 45/45 | 45/45 | 45/45 |
| 1-trait 3-nuisance | 45/45 | 43/45 | 38/45 |
| 1-trait 4-nuisance | 43/45 | 36/45 | 35/45 |
| 2-trait 2-nuisance | 120/180 | 119/180 | 119/180 |
| 2-trait 3-nuisance | 140/180 | 119/180 | 119/180 |

**Table S4** Gradient Forests (GF) sometimes incorrectly identifies the environments driving adaptation. Shown are the proportions of simulation levels (*N* 1-trait = 45 levels; *N* 2-trait = 180 levels; one replicate each) where weighted feature importance output from GF correctly identified the adaptive environments in the top-most ranks. If at least one nuisance environment was ranked above an adaptive environment this was counted as incorrect. Data used to create this table is from the GF models output from the *Nuisance Environment* workflow. Code used to create this table can be found in SC 02.10.02.

### Supplemental Figures

Figs. S1-S3 are in Supplemental Note S1.

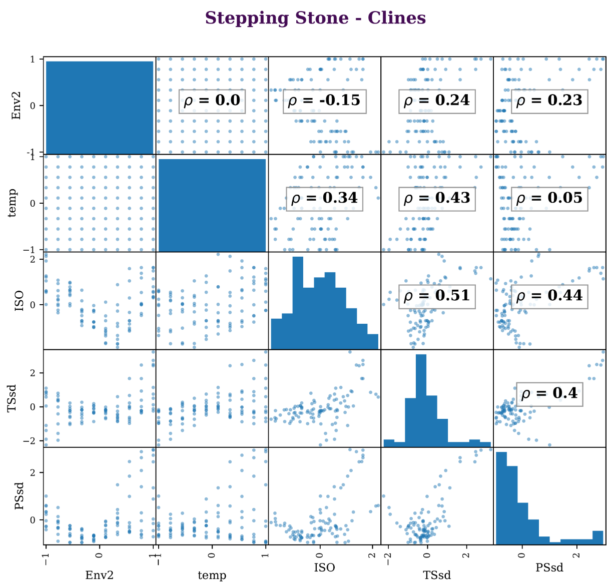

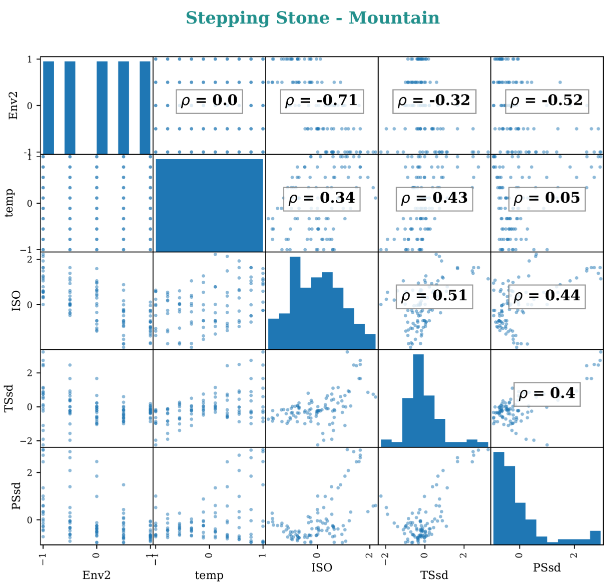

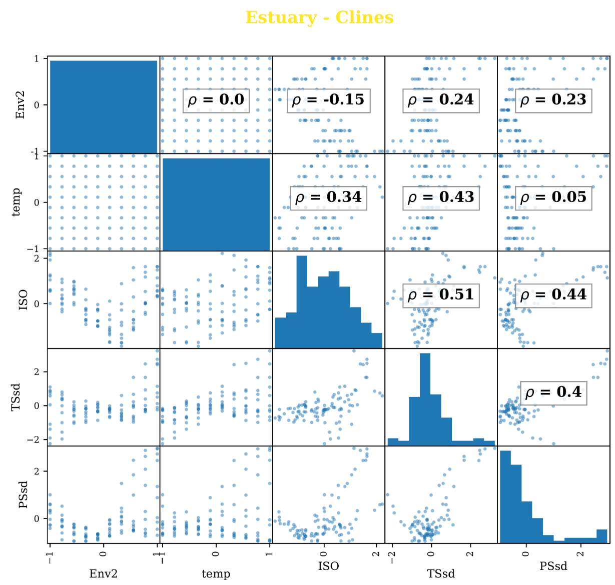

**Fig S4** Correlation (Spearman's rho) among environmental variables faceted by landscape. On-diagonal entries are histograms of environmental values, off-diagonal entries are scatter plots between pairwise variables. Included are environmental variables from 1-trait (*temp*), 2-trait (*temp*, *Env2*), and 6-trait simulations (MAT, MTWetQ, MTDQ, PDM, PwarmQ, PWM), as well as nuisance environmental variables (ISO, PSsd, TSsd). Note *Estuary - Clines* and *Stepping Stone - Clines* have the same correlation structure; *Stepping Stones - Mountain* only differs from these two landscapes with *Env2*. Figure continues on the next page. Code to create these figures can be found in SC 02.07.02.11.

(Fig S4 continued)

6-trait

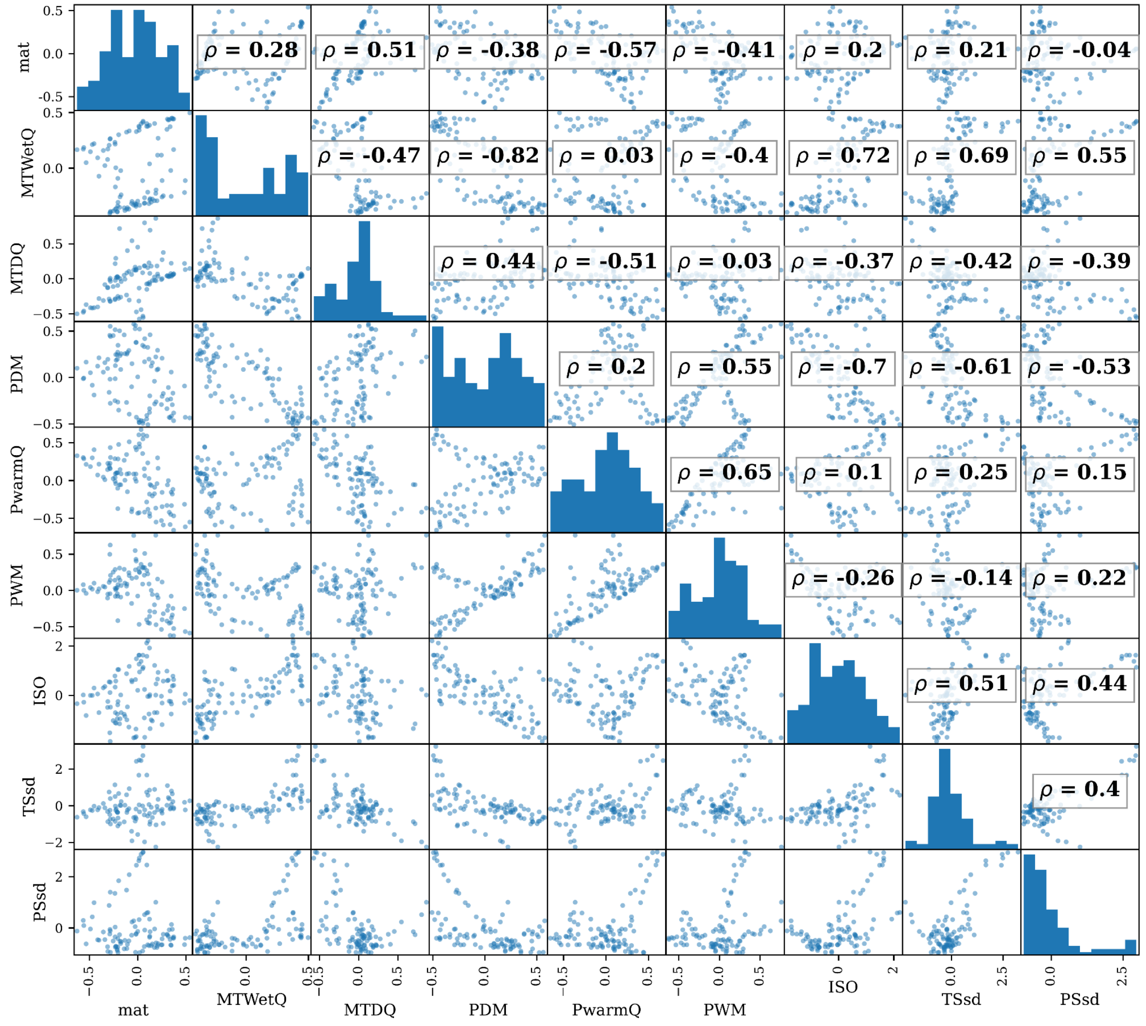

Fig S5 is in Supplemental Note S3

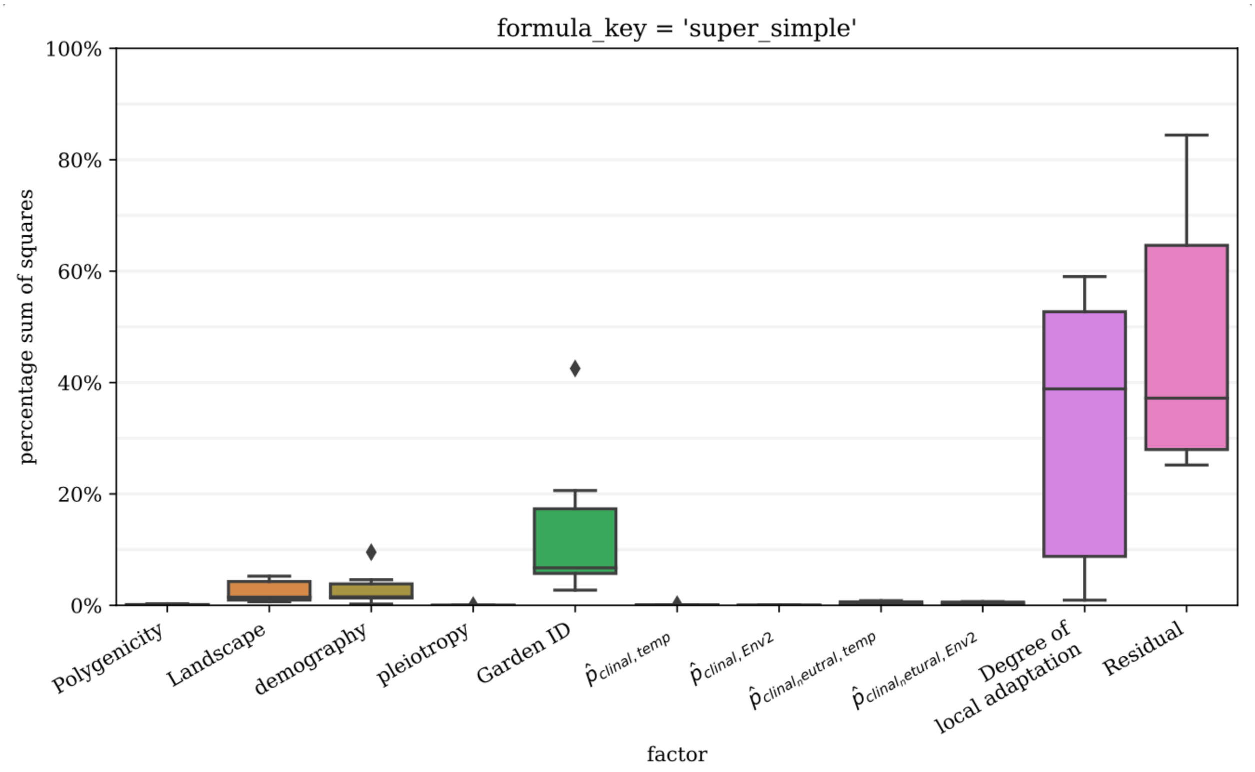

**Fig S6** Percent sum of squares of the various factors from the ANOVA model in Table S1. Boxplots are created from the percent sum of squares from each method's individual ANOVA model. Data included in this figure are from models trained using all markers and simulations with two selective environments with performance evaluated in all 100 common gardens. Code to create this table is in 02.02.01.

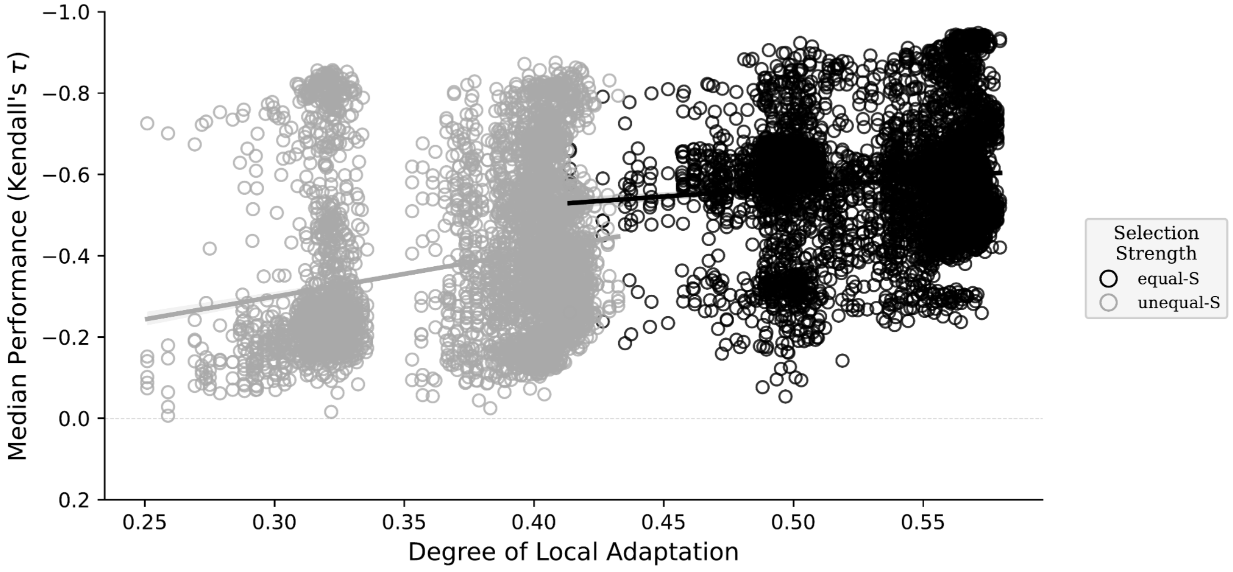

**Fig S7** Effect of the degree of local adaptation (x-axes) on method performance (y-axes) colored by the relative strength of selection on the two traits. Shown are the linear relationships between the median validation scores (circles, taken from validation scores across all 100 common gardens on the landscape) and the simulation's mean level of local adaptation (taken across all 100 populations). Data included in this figure are from models trained using all markers and simulations with two selective environments. Code to create this figure can be found in SC 02.02.02.

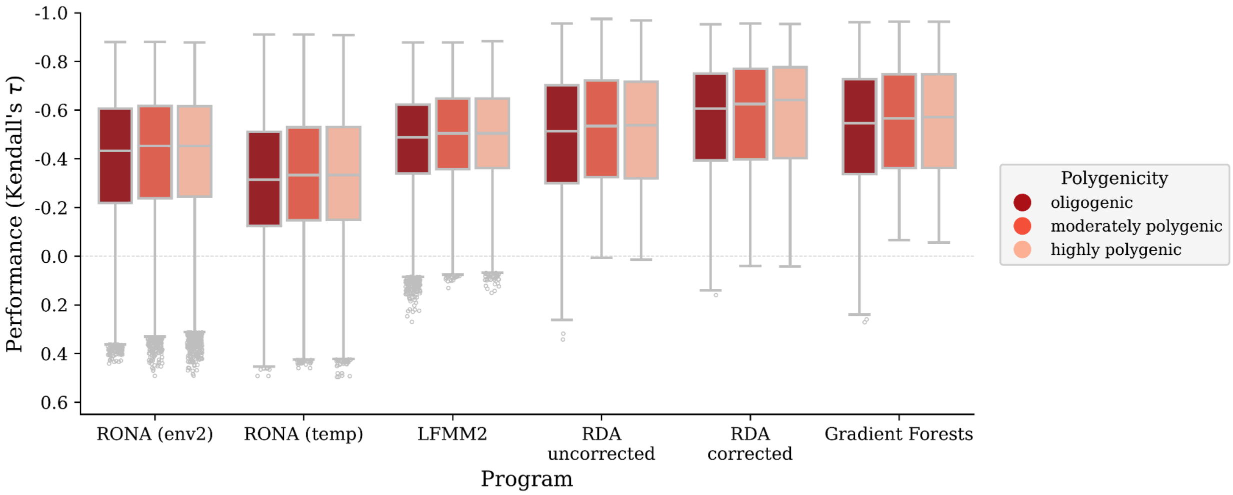

**Fig S8** Effect of polygenicity on performance of offset methods trained using all markers on simulations with two adaptive traits. Code to create this figure can be found in SC 02.02.01.

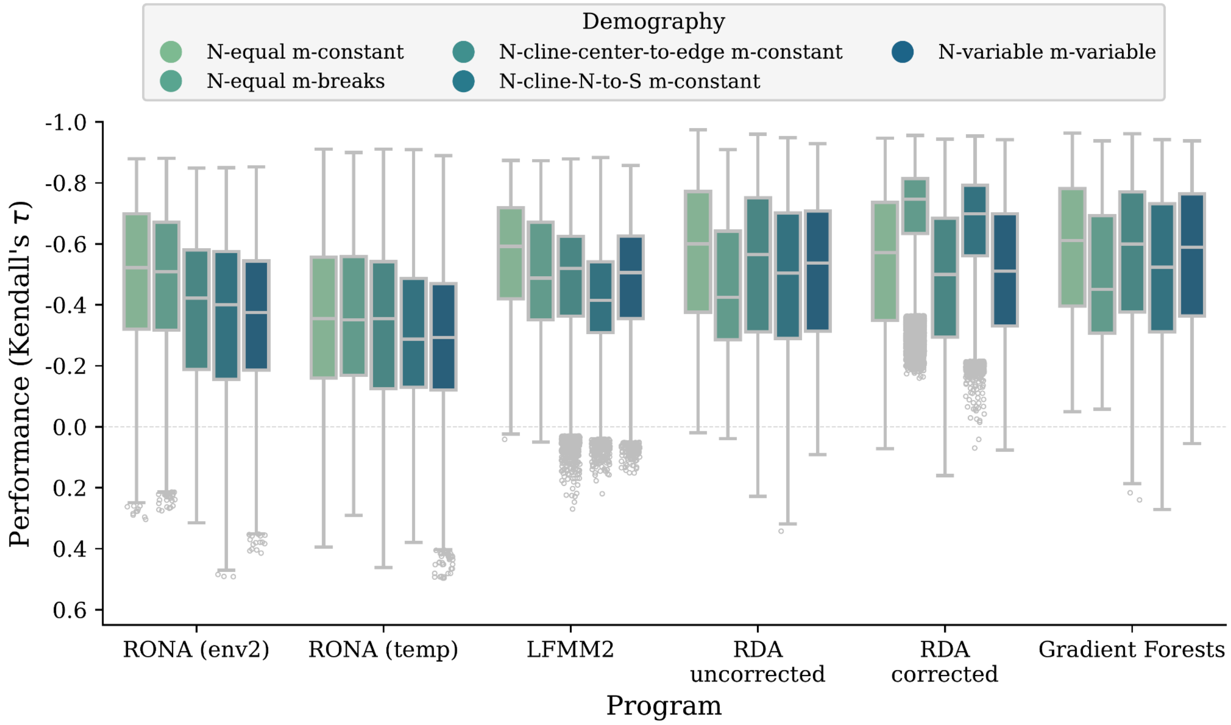

**Fig S9** Effect of demography on performance of offset methods trained using all markers on simulations with two adaptive traits. Code to create this figure can be found in SC 02.02.01.

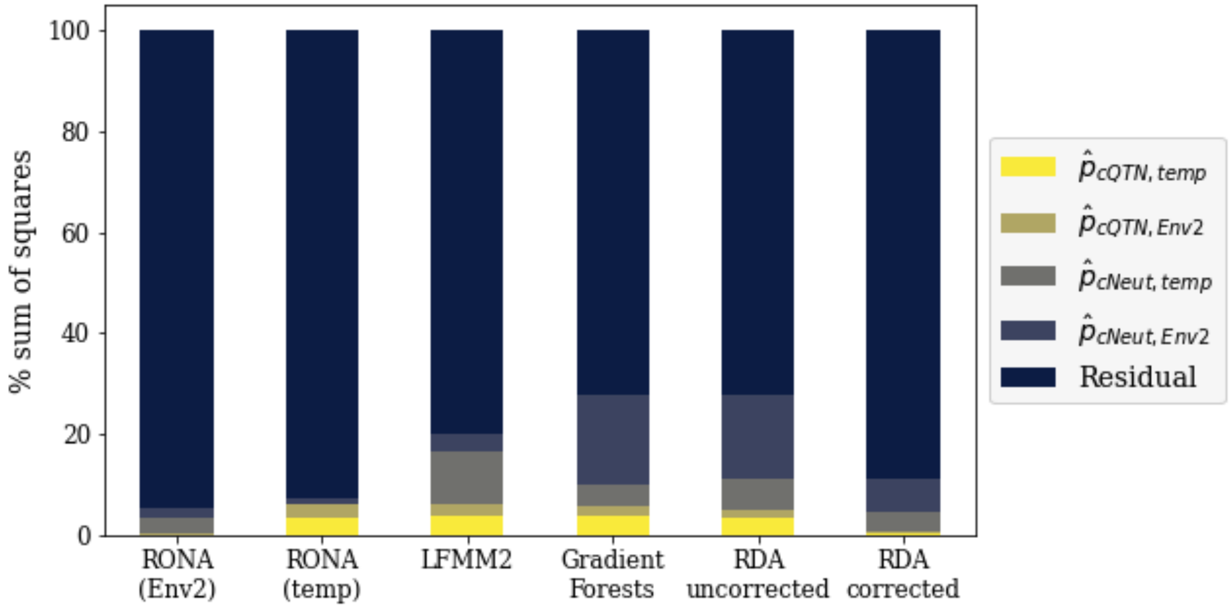

**Fig S10** Stacked bar plot of the percent sum of squares from Type II ANOVAs from regressing the proportion of clinal QTNs and clinal neutral alleles on offset performance (see Equation 2 of the main text). Code to create this table is in 02.02.05.

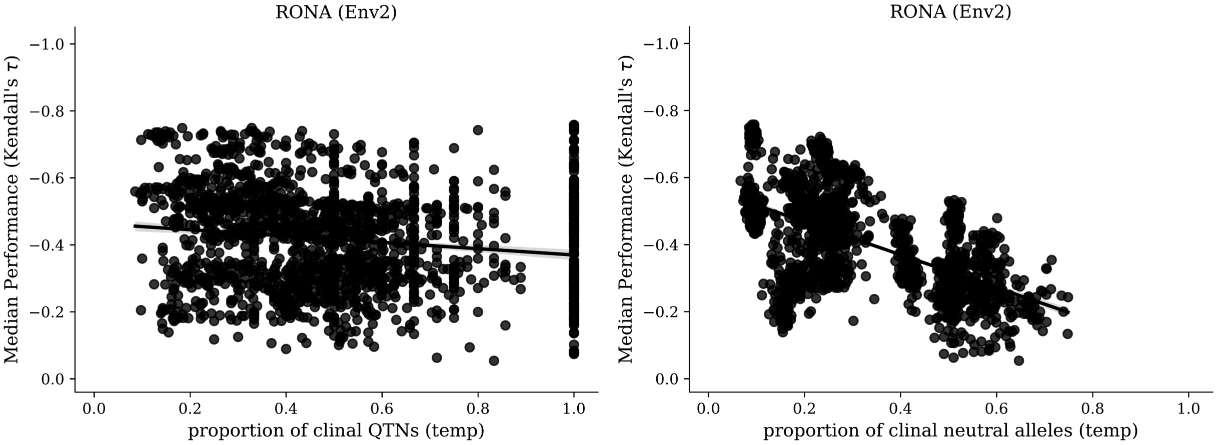
RONA (Env2)
Median Performance (Kendall's τ)
proportion of clinal QTNs (temp)
RONA (Env2)
Median Performance (Kendall's τ)
proportion of clinal neutral alleles (temp)

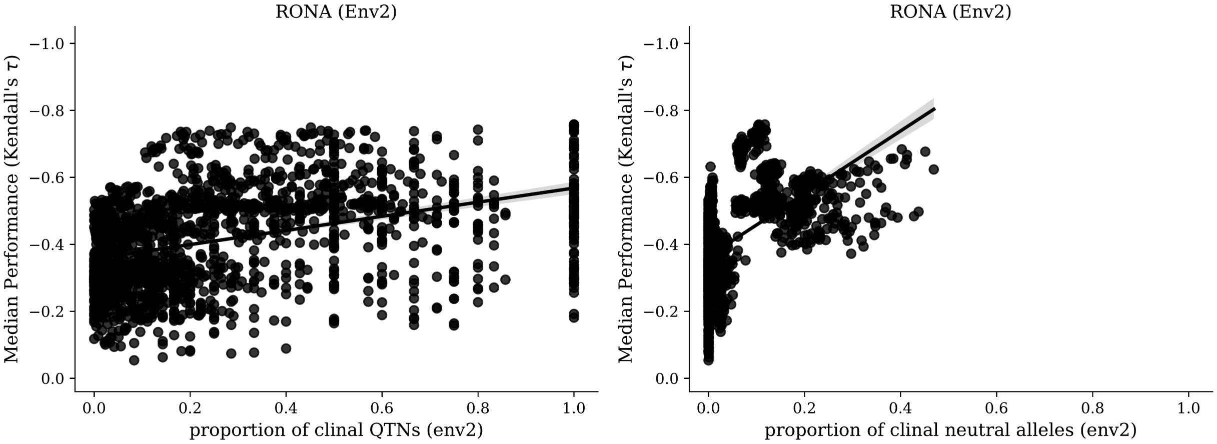
RONA (Env2)
Median Performance (Kendall's τ)
proportion of clinal QTNs (env2)
RONA (Env2)
Median Performance (Kendall's τ)
proportion of clinal neutral alleles (env2)

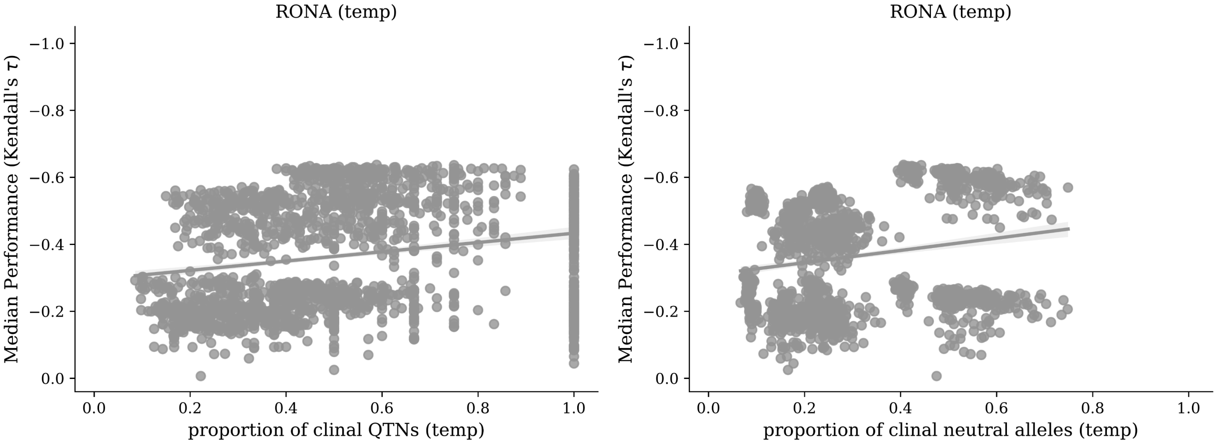
RONA (temp)
Median Performance (Kendall's τ)
proportion of clinal QTNs (temp)
RONA (temp)
Median Performance (Kendall's τ)
proportion of clinal neutral alleles (temp)

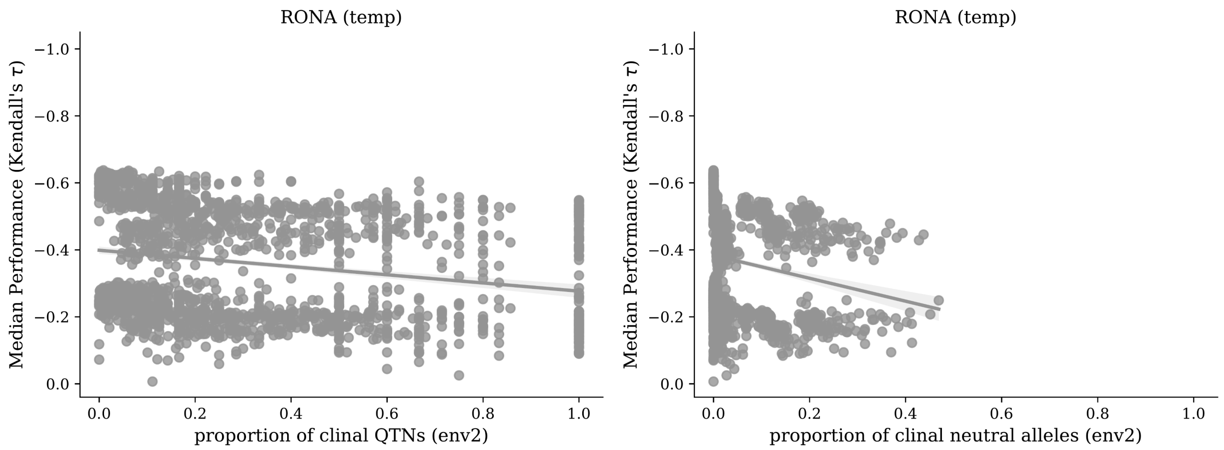
RONA (temp)
Median Performance (Kendall's τ)
−1.0
−0.8
−0.6
−0.4
−0.2
0.0
0.0
0.2
0.4
0.6
0.8
1.0
proportion of clinal QTNs (env2)
RONA (temp)
Median Performance (Kendall's τ)
proportion of clinal neutral alleles (env2)

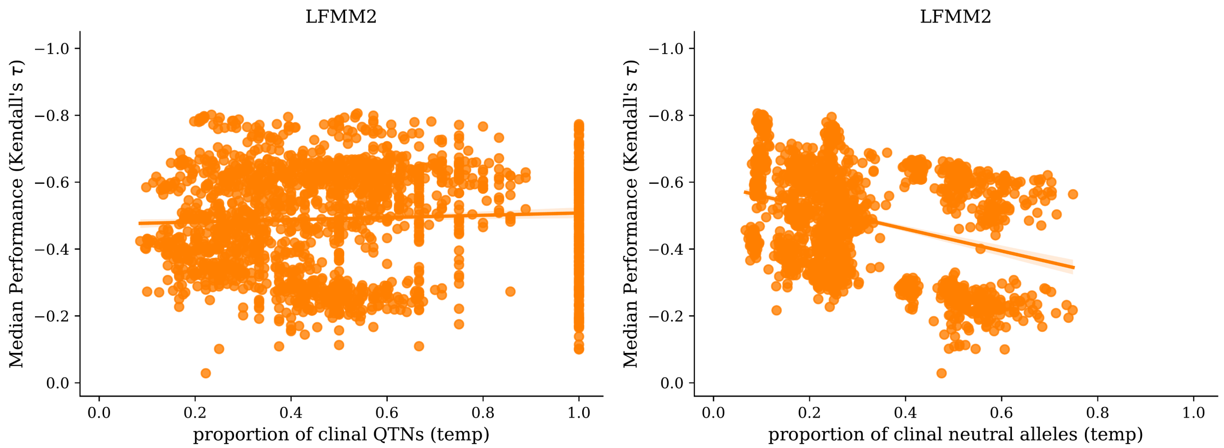
LFMM2
Median Performance (Kendall's τ)
proportion of clinal QTNs (temp)
LFMM2
Median Performance (Kendall's τ)
proportion of clinal neutral alleles (temp)

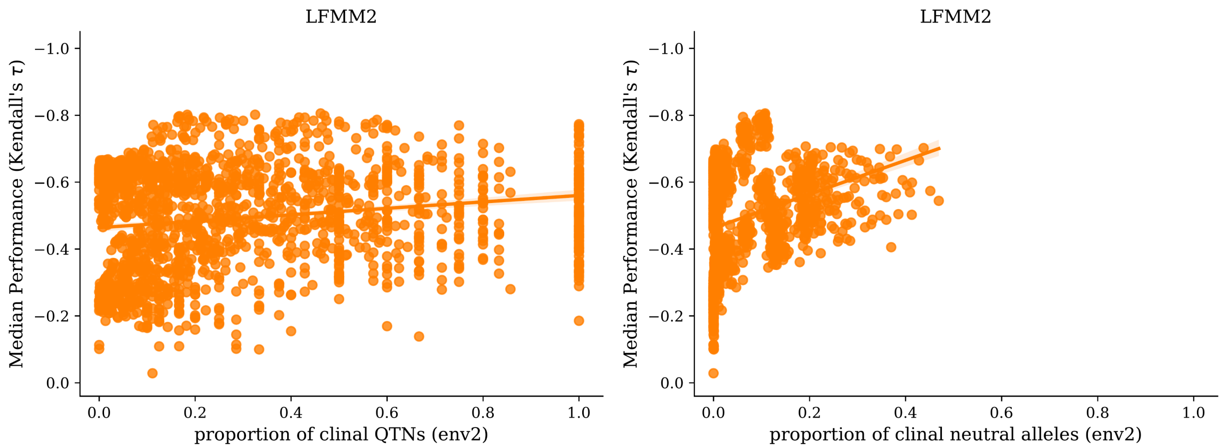
LFMM2
Median Performance (Kendall's τ)
proportion of clinal QTNs (env2)
LFMM2
Median Performance (Kendall's τ)
proportion of clinal neutral alleles (env2)

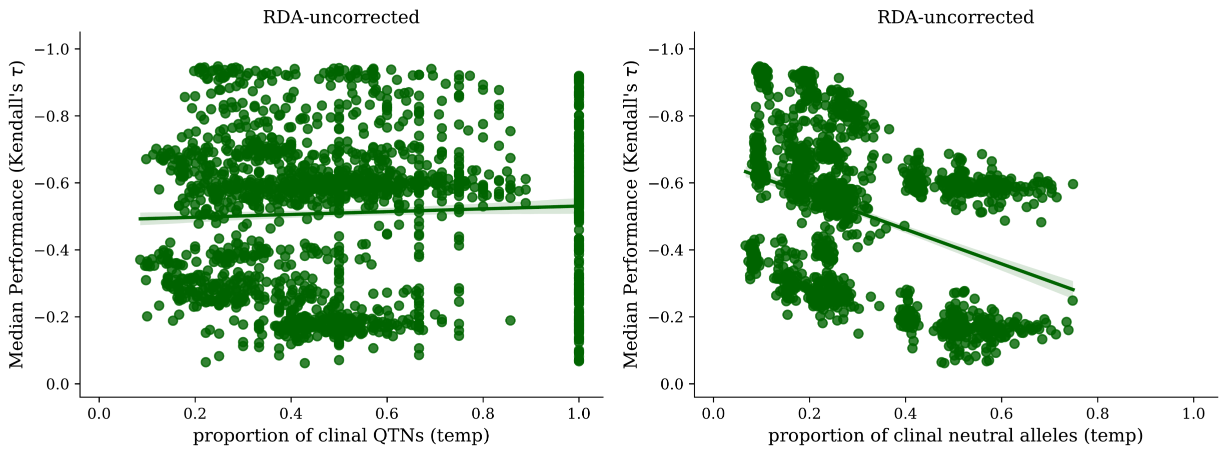
RDA-uncorrected
Median Performance (Kendall's τ)
proportion of clinal QTNs (temp)
RDA-uncorrected
Median Performance (Kendall's τ)
proportion of clinal neutral alleles (temp)

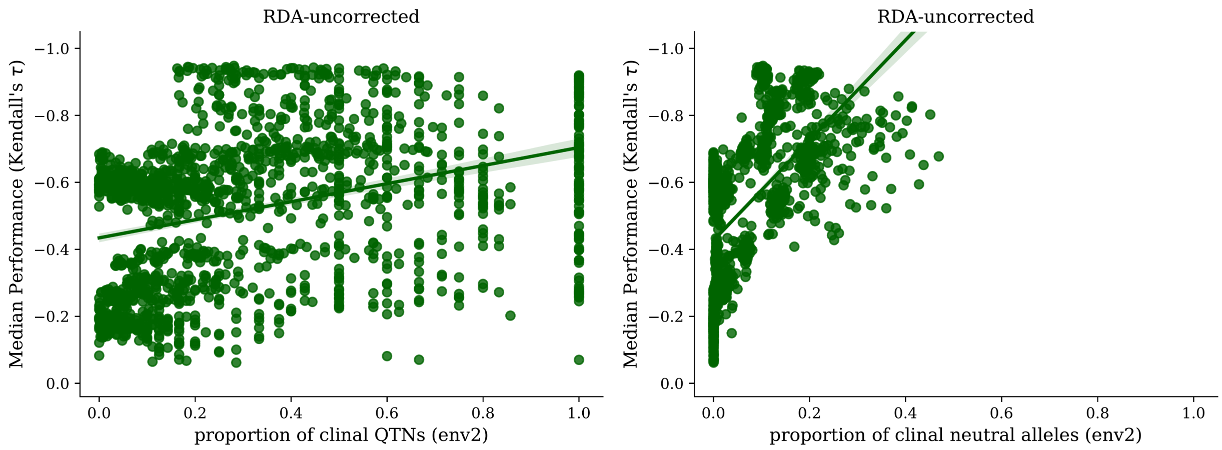
RDA-uncorrected
Median Performance (Kendall's τ)
proportion of clinal QTNs (env2)
RDA-uncorrected
Median Performance (Kendall's τ)
proportion of clinal neutral alleles (env2)

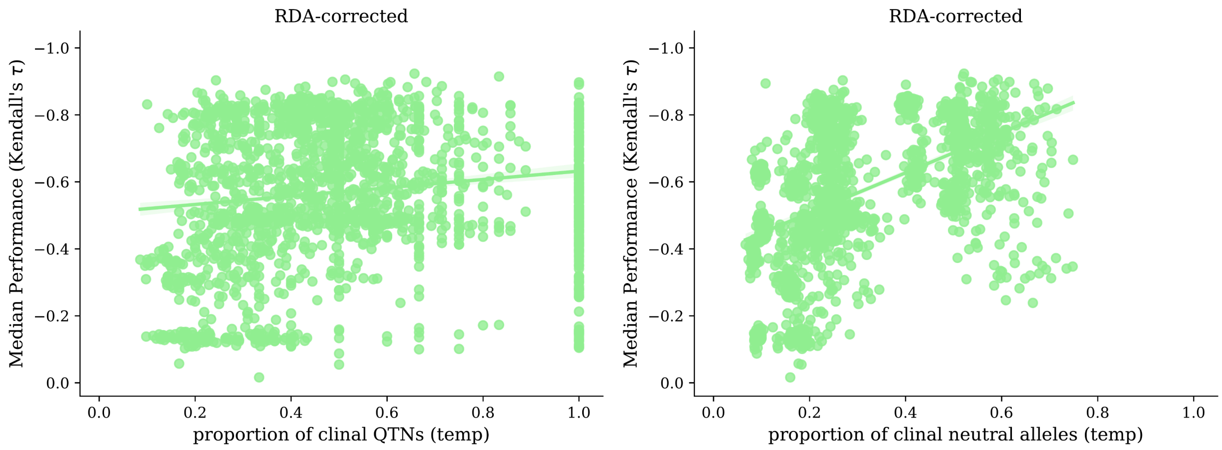
RDA-corrected
Median Performance (Kendall's τ)
proportion of clinal QTNs (temp)
RDA-corrected
Median Performance (Kendall's τ)
proportion of clinal neutral alleles (temp)

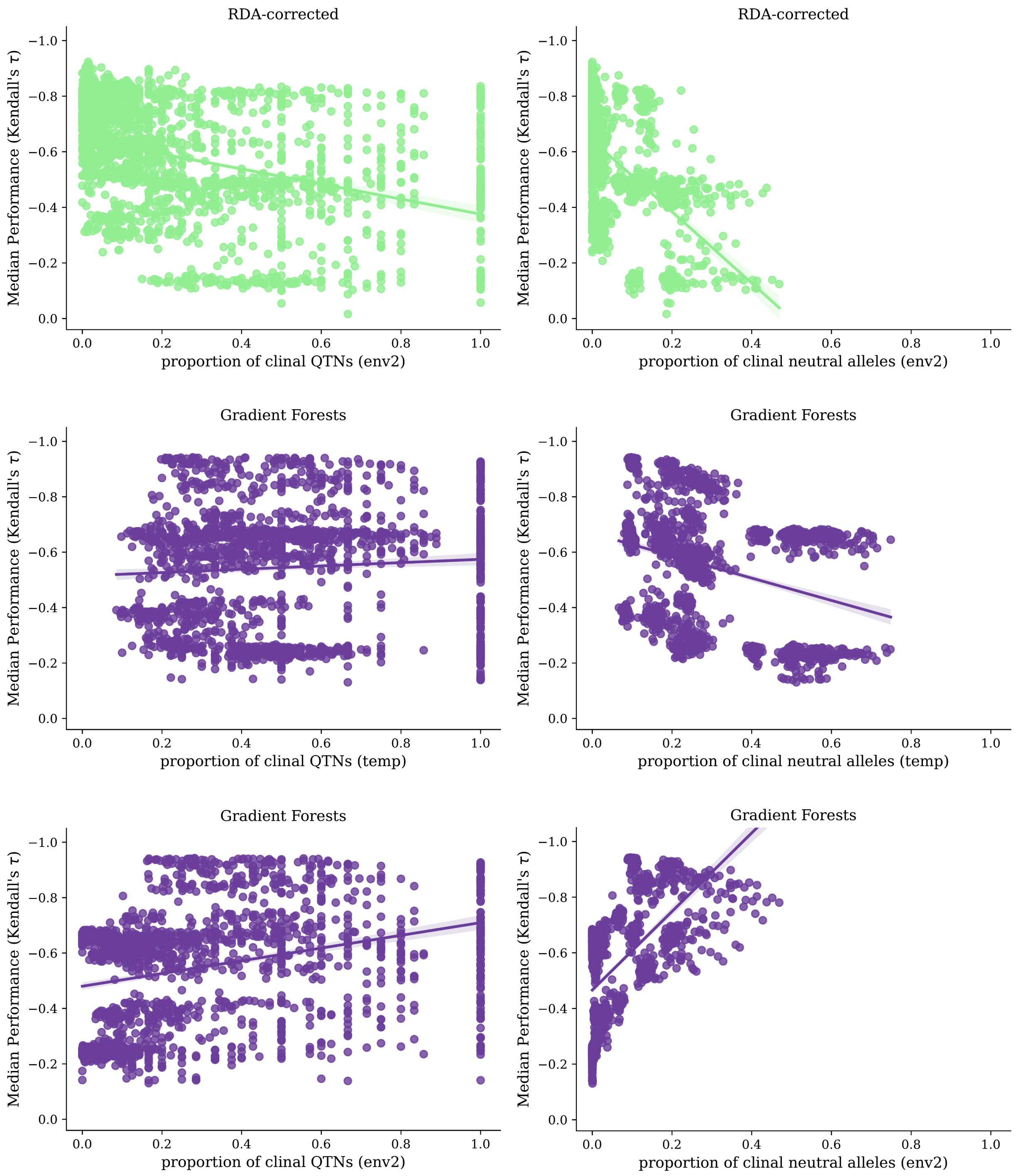

**Fig S11** Impact on method performance (y-axes) from the proportion of QTNs with clinal relationships with *temp* (first column) or *Env2* (second column). Model performance is quantified as Kendall's rank correlation between offset and fitness; shown are median values from scores from 10 replicates per seed (100 common gardens for each replicate). Data included in this figure is from evaluation of 2-trait simulations using *all* markers. Code to create these figures can be found in 02.01.03.

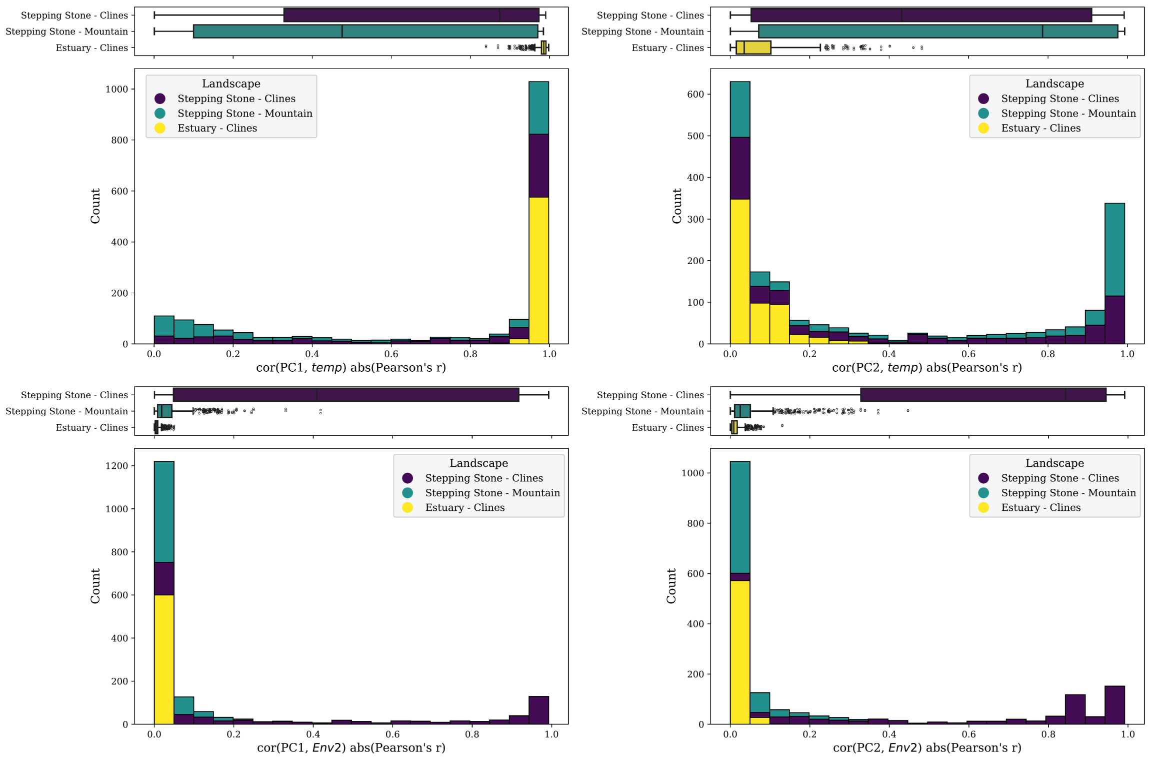

**Fig S12** Stacked bar plot showing correlation between environmental variables (rows) and axes of population genetic structure (Principal Component Analysis axes [PC axes]; columns). Data included in this figure is from all 2-trait simulations. Code to create this figure can be found in SC 02.10.03.

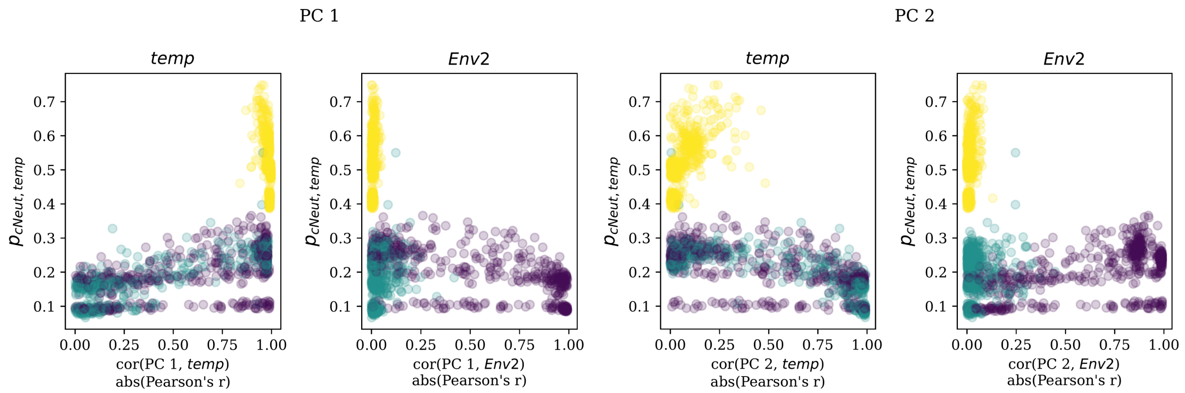

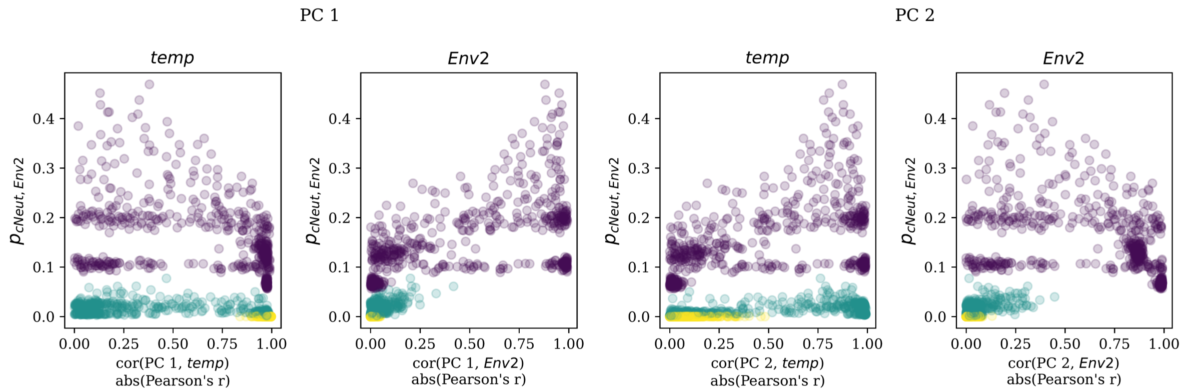

**Fig S13** Relationship between the proportion of clinal neutral loci for *temp* (y-axes, first row) or *Env2* (y-axes, second row) with the strength of the relationship between environmental variables and axes of population genetic structure. Purple = *Stepping Stone - Clines*; teal = *Stepping Stone - Clines*; yellow = *Estuary - Clines*. Data included in this figure is from all 2-trait simulations. Code to create this figure can be found in 02.10.03.

(Fig S14)

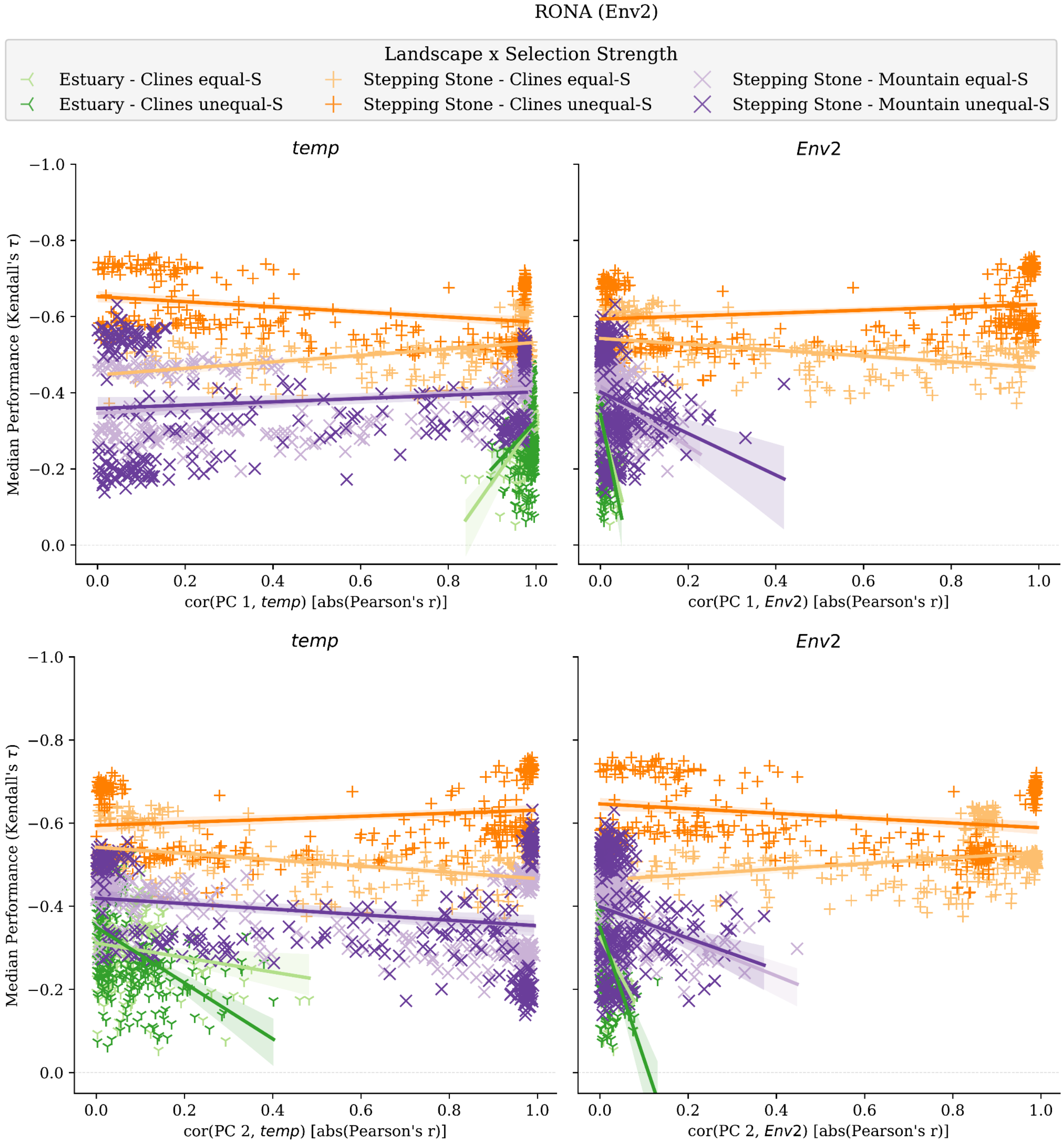
RONA (Env2)
Landscape x Selection Strength
Estuary - Clines equal-S
Estuary - Clines unequal-S
Stepping Stone - Clines equal-S
Stepping Stone - Clines unequal-S
Stepping Stone - Mountain equal-S
Stepping Stone - Mountain unequal-S
temp
Env2
Median Performance (Kendall's τ)
−1.0
−0.8
−0.6
−0.4
−0.2
0.0
0.0
0.2
0.4
0.6
0.8
1.0
cor(PC 1, temp) [abs(Pearson's r)]
cor(PC 1, Env2) [abs(Pearson's r)]
cor(PC 2, temp) [abs(Pearson's r)]
cor(PC 2, Env2) [abs(Pearson's r)]

(Fig S14 continued)

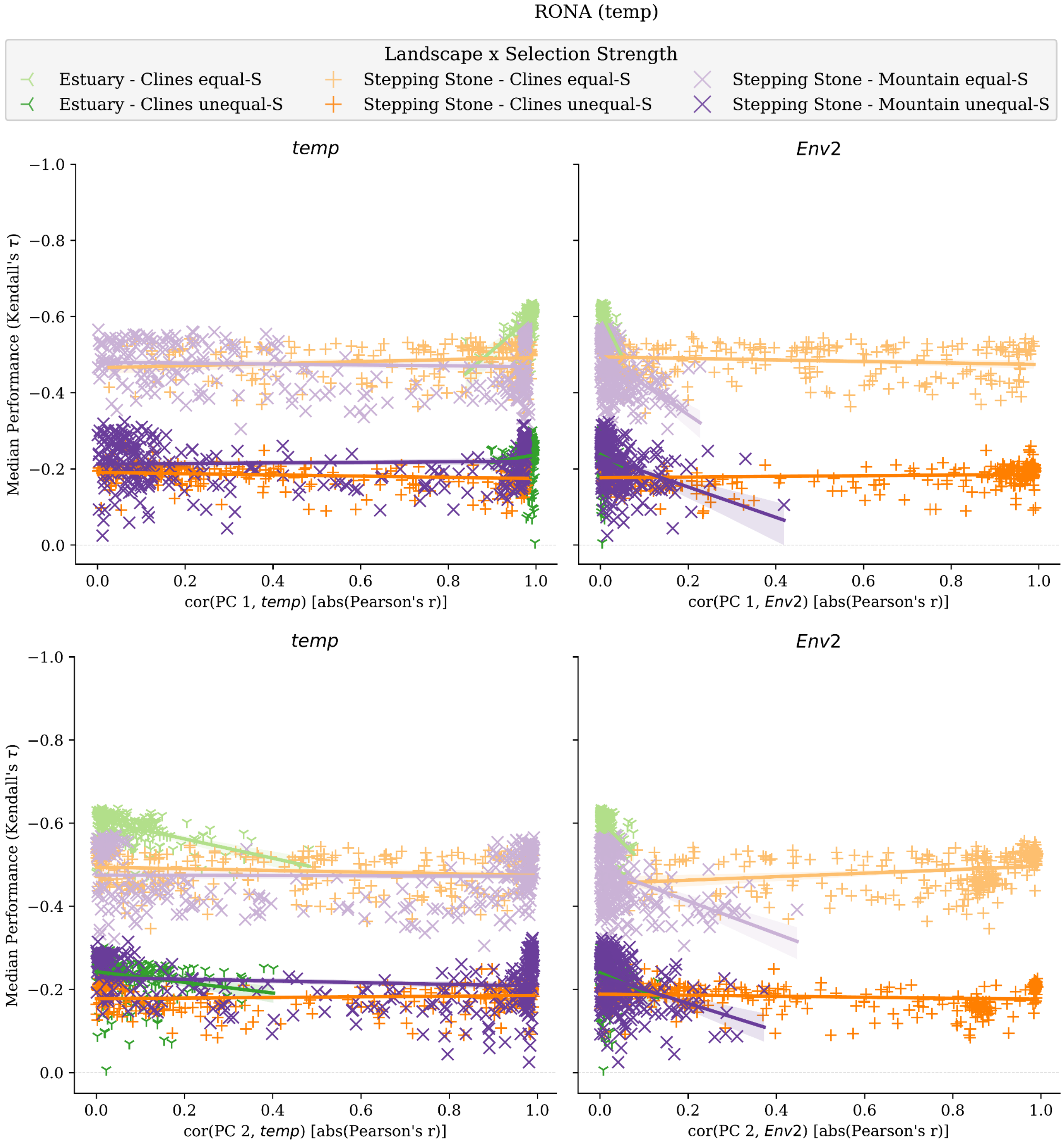

(Fig S14 continued)

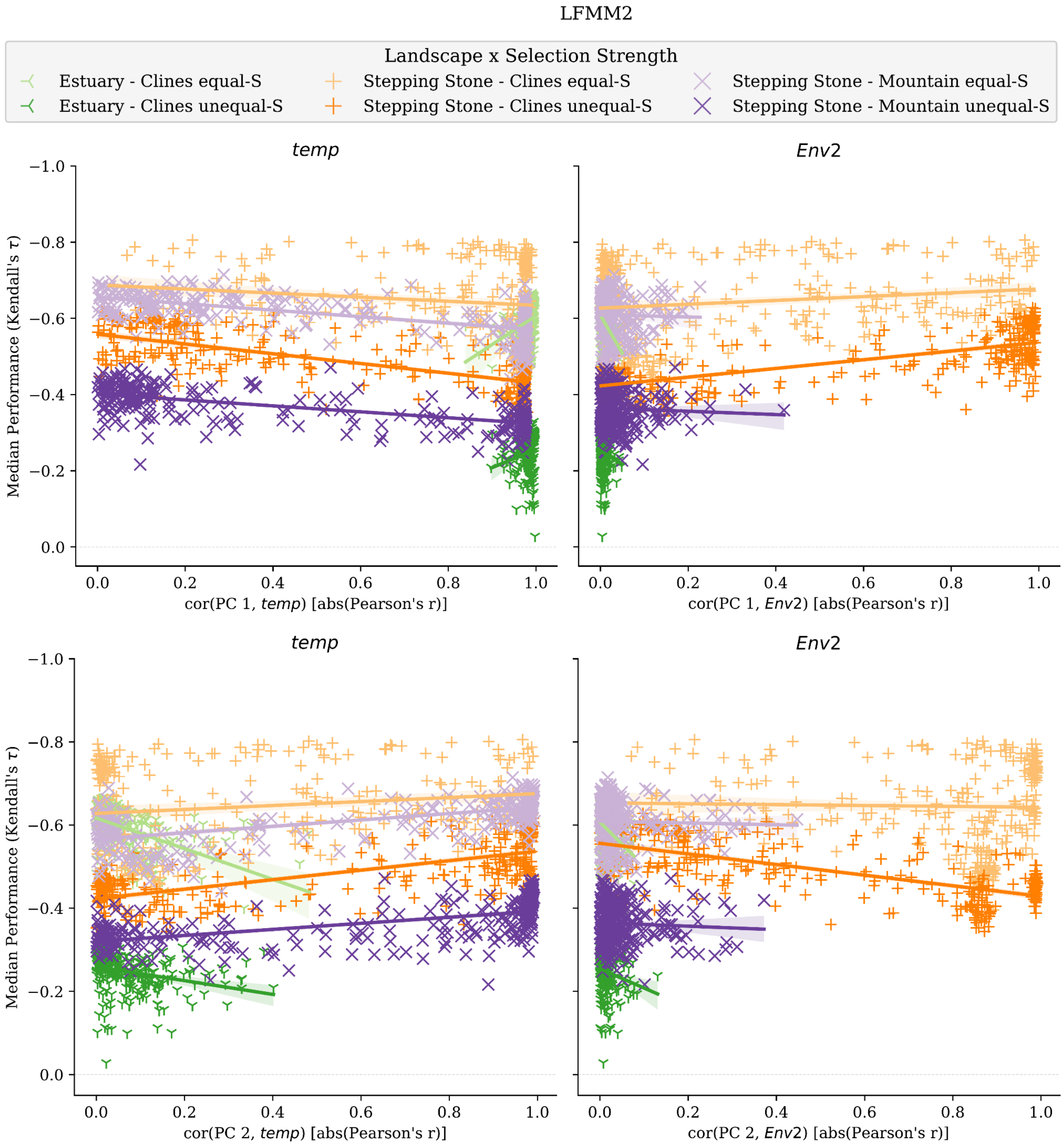

(Fig S14 continued)

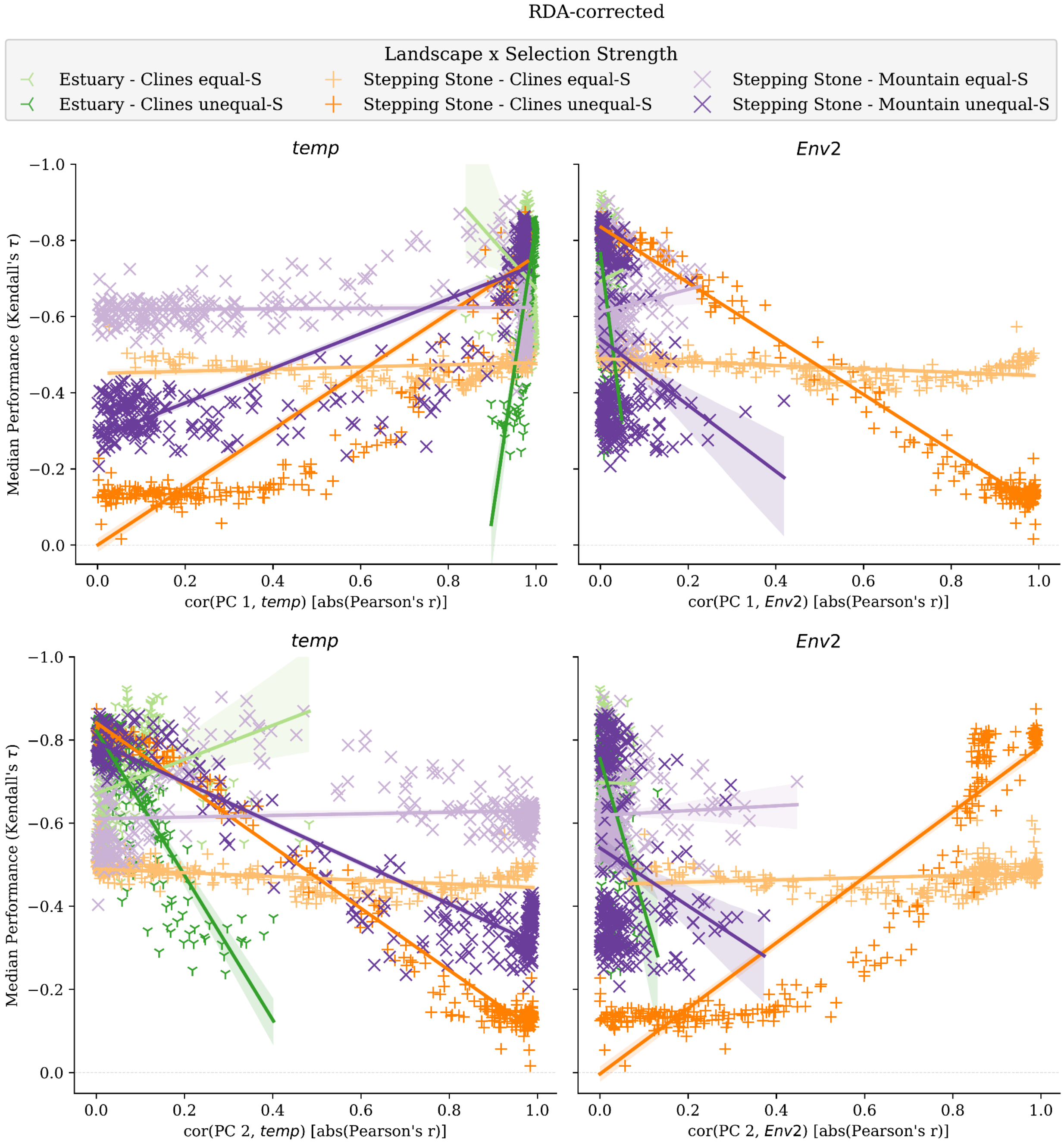

(Fig S14 continued)

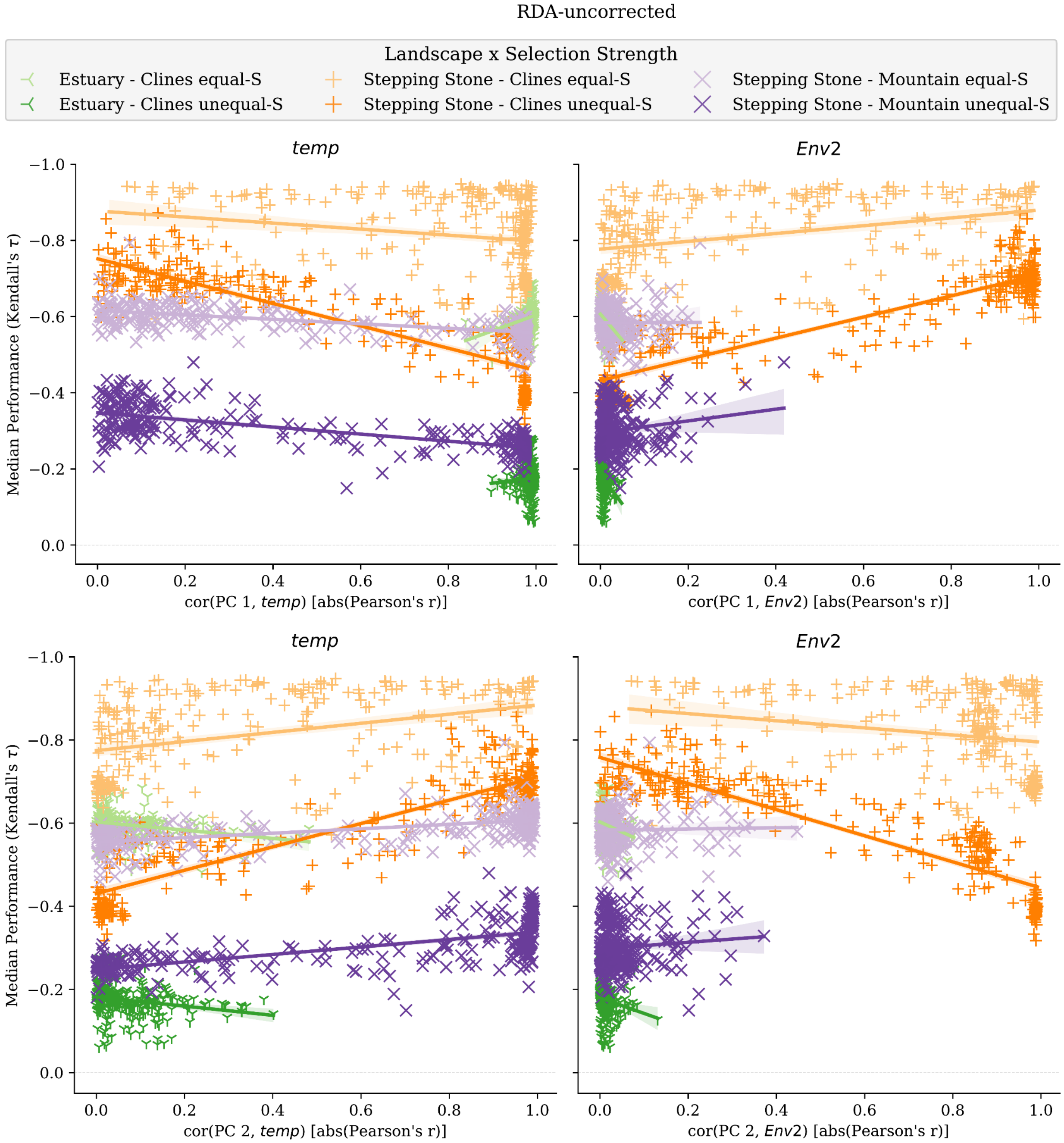

(Fig S14 continued)

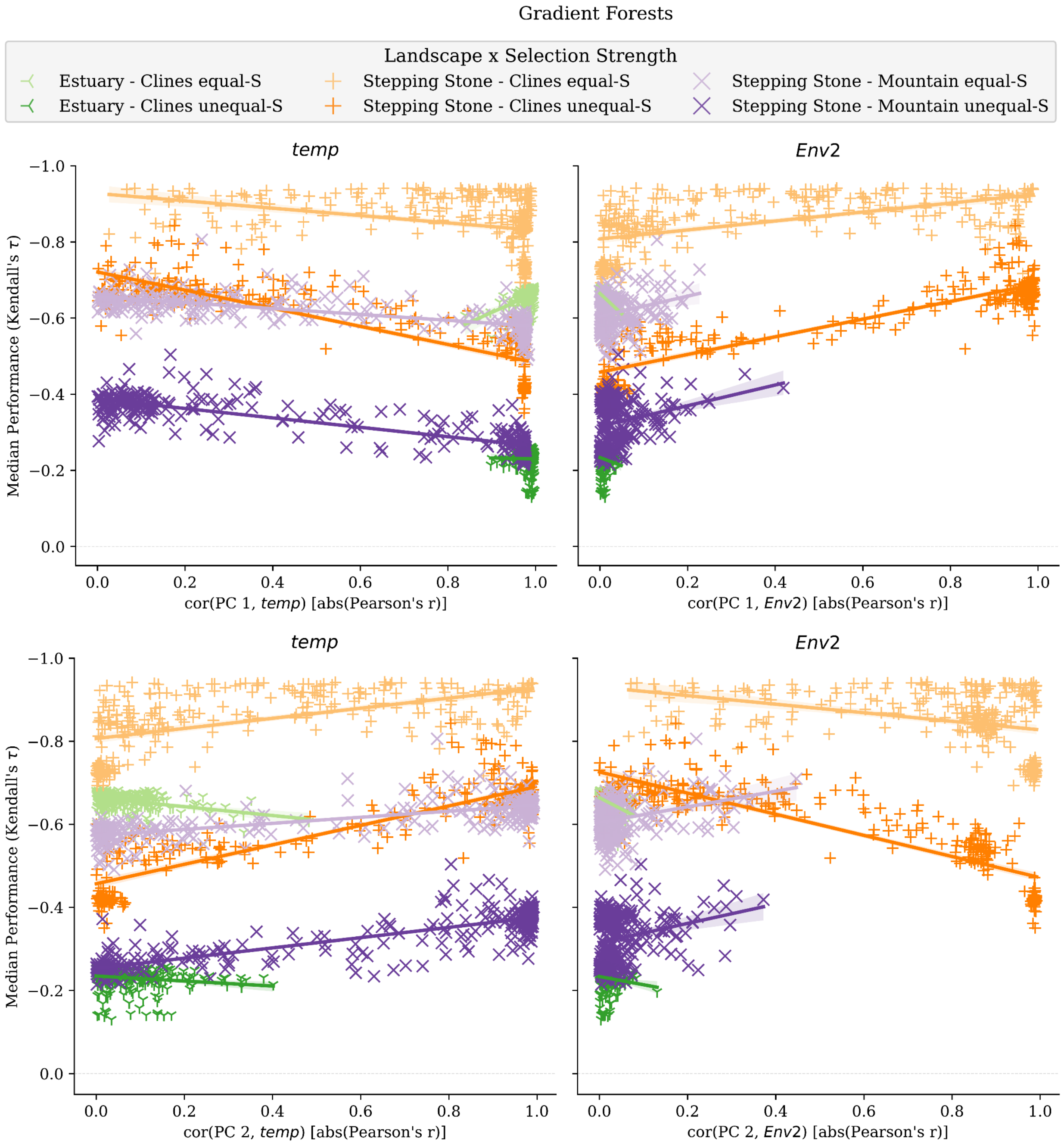

**Fig S14** Relationship between median performance and absolute correlation (Pearson's r) between environmental variables and axes of population genetic structure (principal component analysis axes). Each subfigure is for a different method (see panel titles). Data used in this figure is from 2-trait simulations. Code to create this figure can be found in SC 02.10.03.

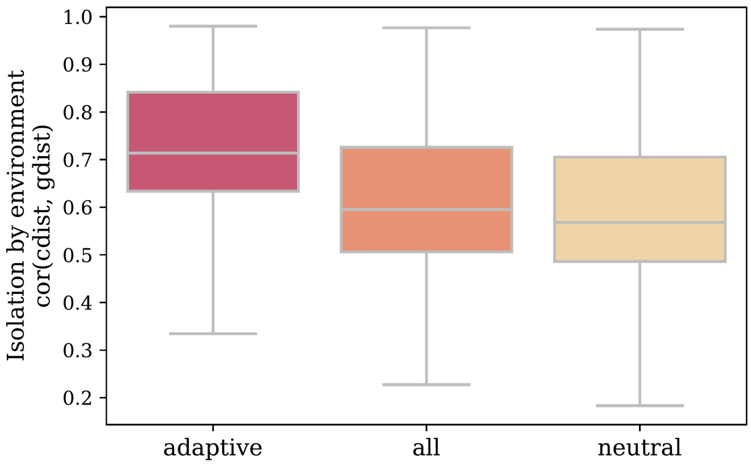

**Fig S15** *Adaptive* markers contain greater levels of isolation-by-environment (IBE) than other marker sets. *IBE* is quantified as Spearman's rank correlation between population pairwise $F_{ST}$ and Euclidean distance of adaptive environments. Code to create this figure can be found in SC 02.02.10.

(Fig S16)

A) *adaptive* markers

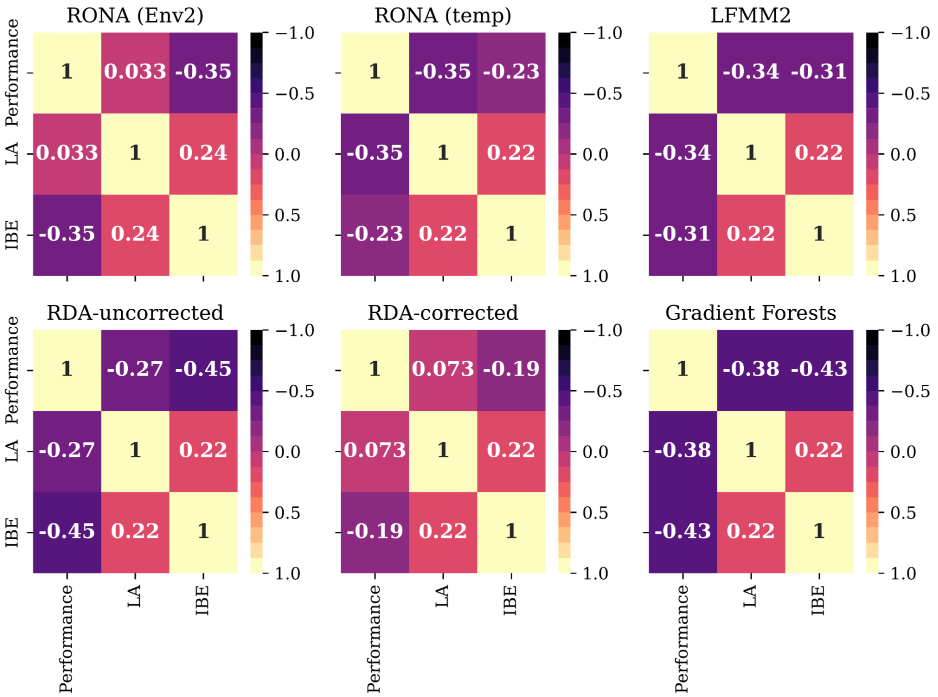

*B) all* markers

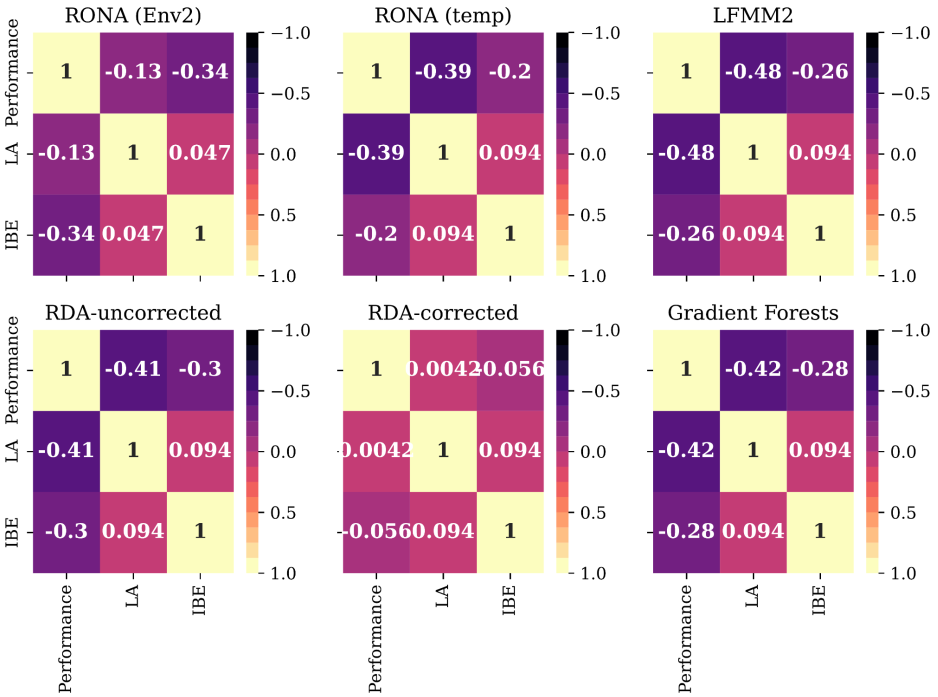

*C) neutral* markers

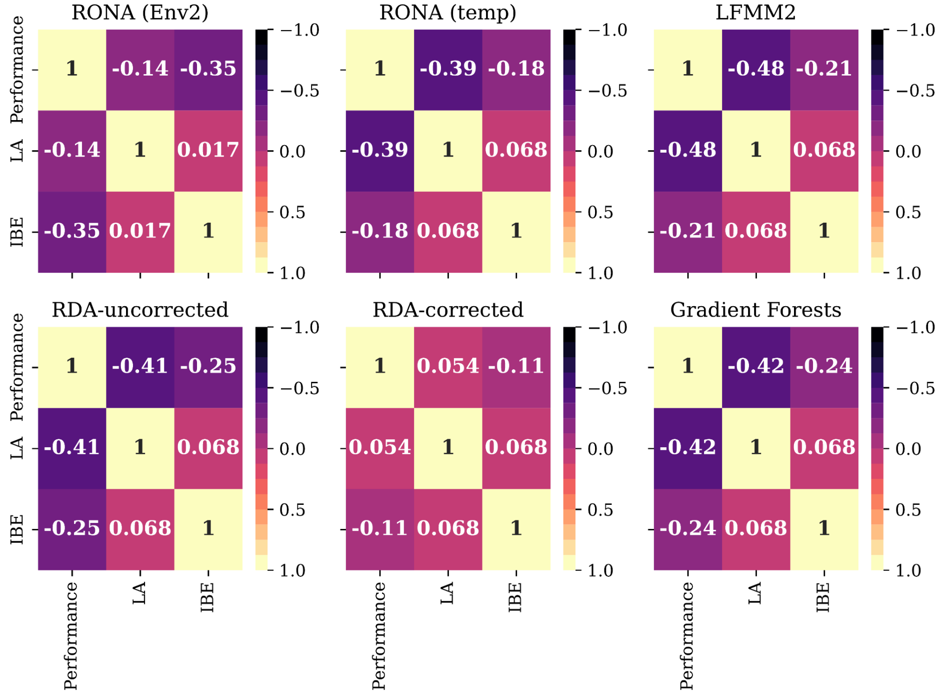

**Fig S16** The relationship between the degree of local adaptation ($LA_{\Delta SA}$), levels of *IBE* within marker sets, and median performance of models trained with one of the three marker sets: (A) *adaptive*, (B) *all*, and (c) *neutral* marker sets. *IBE* is quantified as Spearman's rank correlation between population pairwise $F_{ST}$ and Euclidean distance of adaptive environments. Data included in these figures are from 1- and 2-trait simulations. Code to create these figures can be found in SC 02.02.10.

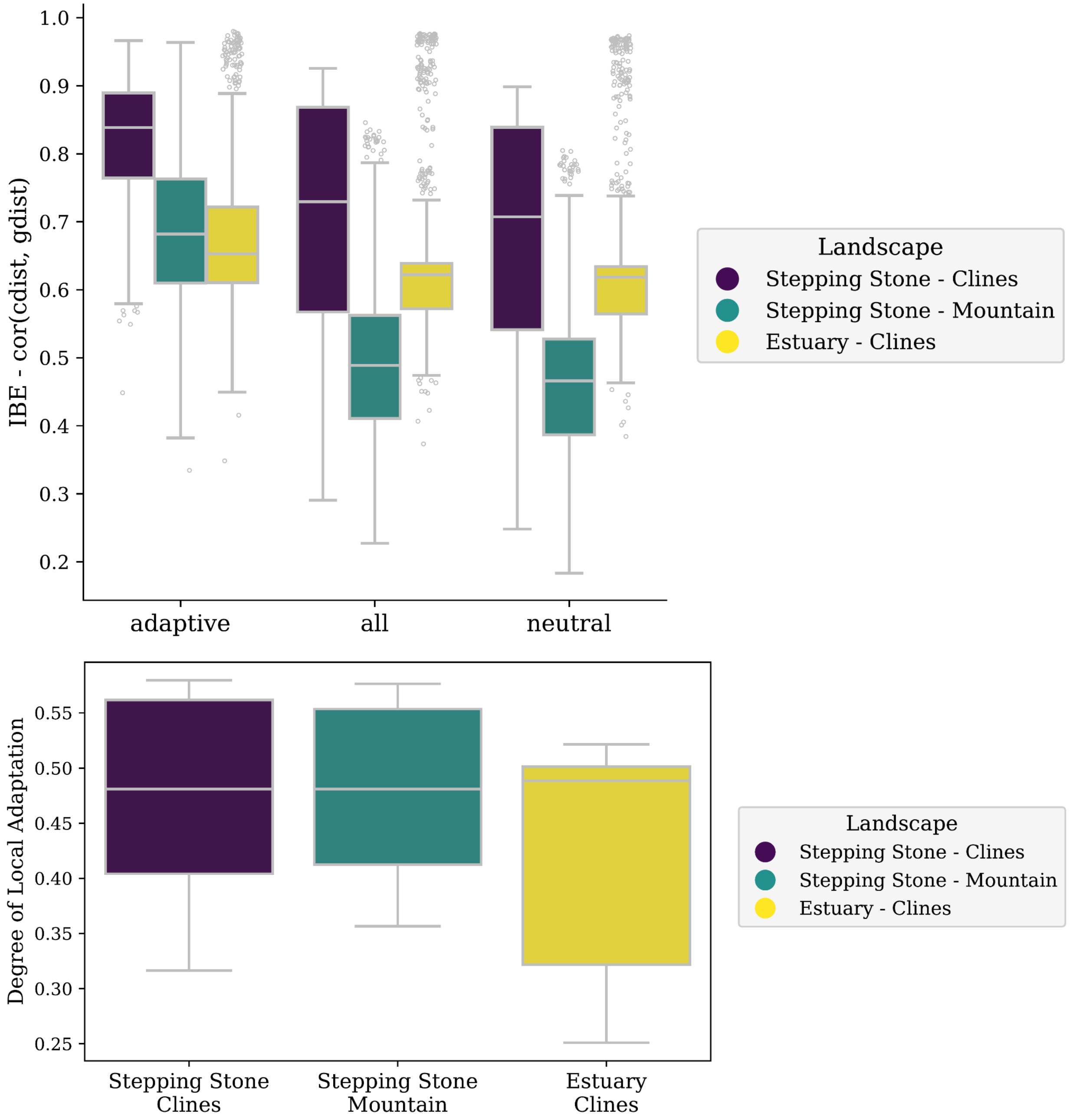

**Fig S17** Levels of isolation-by-environment in marker sets vary across landscapes (A) and the degree of local adaptation reached by metapopulations on these landscapes (B). The pattern in (A) given (B) is in contrast to patterns between levels of IBE and the degree of local adaptation (Fig. S29). *I IBE* is quantified as Spearman's rank correlation between population pairwise $F_{ST}$ (gdist) and Euclidean distance of adaptive environments (cdist). Data in this figure is from all 1- and 2-trait simulations. Code to create this figure can be found in SC 02.02.10.

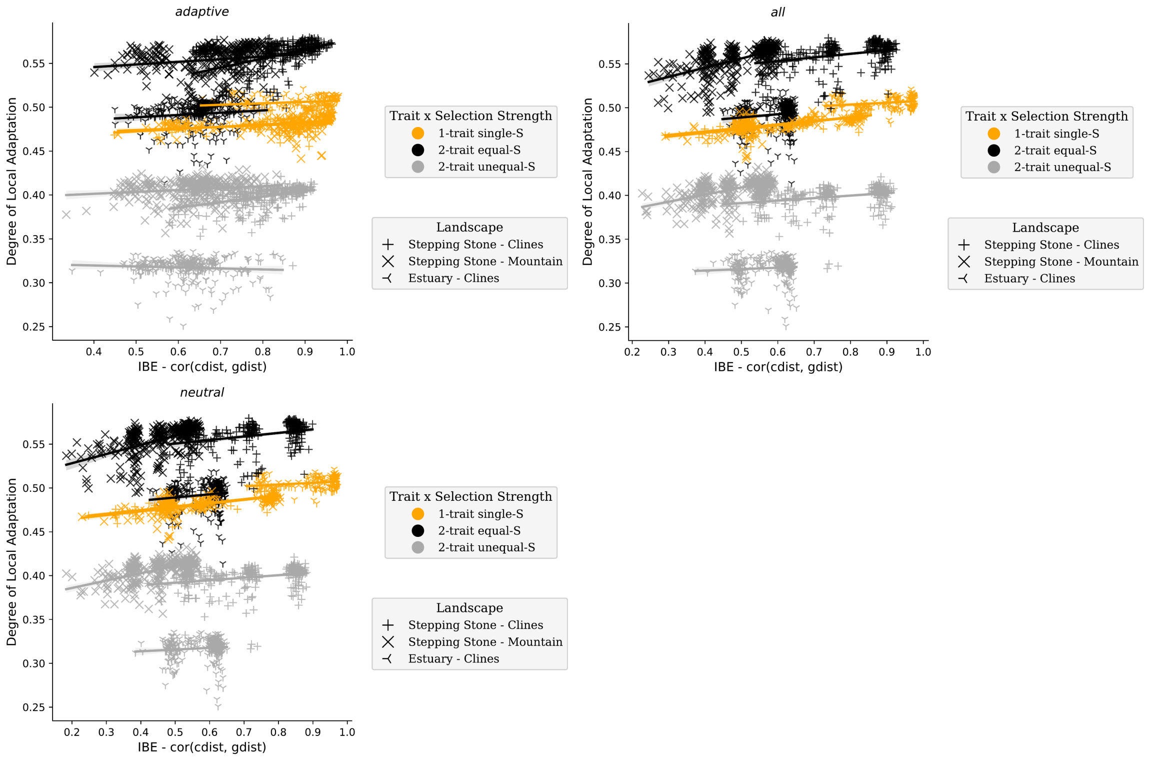

**Fig S18** The levels of isolation-by-distance in marker sets (panels) are weakly correlated with the degree of local adaptation ($LA_{\Delta SA}$) within simulation levels. *IBE* is quantified as Spearman's rank correlation between population pairwise $F_{ST}$ (gdist) and Euclidean distance of adaptive environments (cdist). Data included in this figure is from all marker sets from 1- and 2-trait simulations. Code to create this figure can be found in 02.02.10.

(Fig S19)

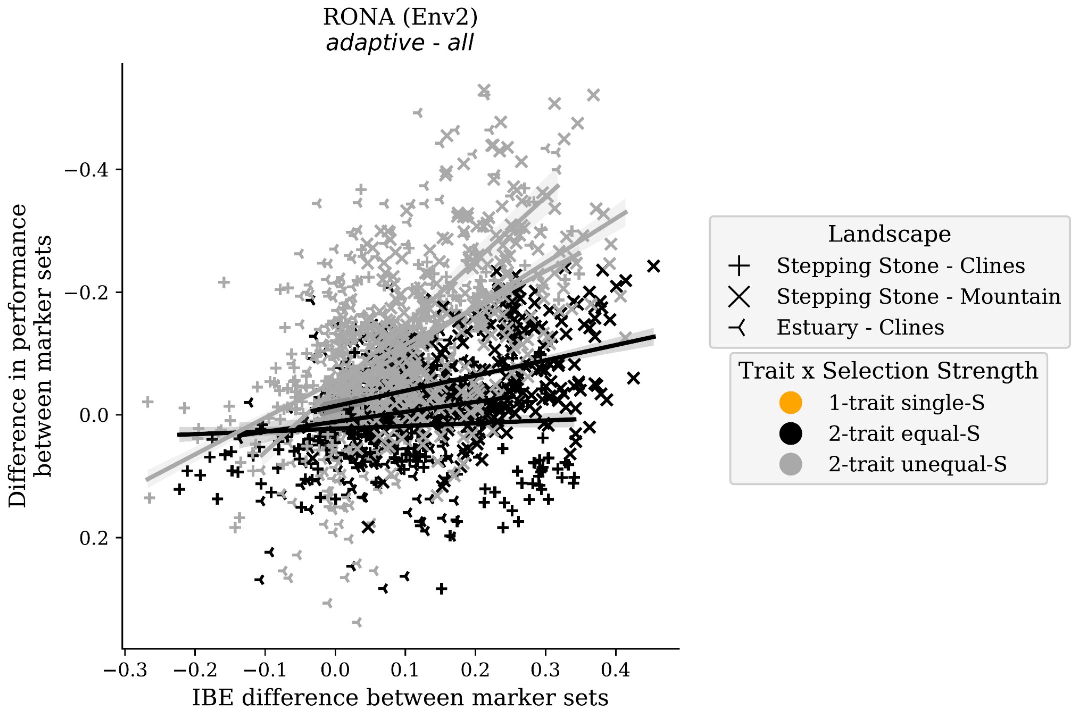
RONA (Env2)
adaptive - all
Difference in performance between marker sets
IBE difference between marker sets
−0.4
−0.2
0.0
0.2
−0.3
−0.2
−0.1
0.0
0.1
0.2
0.3
0.4
Landscape
Stepping Stone - Clines
Stepping Stone - Mountain
Estuary - Clines
Trait x Selection Strength
1-trait single-S
2-trait equal-S
2-trait unequal-S

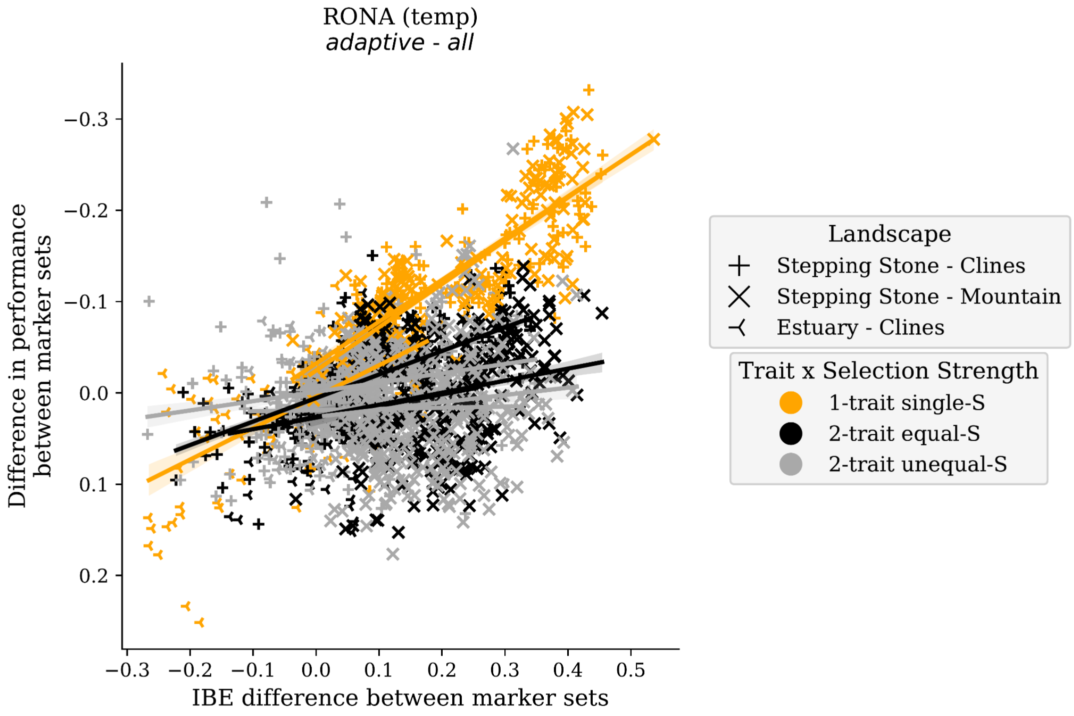
RONA (temp)
adaptive - all
Difference in performance between marker sets
IBE difference between marker sets
−0.3
−0.2
−0.1
0.0
0.1
0.2
−0.3
−0.2
−0.1
0.0
0.1
0.2
0.3
0.4
0.5
Landscape
Stepping Stone - Clines
Stepping Stone - Mountain
Estuary - Clines
Trait x Selection Strength
1-trait single-S
2-trait equal-S
2-trait unequal-S

(Fig S19 continued)

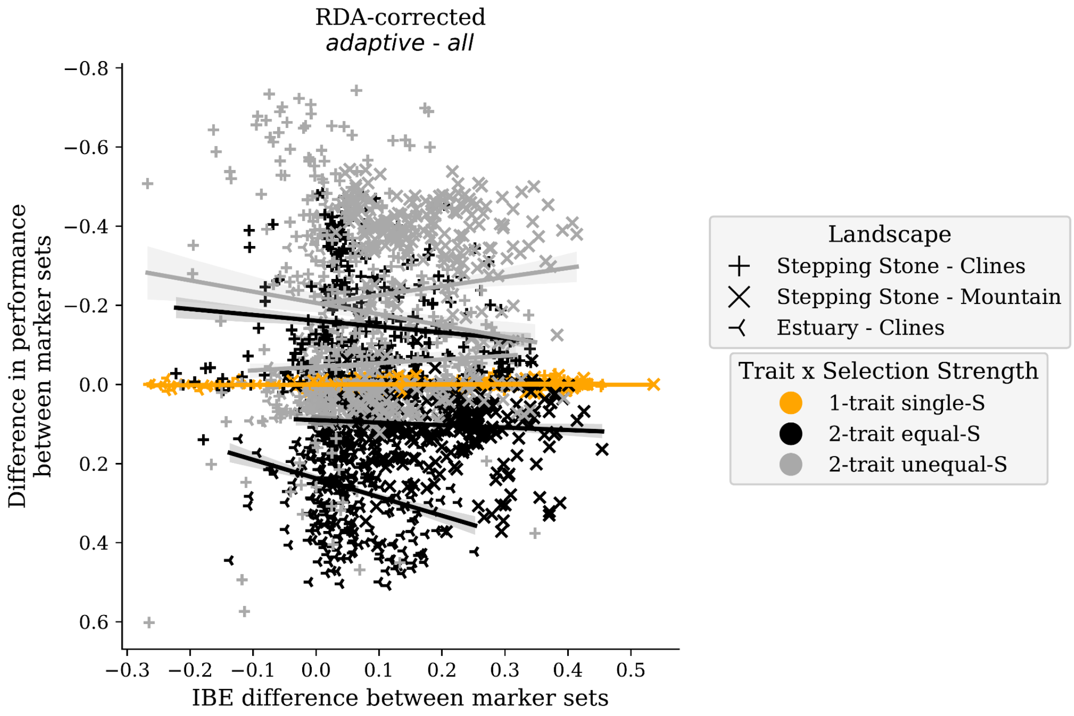

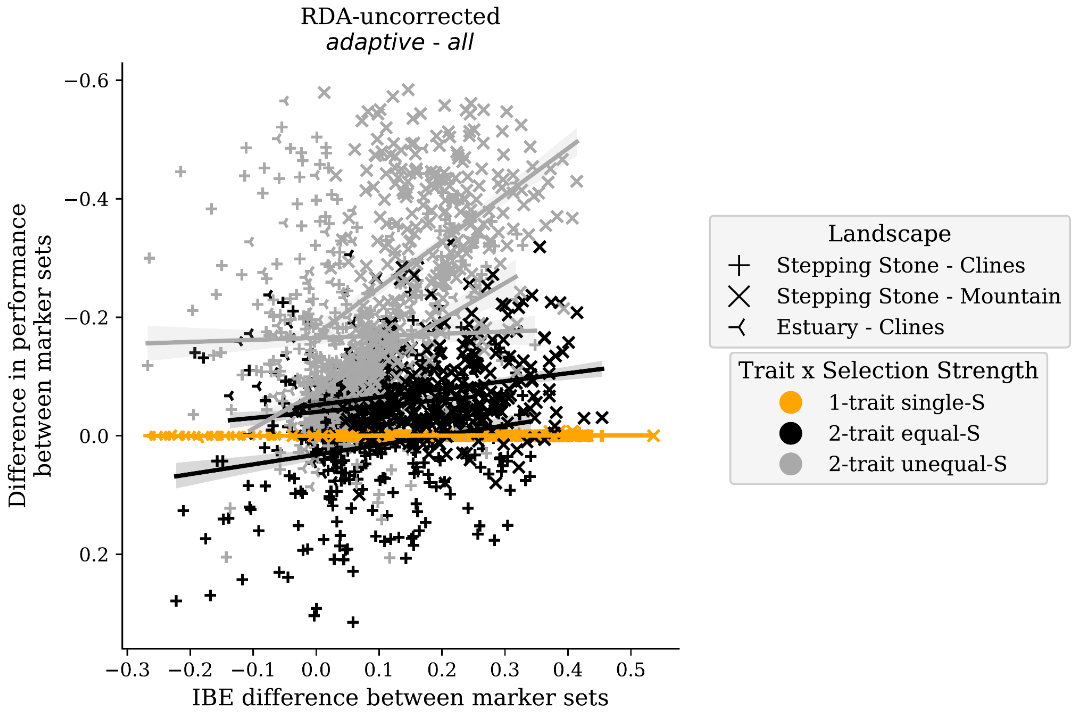

(Fig S19 continued)

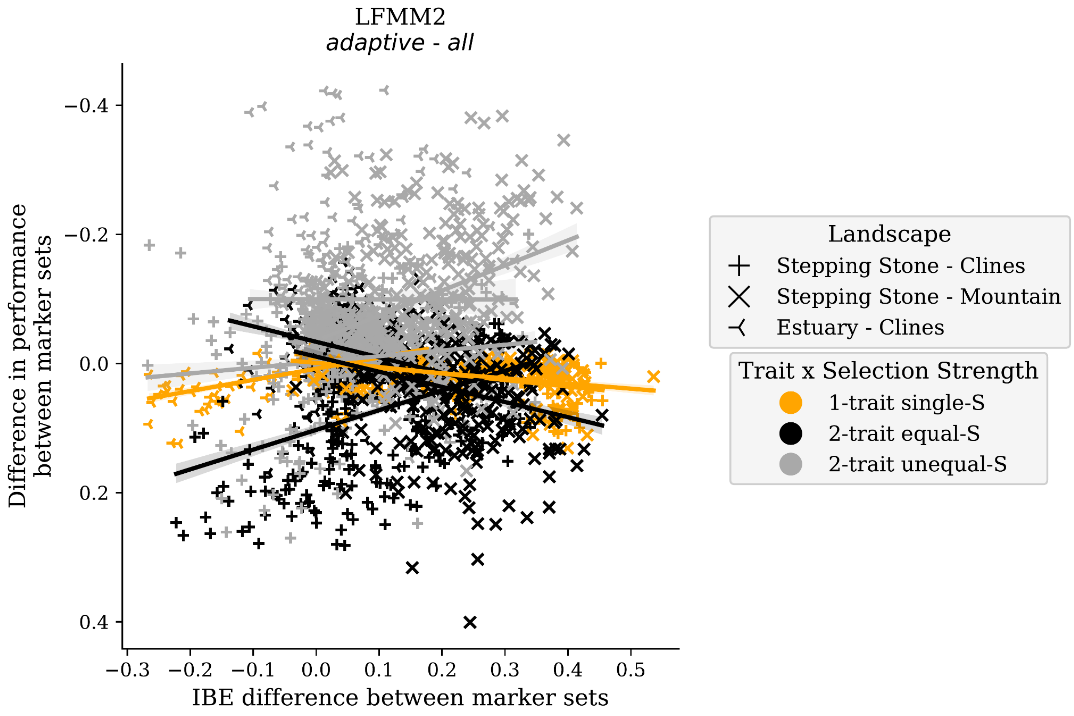

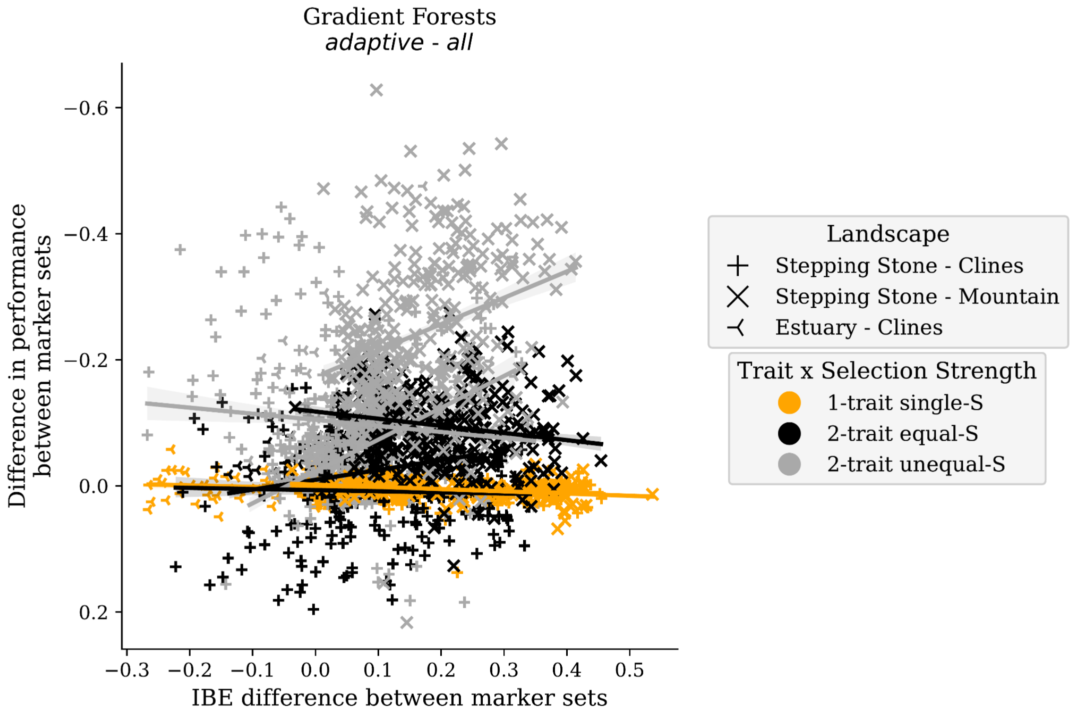

**Fig S19** Differences in levels of IBE between marker sets used to train models is generally unrelated to differences in model performances. Shown is the difference in median performance between *adaptive* and *all* marker sets and the difference in *IBE* between these marker sets. *IBE* is quantified as Spearman's rank correlation between population pairwise $F_{ST}$ and Euclidean distance of adaptive environments. Data in this figure is from 1- and 2-trait simulations. Code to create these figures can be found in SC 02.02.12.

| Latitude | 1 | 2 | 3 | 4 | 5 | 6 | 7 | 8 | 9 | 10 |
|---|---|---|---|---|---|---|---|---|---|---|
| **10** | 91.0 | 92.0 | 93.0 | 94.0 | 95.0 | 96.0 | 97.0 | 98.0 | 99.0 | 100.0 |
| **9** | 81.0 | 82.0 | 83.0 | 84.0 | 85.0 | 86.0 | 87.0 | 88.0 | 89.0 | 90.0 |
| **8** | 71.0 | 72.0 | 73.0 | 74.0 | 75.0 | 76.0 | 77.0 | 78.0 | 79.0 | 80.0 |
| **7** | 61.0 | 62.0 | 63.0 | 64.0 | 65.0 | 66.0 | 67.0 | 68.0 | 69.0 | 70.0 |
| **6** | 51.0 | 52.0 | 53.0 | 54.0 | 55.0 | 56.0 | 57.0 | 58.0 | 59.0 | 60.0 |
| **5** | 41.0 | 42.0 | 43.0 | 44.0 | 45.0 | 46.0 | 47.0 | 48.0 | 49.0 | 50.0 |
| **4** | 31.0 | 32.0 | 33.0 | 34.0 | 35.0 | 36.0 | 37.0 | 38.0 | 39.0 | 40.0 |
| **3** | 21.0 | 22.0 | 23.0 | 24.0 | 25.0 | 26.0 | 27.0 | 28.0 | 29.0 | 30.0 |
| **2** | 11.0 | 12.0 | 13.0 | 14.0 | 15.0 | 16.0 | 17.0 | 18.0 | 19.0 | 20.0 |
| **1** | 1.0 | 2.0 | 3.0 | 4.0 | 5.0 | 6.0 | 7.0 | 8.0 | 9.0 | 10.0 |

Longitude

**Fig S20** A map of Garden ID (unbolded entries) across each landscape for 1-, 2- and 6-trait simulations (latitudinal and longitudinal grids are bolded). This map can be used to interpret the ordering of gardens along x-axes of Figs. S21 S22 and S23. Code used to create this figure can be found in SC 02.02.04.

(Fig. S21)

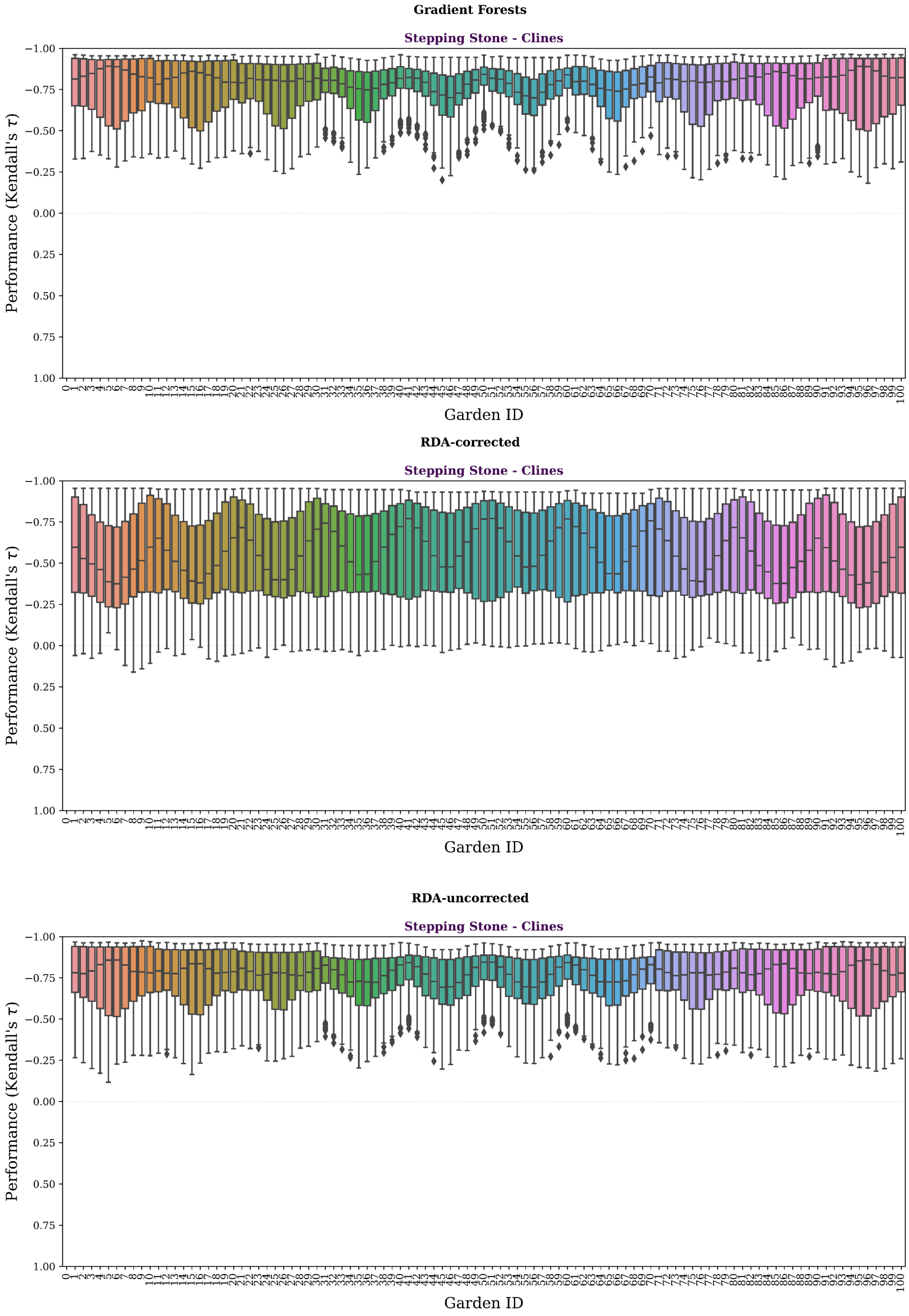

(Fig S21 continued)

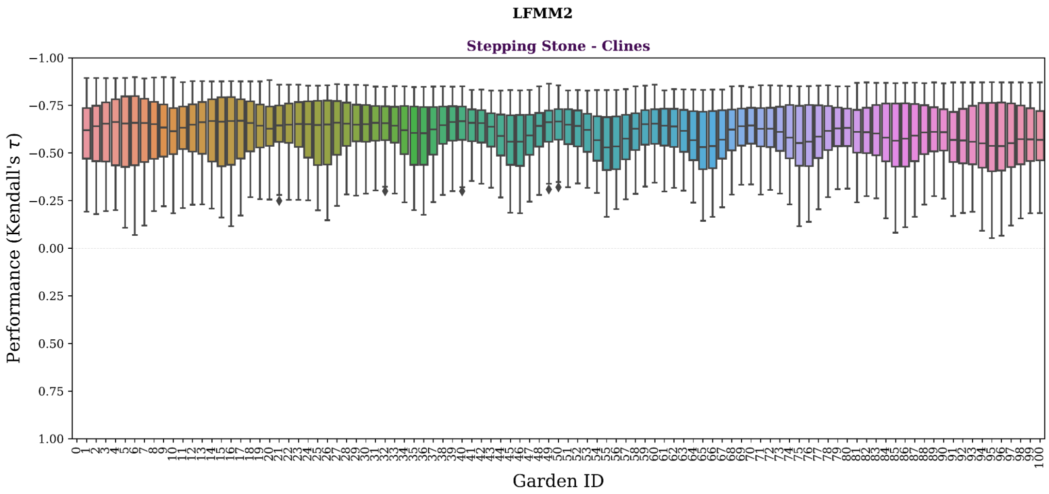
LFMM2
Stepping Stone - Clines
Performance (Kendall's τ)
Garden ID

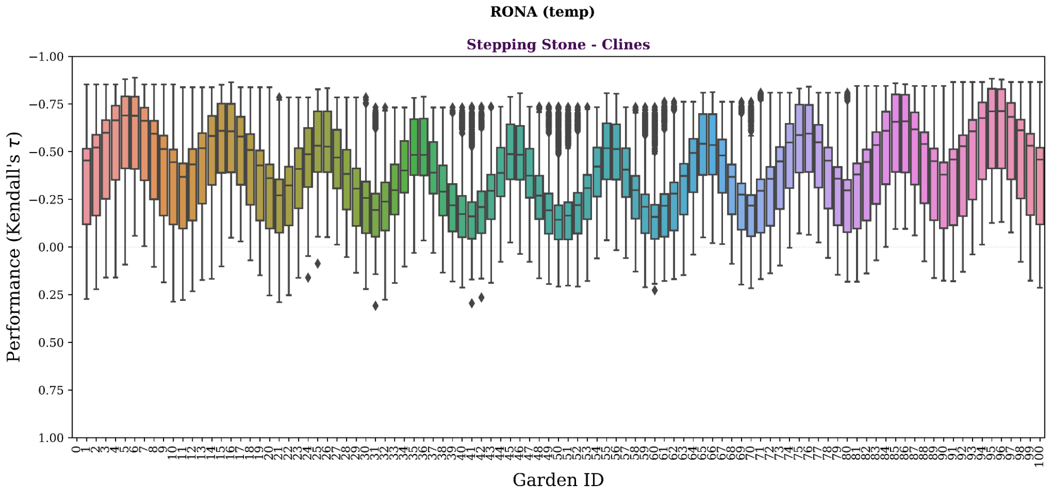
RONA (temp)
Stepping Stone - Clines
Performance (Kendall's τ)
Garden ID

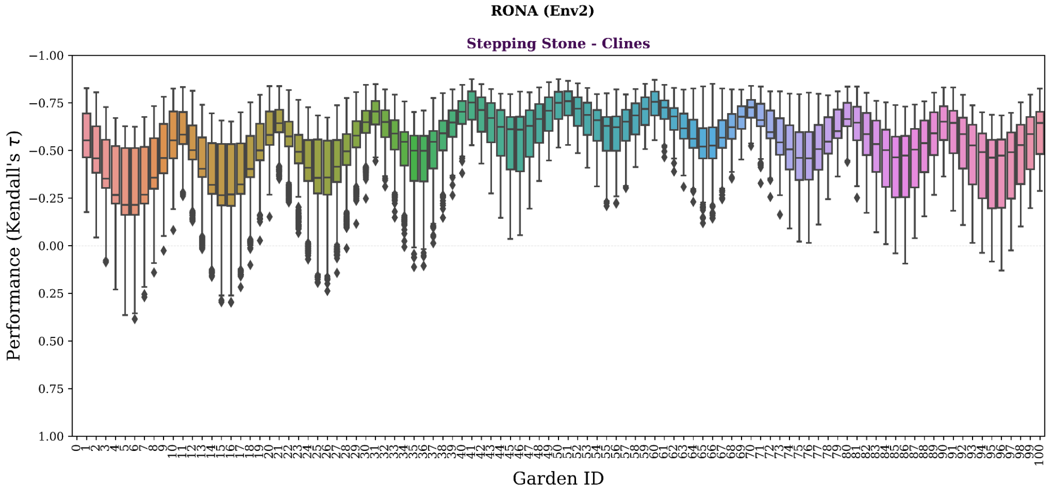
RONA (Env2)
Stepping Stone - Clines
Performance (Kendall's τ)
Garden ID

**Fig S21** Genomic offset methods have variable performance across the *Stepping-Stone Clines* landscape. Shown is the variability of each offset method performance (y-axes) across the 100 common gardens (x-axes). Gardens are ordered from left to right by garden ID. This ordering of gardens is equivalent to the southwestern-most garden first and northeastern-most garden last (see Fig. S20 for a map of garden ID across each landscape). Similar figures for *Stepping-Stone Mountain* and *Estuary-Clines* landscapes can be found in Fig S22 and Fig S23, respectively. Data included in this figure is from evaluation of 1- and 2-trait simulations using *all* markers. Code used to create these figures can be found in SC 02.02.04.

(Fig S22)

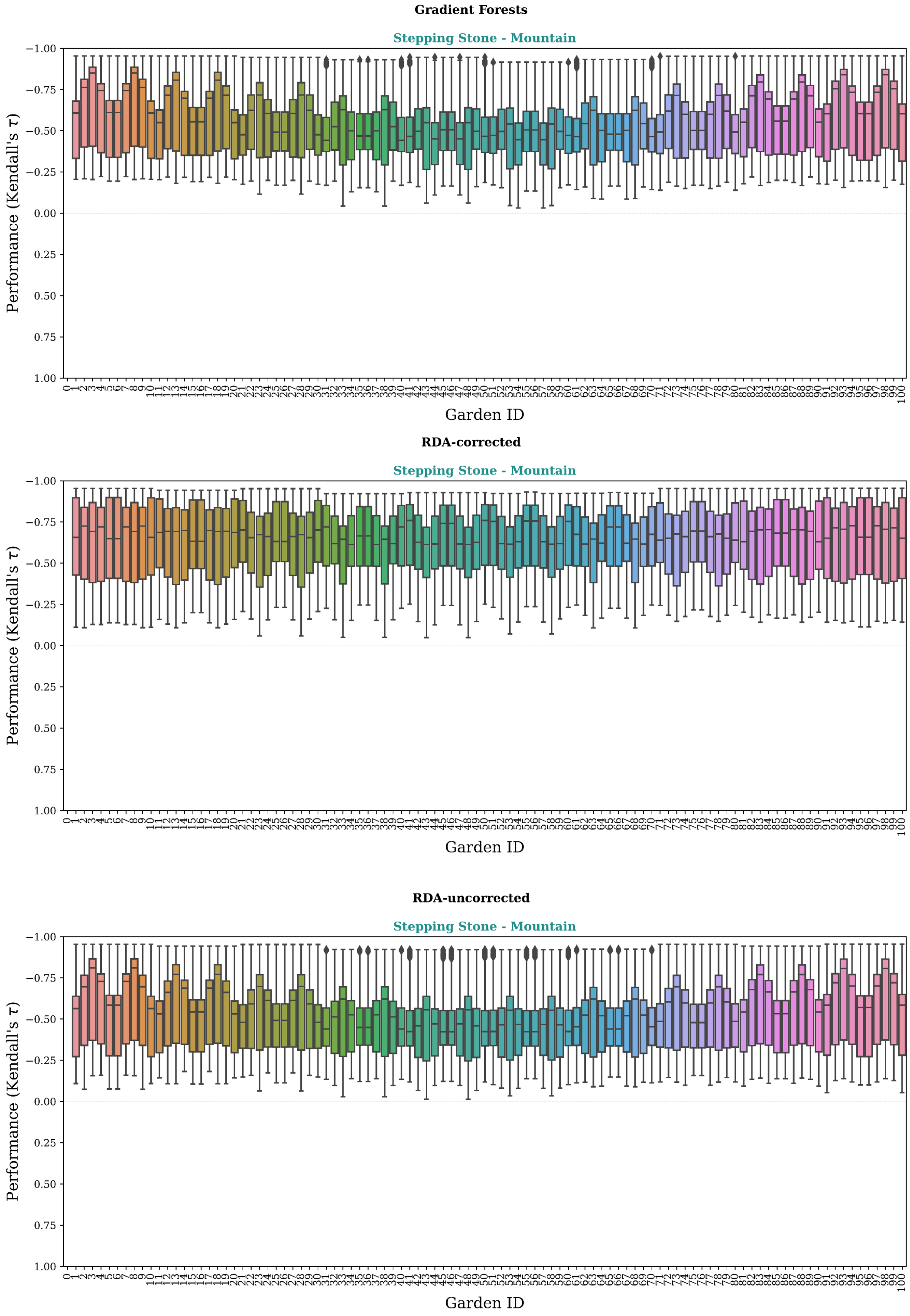
Gradient Forests
Stepping Stone - Mountain
Performance (Kendall's τ)
Garden ID
RDA-corrected
Stepping Stone - Mountain
Performance (Kendall's τ)
Garden ID
RDA-uncorrected
Stepping Stone - Mountain
Performance (Kendall's τ)
Garden ID

(Fig S22 continued)

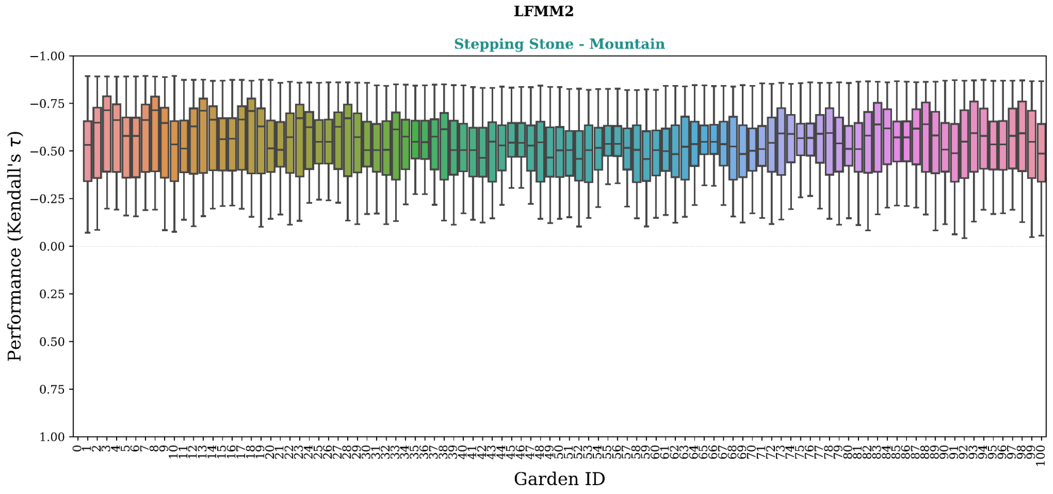

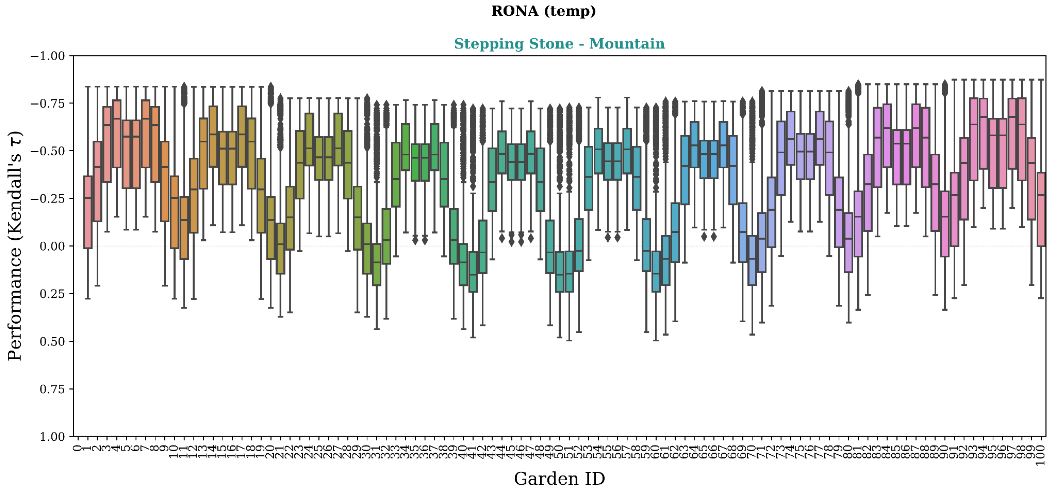

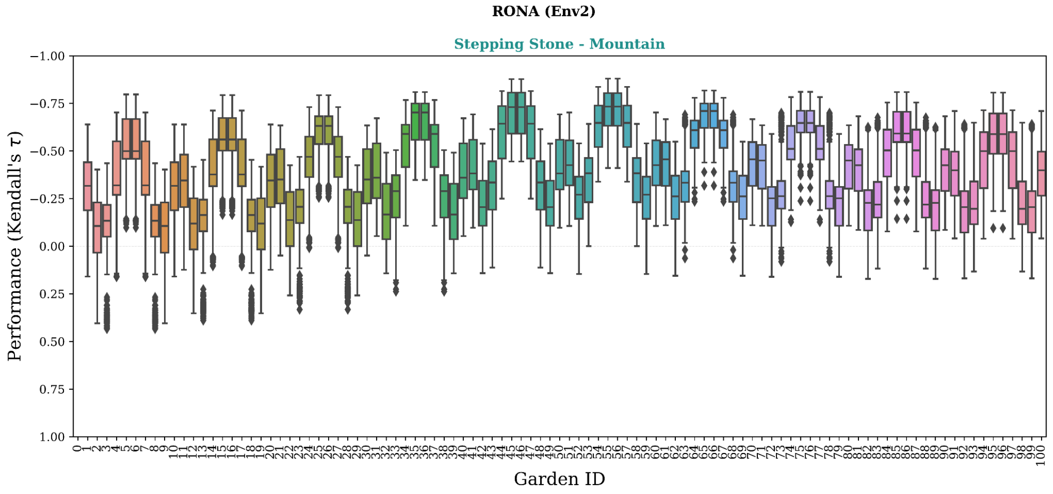

**Fig S22** Genomic offset methods have variable performance across the *Stepping-Stone - Mountain* landscape. Shown is the variability of each offset method performance (y-axes) across the 100 common gardens (x-axes). Gardens are ordered from left to right by garden ID. This ordering of gardens is equivalent to the southwestern-most garden first and northeastern-most garden last (see Fig. S20 for a map of garden ID across each landscape). Similar figures for *Stepping-Stone - Clines* and *Estuary - Clines* landscapes can be found in Fig S21 and Fig S23, respectively. Data included in this figure is from evaluation of 1- and 2-trait simulations using *all* markers. Code used to create this figure can be found in SC 02.02.04.

(Fig S23)

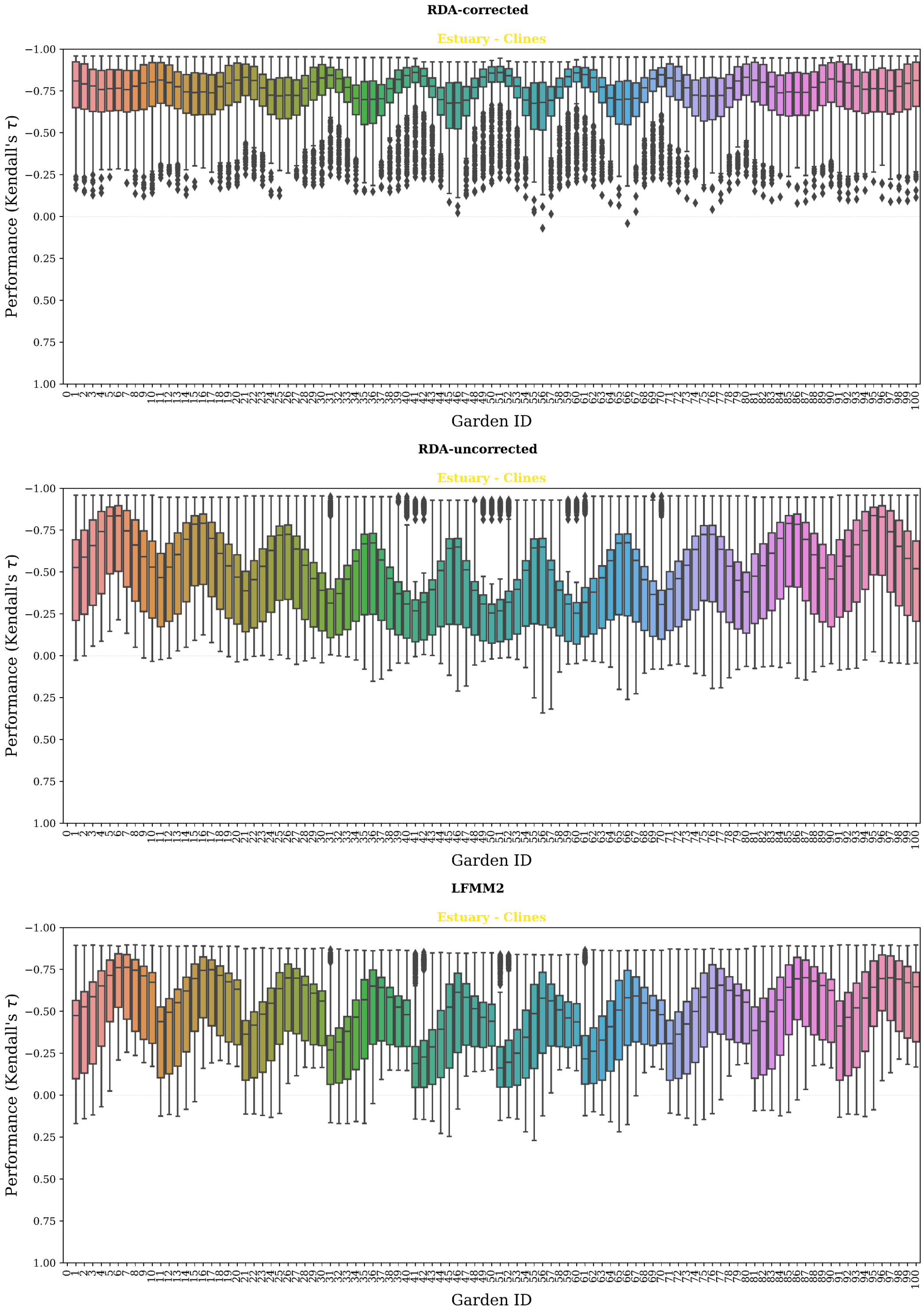
RDA-corrected
Estuary - Clines
Performance (Kendall's τ)
Garden ID
RDA-uncorrected
Estuary - Clines
Performance (Kendall's τ)
Garden ID
LFMM2
Estuary - Clines
Performance (Kendall's τ)
Garden ID

(Fig S23 continued)

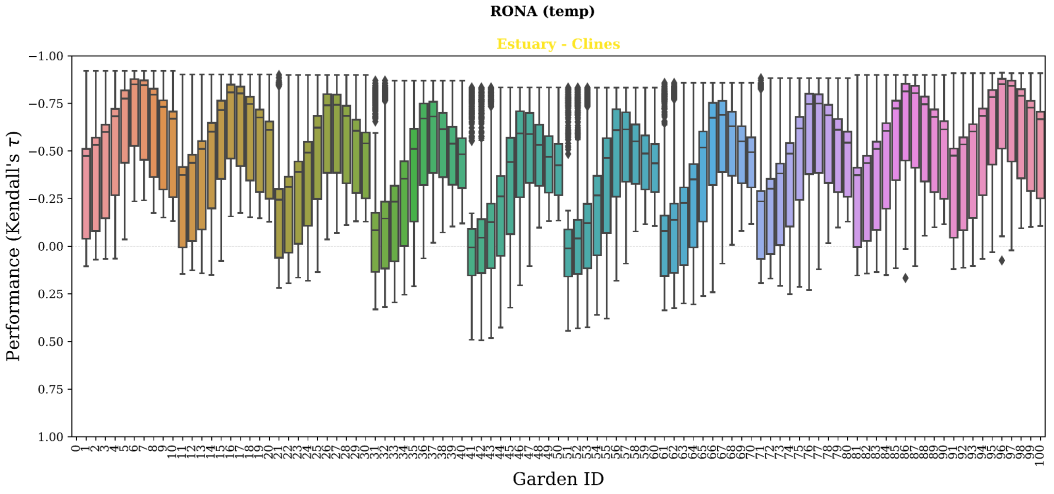

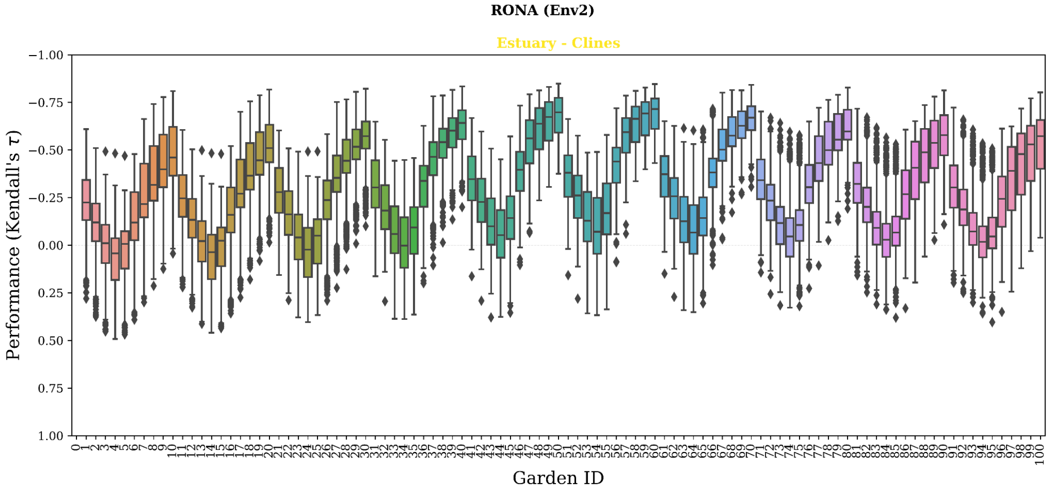

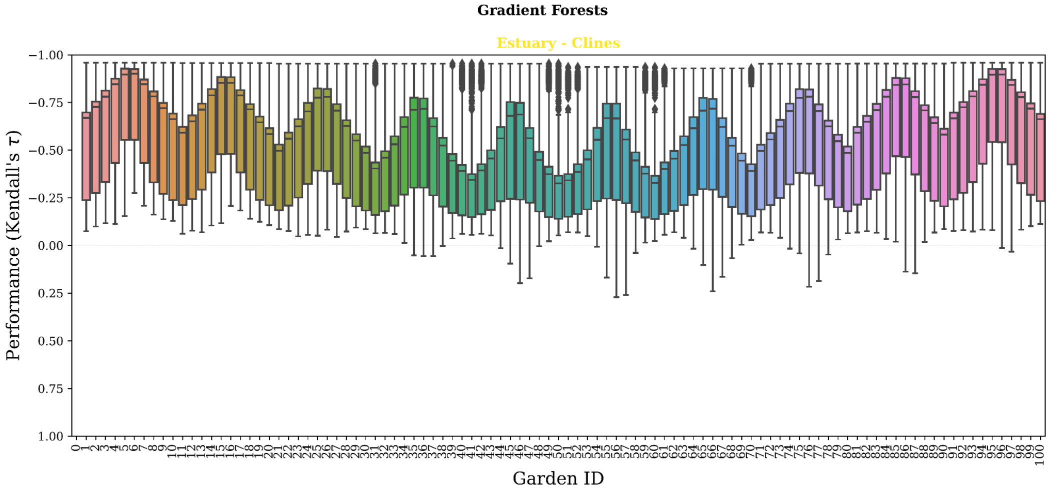

**Fig S23** Genomic offset methods have variable performance across the *Estuary - Clines* landscape. Shown is the variability of each offset method performance (y-axes) across the 100 common gardens (x-axes). Gardens are ordered from left to right by garden ID. This ordering of gardens is equivalent to the southwestern-most garden first and northeastern-most garden last (see Fig. S20 for a map of garden ID across each landscape). Similar figures for *Stepping-Stone - Clines* and *Stepping-Stone - Mountain* landscapes can be found in Fig S21 and Fig S22, respectively. Data included in this figure is from evaluation of 1- and 2-trait simulations using *all* markers. Code used to create this figure can be found in SC 02.02.04.

(Fig. S36)

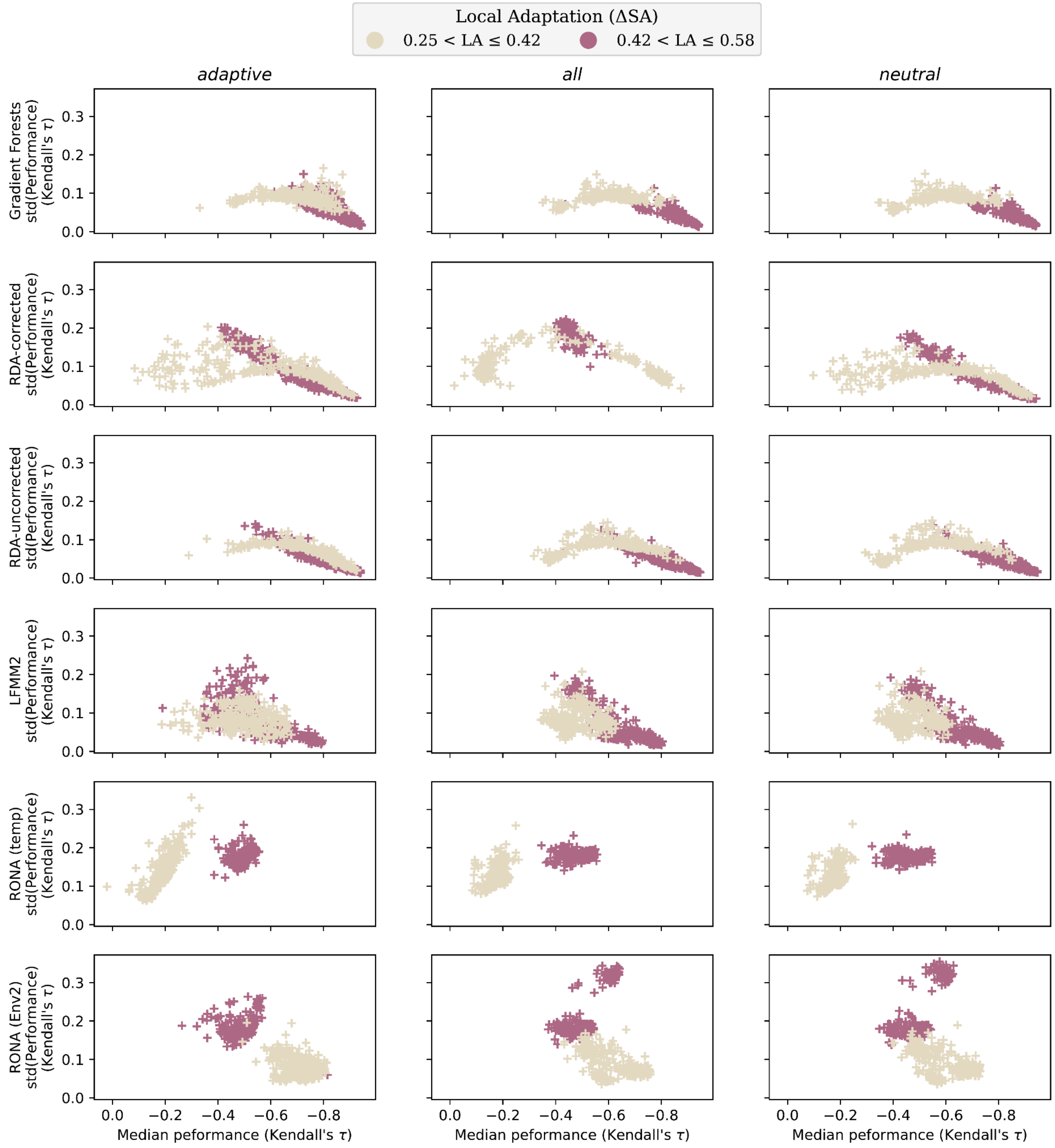
Local Adaptation (ΔSA)
0.25 < LA ≤ 0.42
0.42 < LA ≤ 0.58
adaptive
all
neutral
Gradient Forests std(Performance) (Kendall's τ)
RDA-corrected std(Performance) (Kendall's τ)
RDA-uncorrected std(Performance) (Kendall's τ)
LFMM2 std(Performance) (Kendall's τ)
RONA (temp) std(Performance) (Kendall's τ)
RONA (Env2) std(Performance) (Kendall's τ)
0.0
0.1
0.2
0.3
0.0
−0.2
−0.4
−0.6
−0.8
Median peformance (Kendall's τ)

**Fig S24** Variability of genomic offset performance (y-axes) for a given model (+) often decreases with increasing median performance (x-axes). Shown are patterns from each offset method (rows) for each marker set (columns) used in training. Data included in this figure is from evaluation of 2-trait simulations from *Stepping-Stone - Clines* landscapes processed through the *Adaptive Environment* workflow. For similar figures for *Stepping-Stone - Mountain* and *Estuary - Clines* landscapes, see Figs. S25-S26, respectively. Code used to create these figures can be found in SC 02.02.07.

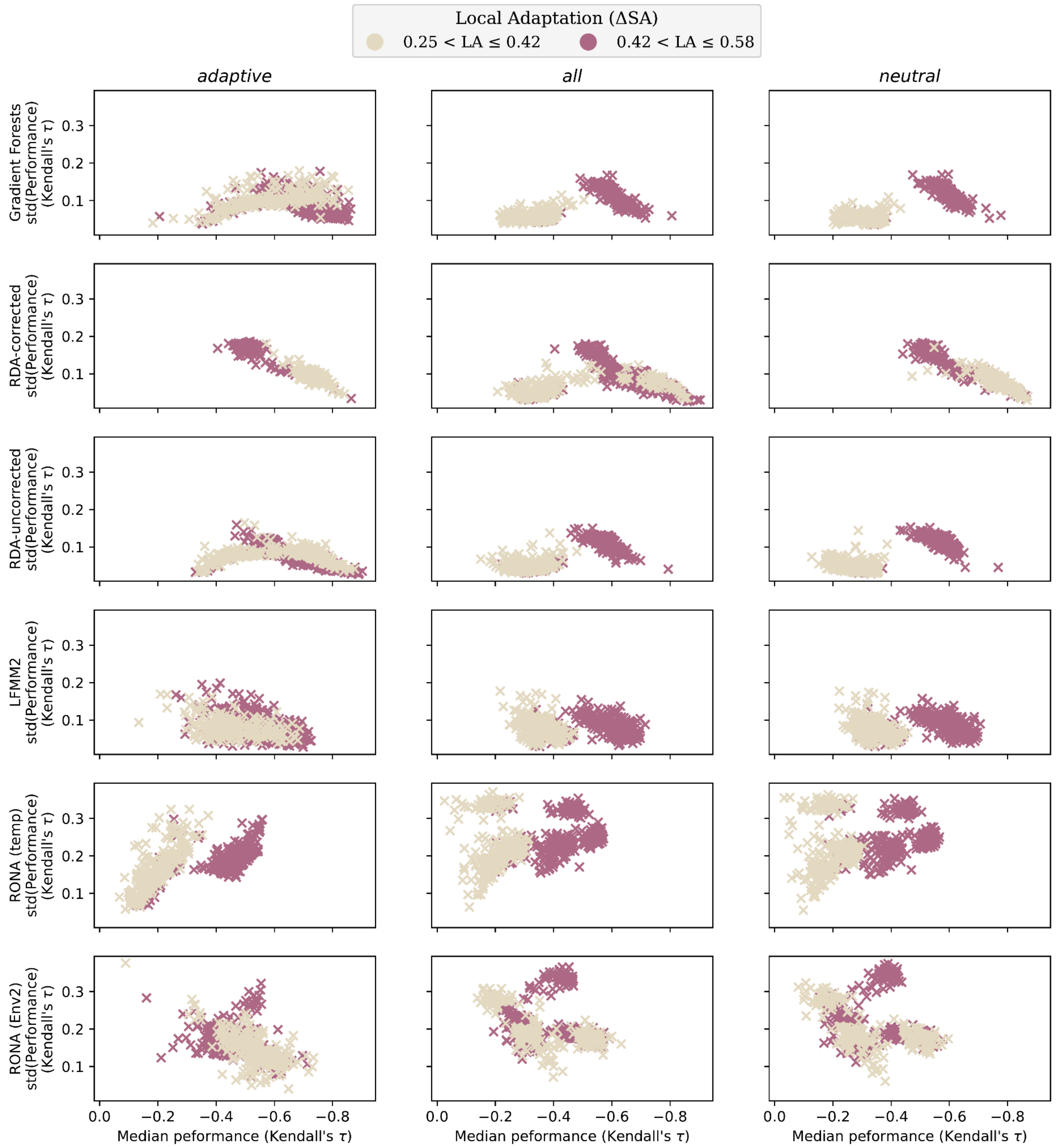

**Fig S25** Variability across evaluations of genomic offsets often decreases with increasing average performance across marker sets. Data included in this figure is from evaluation of 2-trait simulations from *Stepping-Stone - Mountain* landscapes. For similar figures for *Stepping-Stone - Clines* and *Estuary - Clines* landscapes, see Figs. 24 and S26, respectively. Code used to create these figures can be found in SC 02.02.07.

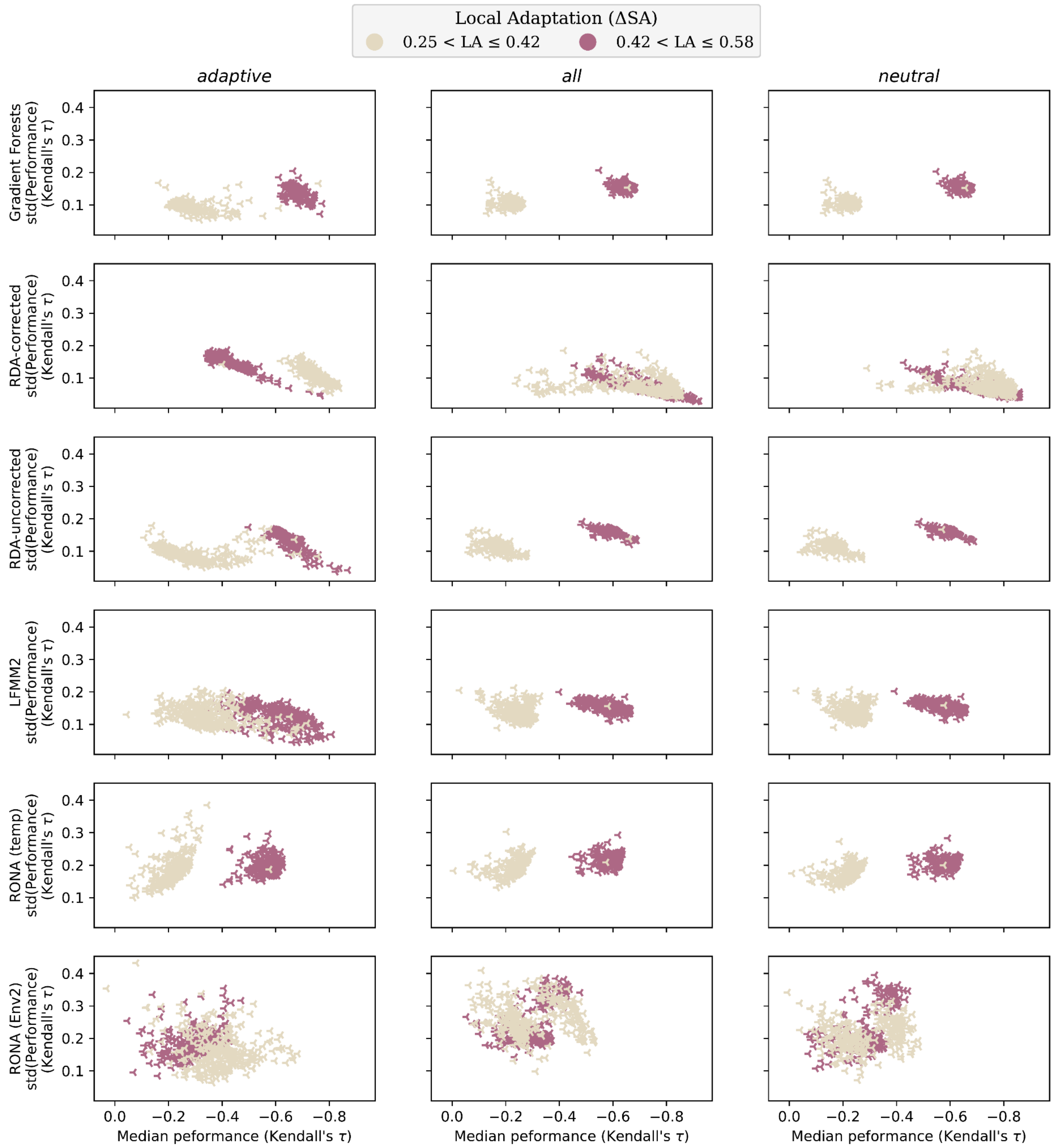

**Fig S26** Variability across evaluations of genomic offsets often decreases with increasing average performance across marker sets. Data included in this figure is from evaluation of 2-trait simulations from *Estuary - Clines* landscapes. For similar figures for *Stepping-Stone - Clines* and *Stepping-Stone - Mountain* landscapes, see Figs. S24 and S25, respectively. Code used to create these figures can be found in SC 02.02.07.

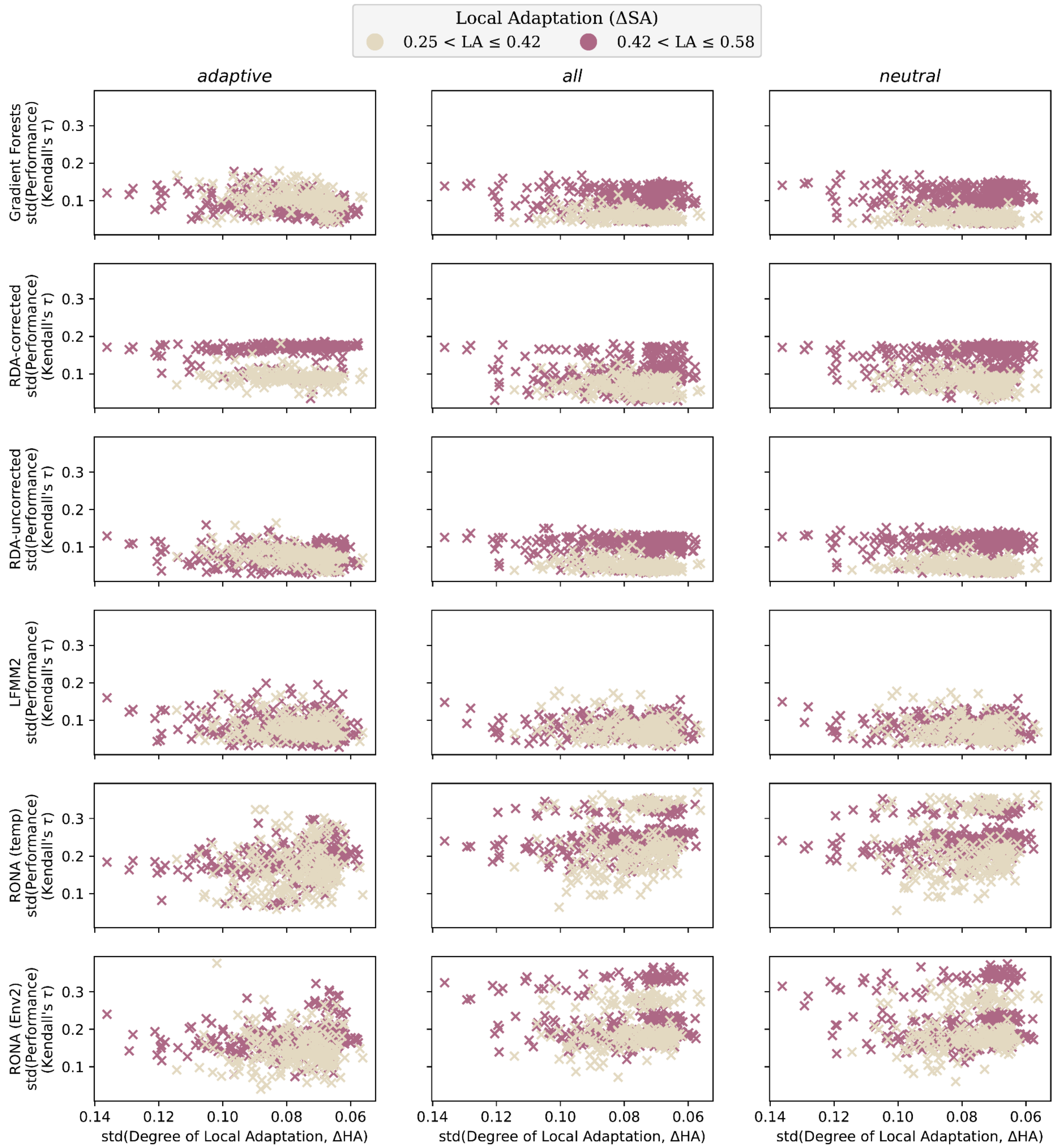

**Fig S27** Variability across evaluations of genomic offsets is often unrelated to the variability in the degree of local adaptation across populations. Data included in this figure is from evaluation of 2-trait simulations from *Stepping-Stone - Mountain* landscapes. For similar figures for *Stepping-Stone - Clines* and *Estuary - Clines* landscapes, see Figs. S27 and S28, respectively. Code used to create these figures can be found in SC 02.02.07.

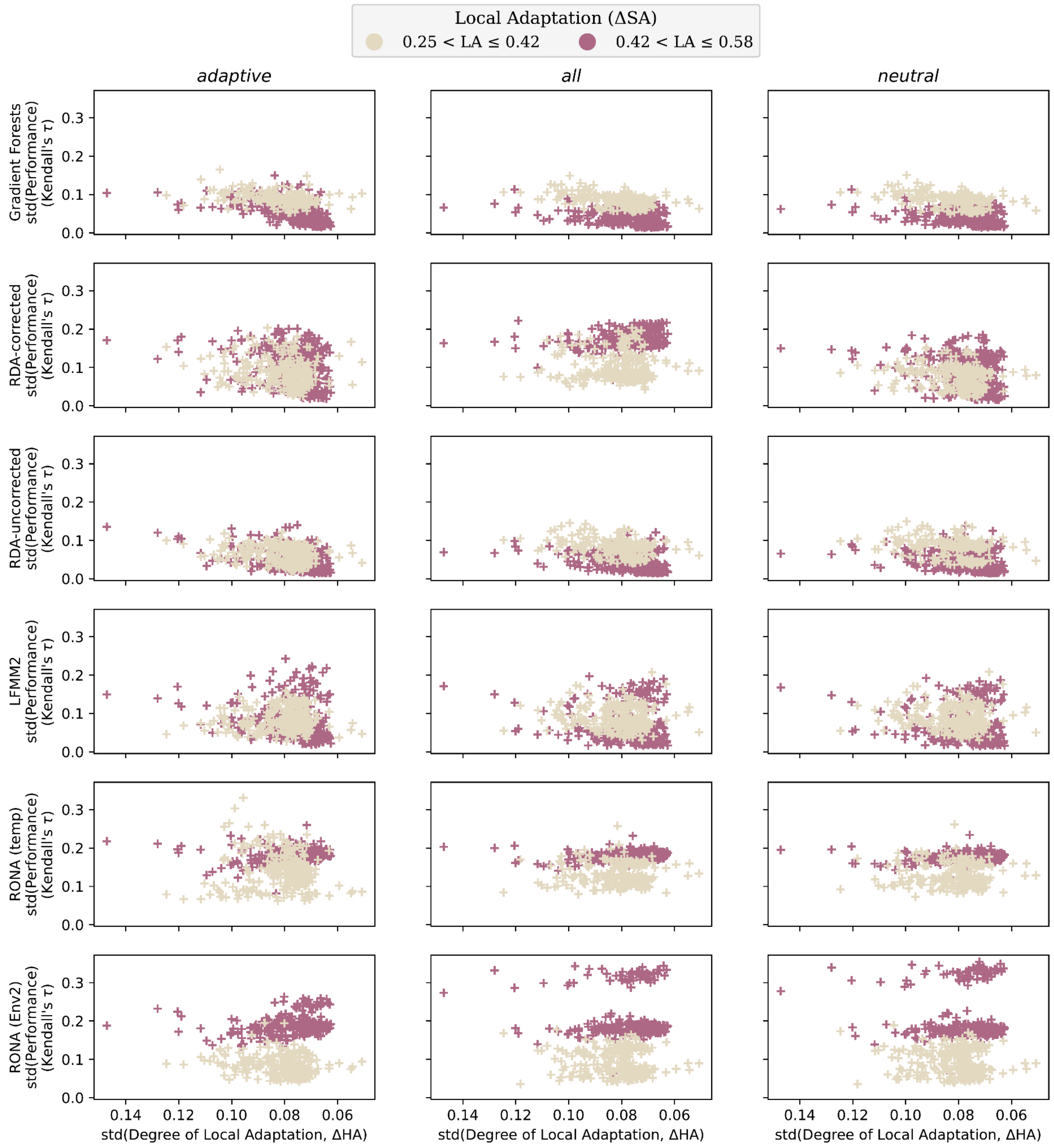

**Fig S28** Variability across evaluations of genomic offsets is often unrelated to the variability in the degree of local adaptation across populations. Data included in this figure is from evaluation of 2-trait simulations from *Stepping Stone - Clines* landscapes. For similar figures for *Estuary - Clines* and *Stepping-Stone - Mountain* landscapes, see Figs. S26 and S28, respectively. Code used to create these figures can be found in SC 02.02.07.

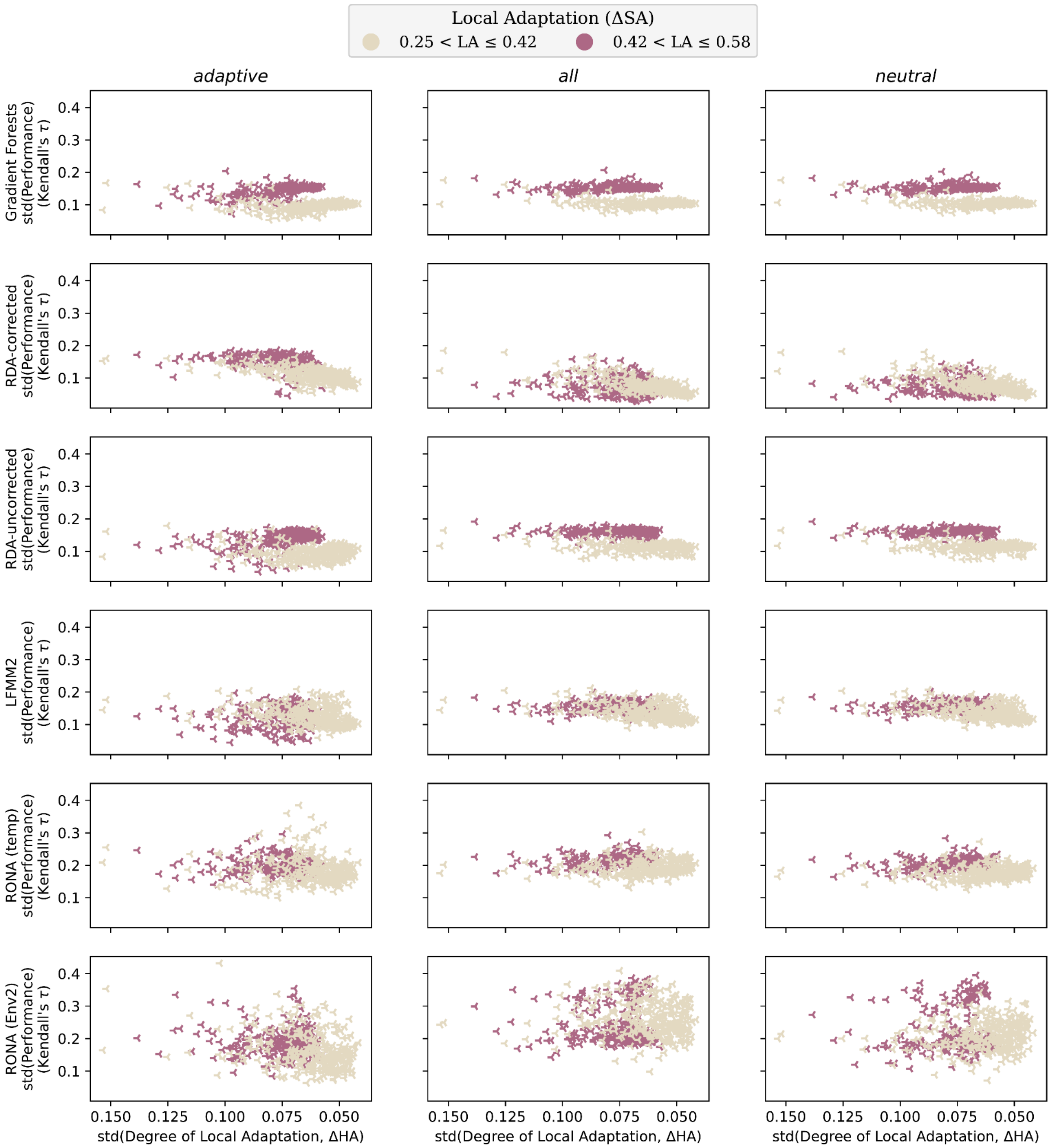

**Fig S29** Variability across evaluations of genomic offsets is often unrelated to the variability in the degree of local adaptation across populations. Data included in this figure is from evaluation of 2-trait simulations from *Estuary - Clines* landscapes. For similar figures for *Stepping-Stone - Clines* and *Stepping-Stone - Mountain* landscapes, see Figs. S26 and S27, respectively. Code used to create these figures can be found in SC 02.02.07.

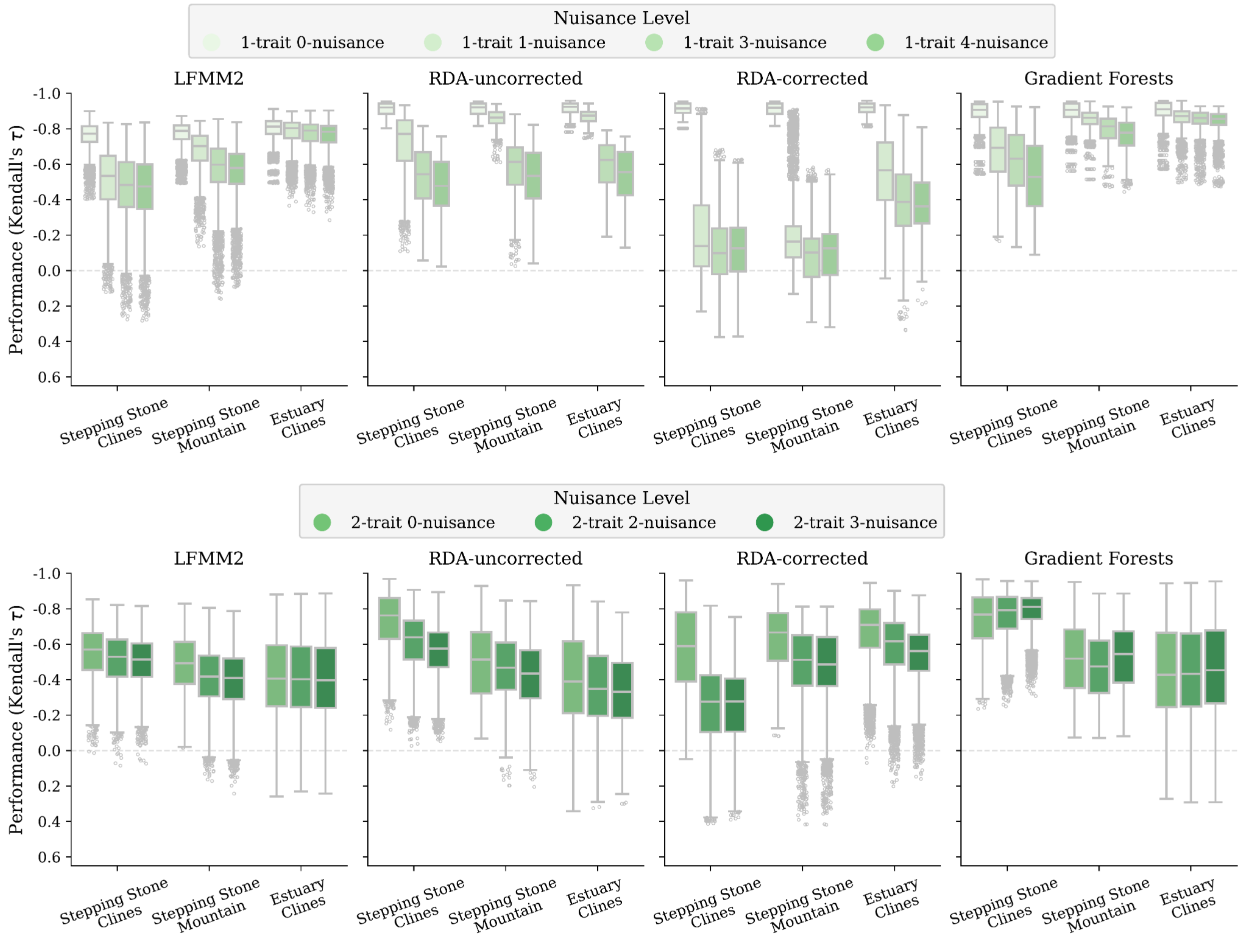

**Fig S30** Effect of non-adaptive nuisance environmental variables on offset performance faceted by landscape. Shown are offsets from 1- and 2-trait simulations trained using only adaptive environments (0-nuisance) or with adaptive environments and the addition of N>0 non-adaptive environmental variables (N-nuisance). RONA is not shown because it is univariate with respect to environmental variables. The nuisance variables for 1-trait simulations are: Env2, ISO, TSsd, PSsd; and for 2-trait simulations are ISO, TSsd, PSsd; see Table 2. The *Nuisance Environment* workflow was used to produce this data. Code to create these figures can be found in SC 02.02.06.

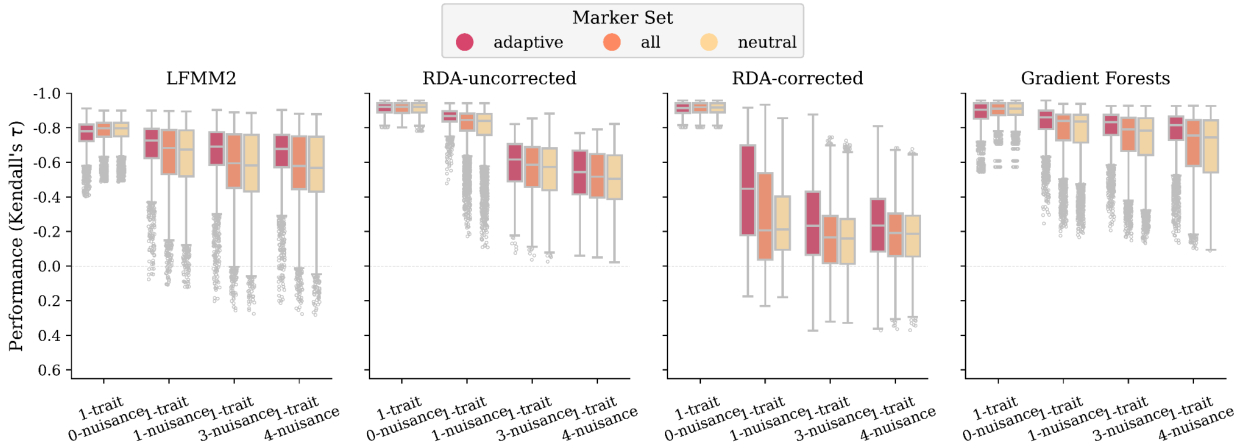

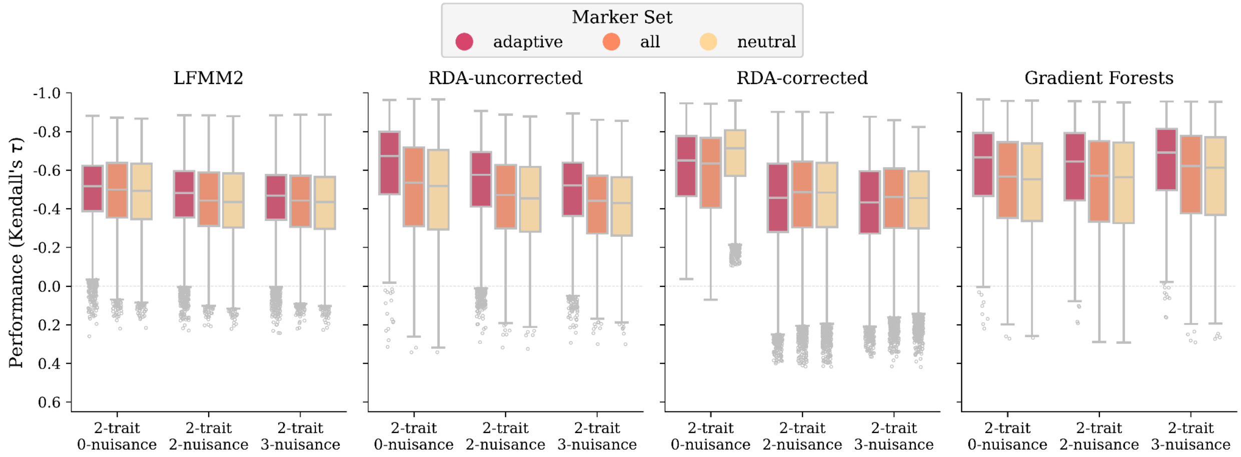

**Fig S31** Effect of non-adaptive nuisance environmental variables on offset performance faceted by marker set. Shown are offsets from 1- (A) and 2-trait (B) simulations trained using only adaptive environments (0-nuisance) or with adaptive environments and the addition of N>0 non-adaptive environmental variables (N-nuisance). RONA is not shown because it is univariate with respect to environmental variables. The nuisance variables for 1-trait simulations are: Env2, ISO, TSsd, PSsd; and for 2-trait simulations are ISO, TSsd, PSsd; see Table 2. Code to create these figures can be found in SC 02.02.06.

(Fig. S31)

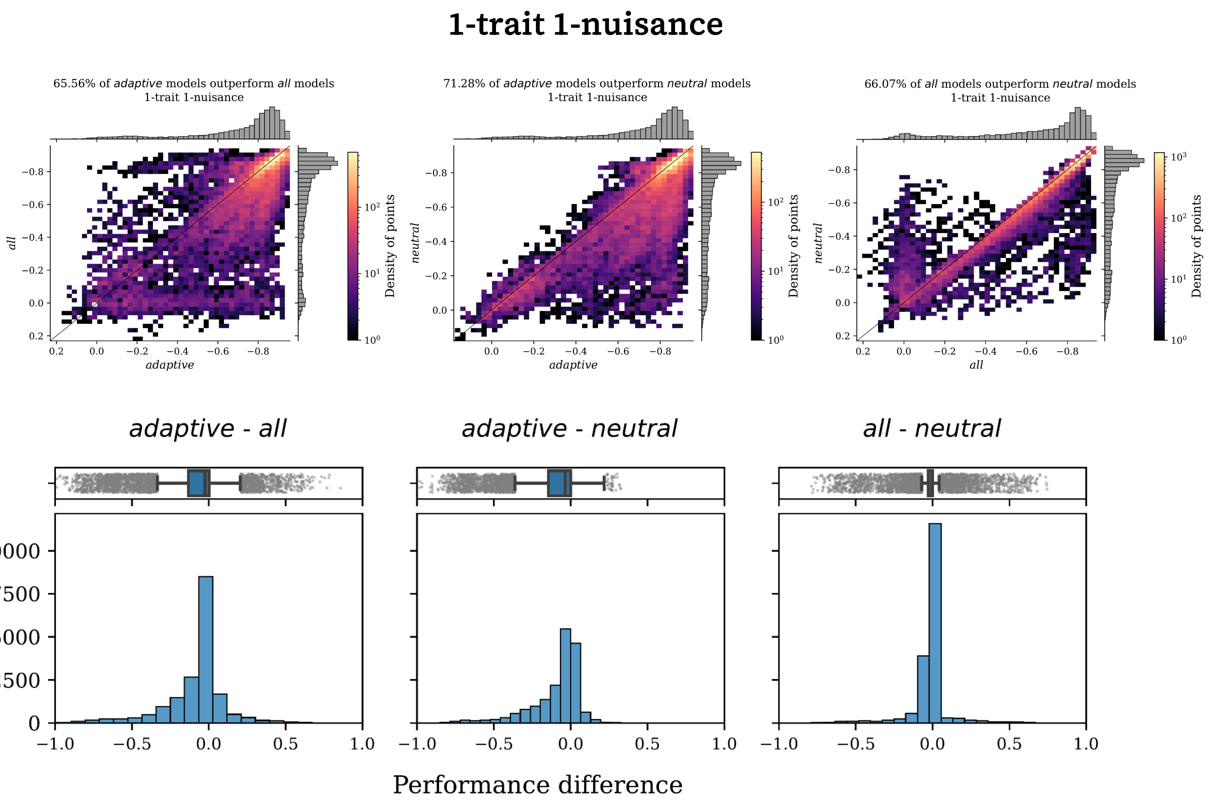
1-trait 1-nuisance
65.56% of adaptive models outperform all models
1-trait 1-nuisance
71.28% of adaptive models outperform neutral models
1-trait 1-nuisance
66.07% of all models outperform neutral models
1-trait 1-nuisance
adaptive
all
neutral
Density of points
adaptive - all
adaptive - neutral
all - neutral
Performance difference

(Fig. S31 continued)

**1-trait 3-nuisance**

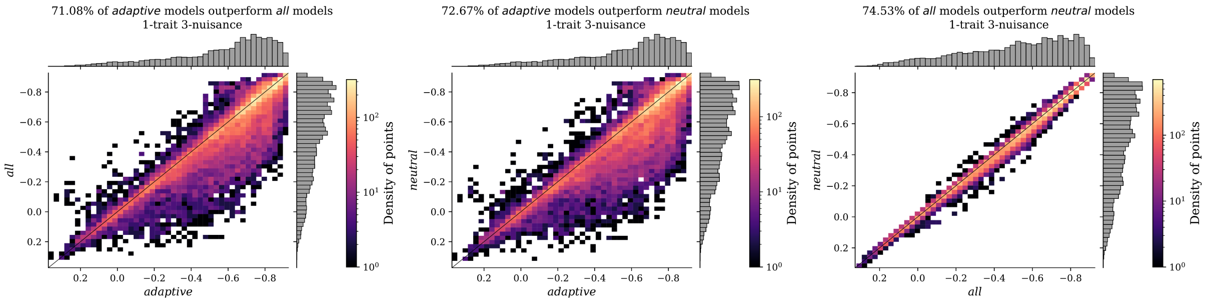

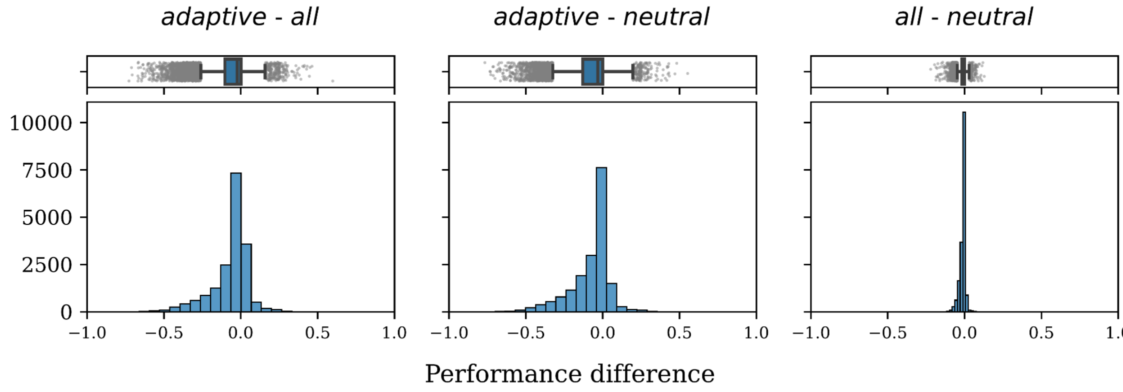

(Fig. S31 continued)

**1-trait 4-nuisance**

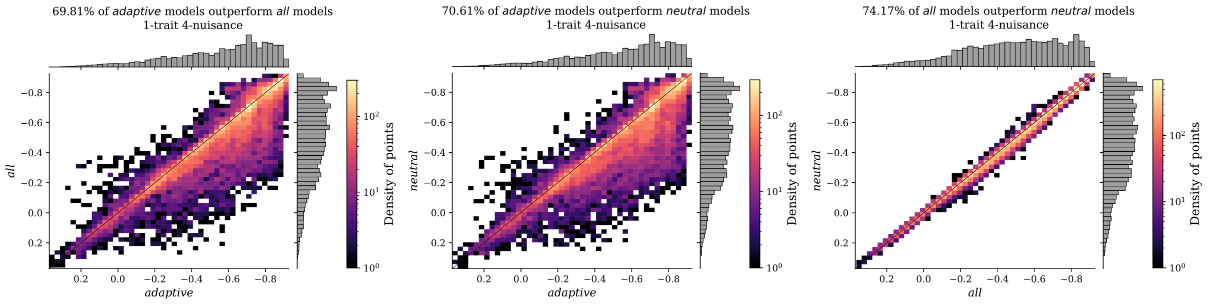

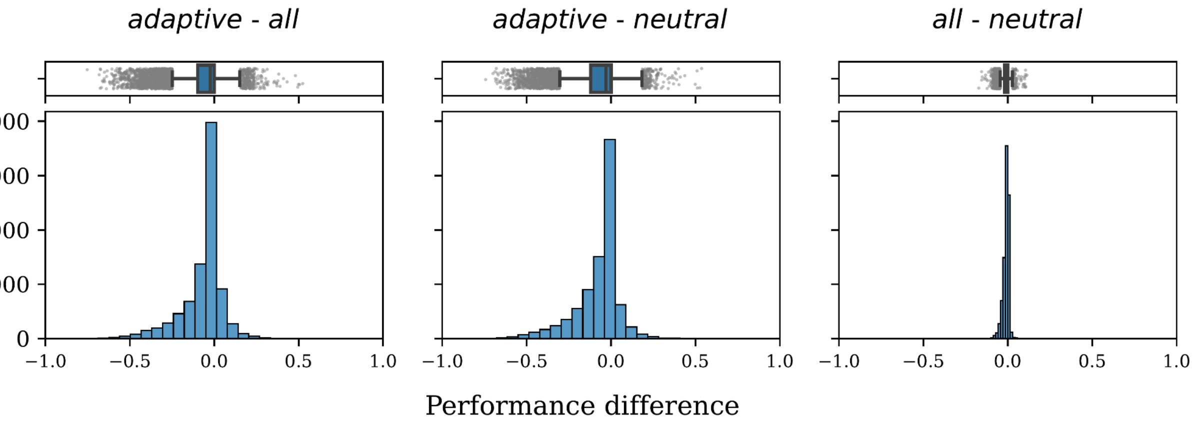

(Fig. S31 continued)

**2-trait 2-nuisance**

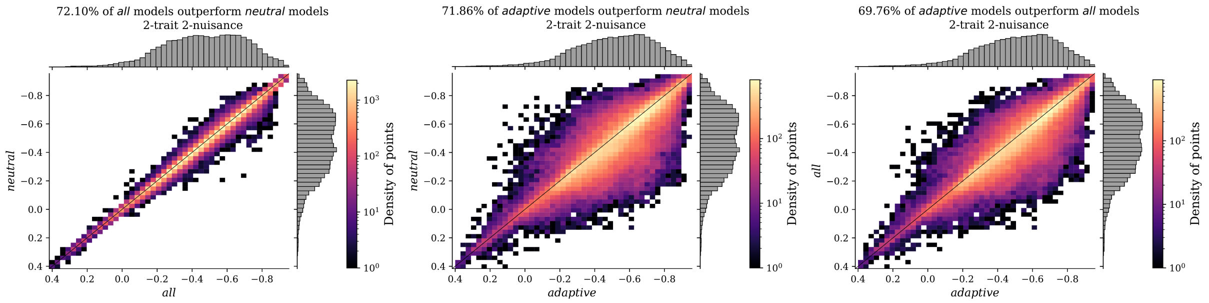

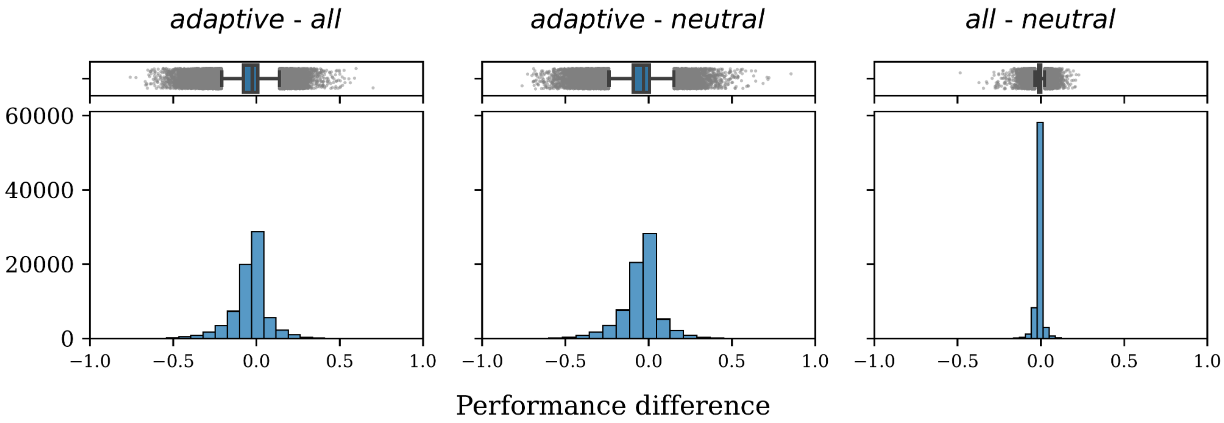

(Fig. S31 continued)

**2-trait 3-nuisance**

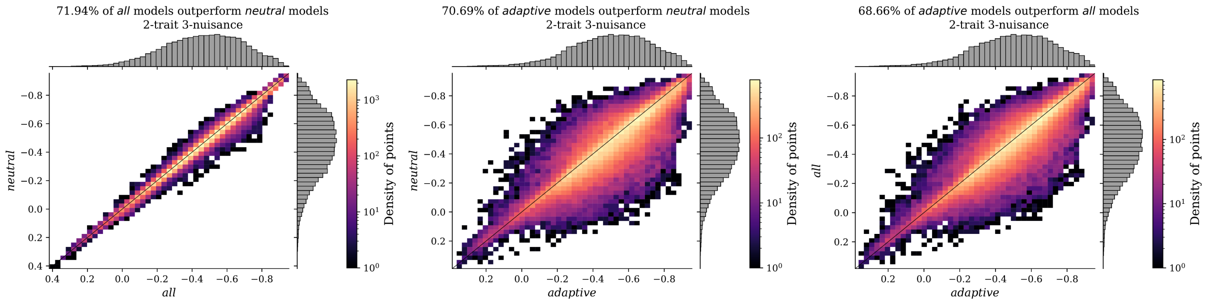

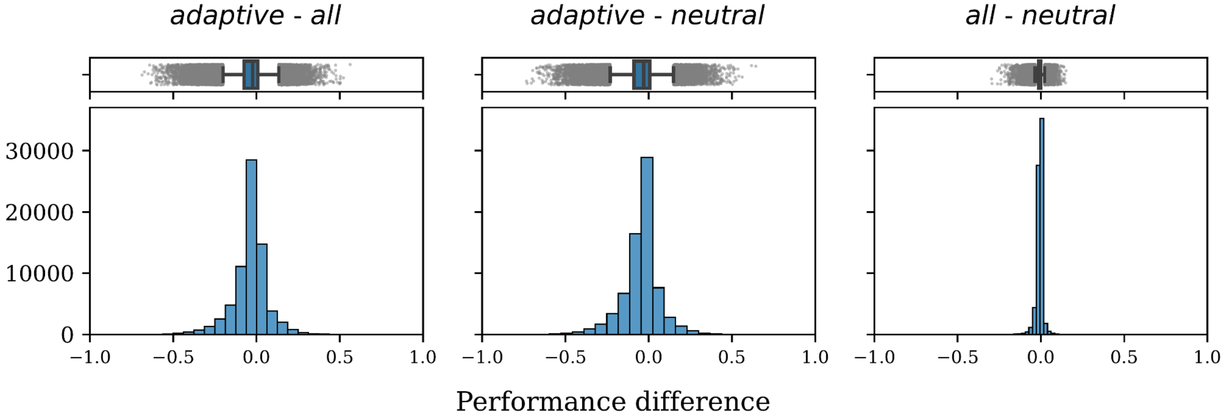

**Fig S32** Pairwise comparison of performance differences between marker sets for *Nuisance Environment* scenarios. The first row for each nuisance level (*N-trait N-nuisance*) are scatterplots of pairwise comparisons of performance between marker sets (histograms in each margin) from both 1- and 2-trait models where density of points is indicated by color in legend (note color scale is different for each figure to accentuate patterns in data). The second row for each nuisance level are histograms for the difference in performance between marker sets for a given model. Method-specific figures are not shown except in SC 02.02.06. Data for these figures includes 1- and 2-trait *Nuisance Environment* evaluations. Code to create these figures can be found in SC 02.02.06.

Fig S33 is in Supplemental Note S4

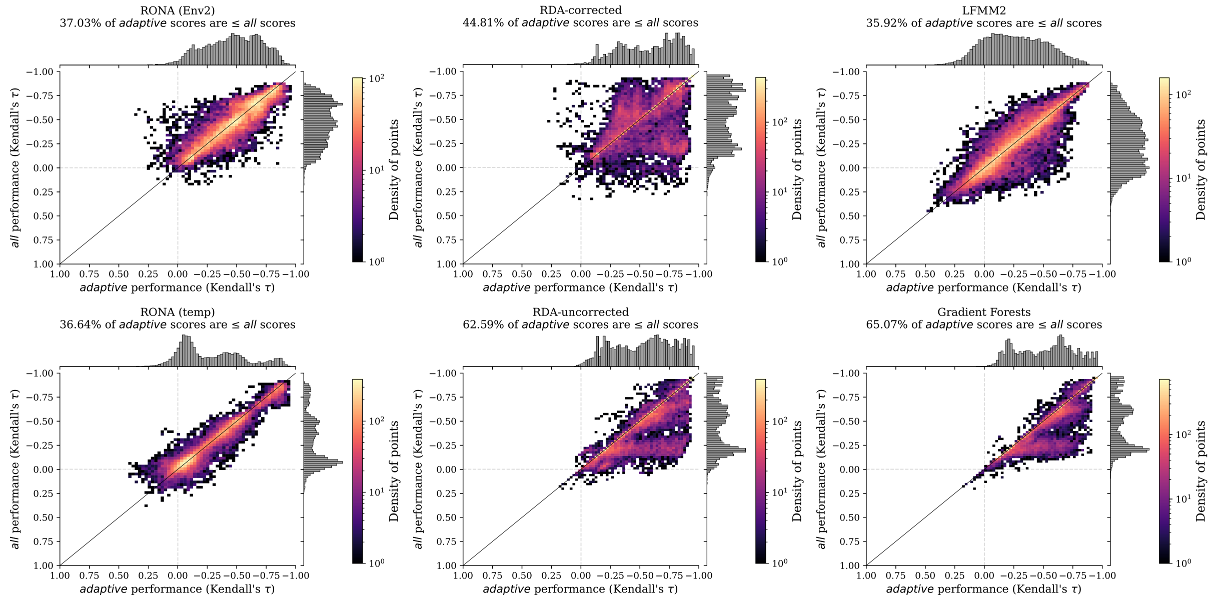

**Fig S34** Pairwise comparison of performance differences between marker sets for *Climate Novelty* scenarios. Shown are scatterplots of pairwise comparisons of performance between marker sets (histograms in each margin) from both 1- and 2-trait models where density of points is indicated by color in legend (note color scale is different for each figure to accentuate patterns in data). Data for these figures includes 1- and 2-trait *Climate Novelty* evaluations. Code to create these figures can be found in SC 02.04.05.
